# Bacteriocin Prediction Through Cross-Validation-Based and Hypergraph-Based Feature Evaluation Approaches

**DOI:** 10.1101/2025.07.12.664537

**Authors:** Suraiya Akhter, John H. Miller

## Abstract

Bacteriocins offer a promising solution to antibiotic resistance, possessing the ability to target a wide range of bacteria with precision. Thus, there is an urgent need for a computational model to predict new bacteriocins and aid in drug development. This work centers on constructing predictive models with XGBoost machine learning algorithm, using physicochemical structural properties and sequence profiles of protein sequences. We employed correlation analyses, cross-validation, and hypergraph-based techniques to select features. Cross-validation feature evaluation (CVFE) partitions the dataset, selects features within each partition, and identifies common features, ensuring representativeness. On the contrary, hypergraph-based feature evaluation (HFE) focuses on minimizing hypergraph cut conductance, leveraging higher-order data relationships to precisely utilize information regarding feature and sample correlations. The XGBoost models were built using the selected features obtained from these two feature evaluation methods. Our HFE-based approach achieved 99.11% accuracy and an AUC of 0.9974 on the test data, overall outperforming the CVFE-based feature evaluation method and yielding results comparable to existing approaches. We also analyzed the feature contributions directly from the best model using SHapley Additive exPlanations (SHAP). Our web application, accessible at https://shiny.tricities.wsu.edu/bacteriocin-prediction/, offers prediction results, probability scores, and SHAP plots using both cross-validation- and hypergraph-based methods, along with previously implemented approaches for feature selection.

## Introduction

Antibiotics have been extensively utilized in animal husbandry and food processing to combat pathogens and extend shelf life, yet their usage has precipitated concerning consequences including bacterial resistance and the dissemination of antibiotic resistance genes [1-3]. This has prompted a shift towards natural alternatives, driven further by consumer preferences for additive-free, healthy foods [4]. Consequently, there is a growing interest in exploring alternative antibacterial agents to control foodborne pathogens. Bacteriocins, proteins synthesized by bacteria, have emerged as promising antimicrobial agents due to their effectiveness against various microbes, including genetically similar strains [2]. They offer advantages such as high efficacy, low toxicity, and minimal residue production, making them attractive substitutes for conventional antibiotics [5-7]. Despite their potential, identifying and characterizing bacteriocins pose challenges. While conventional techniques such as screening assays, chromatography, and mass spectrometry are employed for this task [8-10], they frequently demonstrate to be lengthy, laborious, and expensive, possibly disregarding the breadth and originality of bacteriocins within intricate microbial populations [11].

To overcome constraints in identifying bacteriocins, computational methods like BLASTP are employed to predict them by recognizing patterns or motifs in bacteriocin sequences [12]. Additional tools include BACTIBASE, which integrates microbial data from PubMed alongside protein examination utilities for the characterization of bacteriocins [13], and BAGEL, which categorizes bacteriocin sequences based on homology [14], each upholding repositories of validated bacteriocin sequences. Despite their utility, these methods rely on sequence alignment and may struggle with novel or highly diverse bacteriocins. antiSMASH, a different tool, utilizes hidden Markov models in conjunction with BLAST searches across a database of bacteriocin biosynthetic gene clusters to uncover potential clusters. [15]. While some tools like BOA aim to address bacteriocin diversity, they still rely on homology-based identification, limiting their ability to detect highly dissimilar bacteriocins lacking conserved context genes [16].

Machine learning algorithms offer a distinct approach from traditional sequence matching methods in bacteriocin prediction, enabling the detection of patterns and characteristics beyond mere similarity to known sequences. These algorithms can analyze physicochemical properties, sequence profiles, and secondary structures to uncover novel bacteriocins with significant dissimilarities. Recent advancements include the utilization of *k*-mer features and word embedding techniques, such as those presented by [17, 18]. Furthermore, the RMSCNN technique, based on convolutional neural networks (CNNs), has been developed for bacteriocin prediction [19]. Despite these advancements, existing methods overlook the importance of analyzing both primary and secondary peptide structures and lack feature evaluation mechanisms. To address these limitations, recently we unveiled BaPreS, a machine learning-based software tool, and BPAG, a machine learning-based web application, which employ support vector machines (SVMs) and feature evaluation techniques such as *t*-tests, genetic algorithm and alternating decision tree to precisely identify novel bacteriocins [20, 21].

In this work, our objective was to create predictive models utilizing the XGBoost machine learning algorithm [22], incorporating physicochemical features, sequence profiles and structural properties of bacteriocin and non-bacteriocin sequences chosen through correlation analysis followed by sophisticated techniques like cross-validation [23] and hypergraph-based methods [24]. Subsequently, we evaluated the prediction performance of the models and examined the influence or significance of these selected features within the models using SHapley Additive exPlanations (SHAP) [25]. We integrated cross-validated feature evaluation (CVFE) and hypergraph-based feature evaluation (HFE) methods, along with corresponding SHAP analyses, into our existing web-based tool available at https://shiny.tricities.wsu.edu/bacteriocin-prediction/ [21]. This enhancement allows users to choose CVFE and HFE techniques, along with selecting features based on alternating decision tree, genetic algorithm, linear SVC, or *t*-test methods to obtain precise prediction results and probability scores by cross-checking with different feature evaluation methods. Our web tool also automatically generates the necessary features from user-supplied protein sequences. Users have the capability to concurrently assess numerous sequences and incorporate fresh data, thereby boosting the capacity of the predictive models embedded within the web application.

## Materials and methods

The complete process of our approach is illustrated in Figure 1. It involves gathering protein datasets for both bacteriocin (positive) and non-bacteriocins (negative), creating possible candidate features, employing feature assessment methods, applying the Pearson correlation coefficient, followed by CVFE and hypergraph-based feature evaluation (HFE) techniques to remove the least significant and irrelevant features. Subsequently, selected feature sets are used to construct machine learning models for assessing prediction performance.

**Figure 1:**
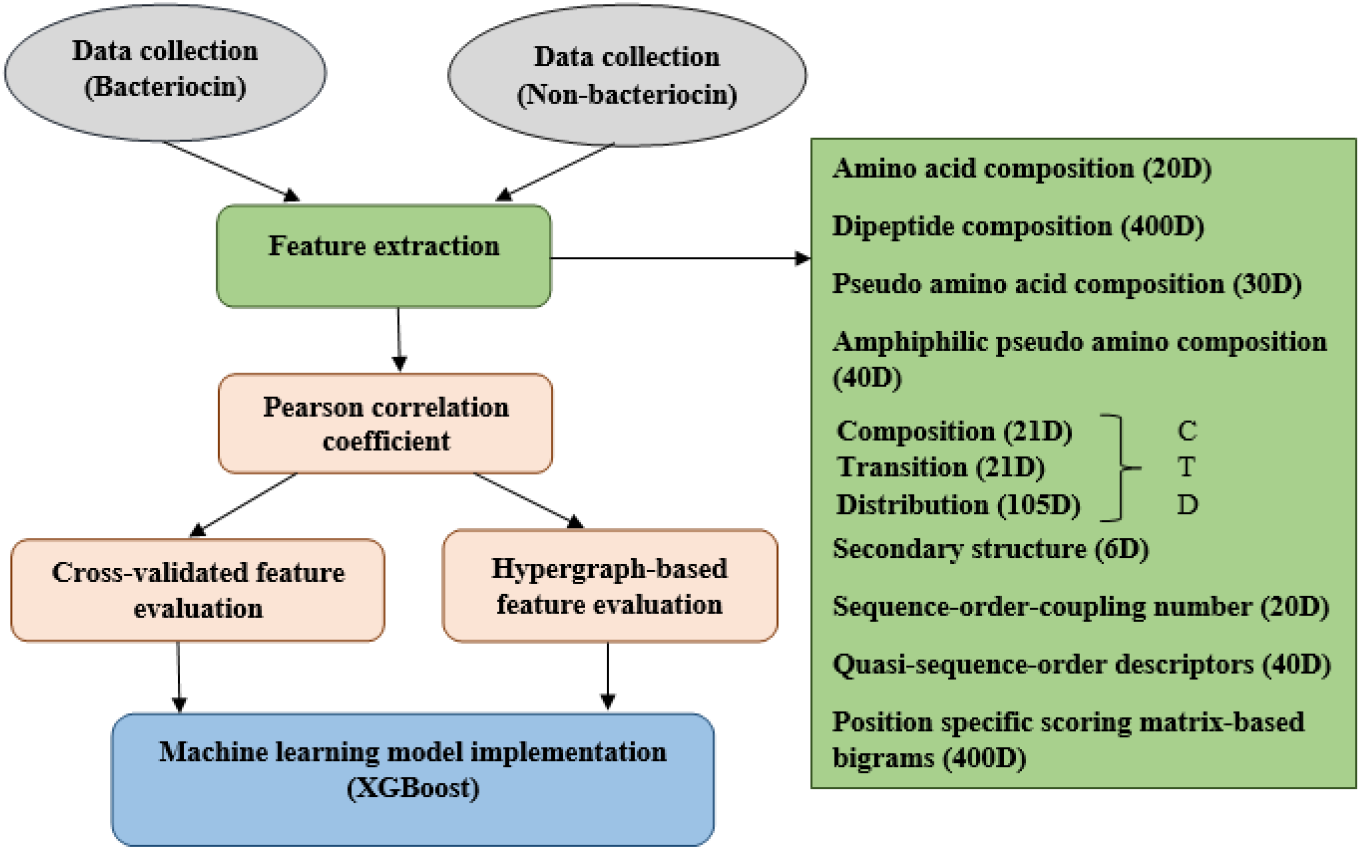
Illustrating the procedure for bacteriocin detection.

### Dataset

The datasets utilized in this work align with those employed in the creation of our previously released software and web applications [20, 21]. The ultimate data set included 283 different positive sequences and 497 distinct negative sequences. To address the challenge of imbalanced data, random sampling was employed, resulting in a reduction of negative sequences to 283, thereby attaining equilibrium between these two groups of the sequences. For training purposes, 80% of the data set was assigned, leaving the remaining 20% for testing. The **Supplementary Material** contains both the training and testing datasets.

### Features

Constructing robust predictive machine learning models depends greatly on identifying and extracting potential attributes. In our investigation, we developed various sets of features to encompass diverse aspects of protein sequences. These comprised a 20-dimensional amino acid composition (AAC), a 400-dimensional dipeptide composition (DC), a 30-dimensional pseudo amino acid composition (PseAAC), and a 40-dimensional amphiphilic pseudo amino acid composition (APseAAC). Moreover, we applied the composition/transition/distribution (CTD) model [26], yielding 147-dimensional attribute sets that account for a range of physicochemical characteristics of amino acids. Furthermore, we devised 6-dimensional feature sets to illustrate the secondary structure (SS) nuances such as α-helix, β-strand, and γ-coil within individual protein sequences. We employed the amino acid distance matrix to generate 20-dimensional sequence-order-coupling number (SOCN) feature sets and 40-dimensional quasi-sequence-order (QSO) feature sets for every sequence [27]. We also employed the position-specific scoring matrix (PSSM) to extract features reflecting evolutionary trends, resulting in a 400-dimensional attribute set for each sequence by calculating transition scores between adjacent amino acids derived from the PSSM [28, 29]. Detailed elucidations of these feature sets are provided in our developed BaPreS software tool [20] and BPAGS web application [21]. In total, 1,103 features were considered as candidates.

### Feature assessment

To ensure the efficacy of a predictive model, it’s essential to eliminate irrelevant features before building the model. Our initial step involved examining the correlation among these features using the Pearson correlation coefficient, employing a methodology akin to that utilized in our earlier implemented BaPreS [20] and BPAGS [21] tools.To prevent any inadvertent sharing of information between training and testing datasets, our focus remained solely on features within the training data. When two features exhibited a high correlation (≥ 0.9), one was retained while the other was discarded. This choice led to a reduction in the number of features from 1,103 to 602. The **Supplementary Material**, specifically Supplementary Table S1, provides a comprehensive list of reduced features utilized in our study. These features are represented by abbreviations such as “aac,” “dipep,” “pseudo,” “amphipseudo,” “comp,” “tran,” “dist,” “ss,” “qso,” and “pssm,” which correspond to amino acid composition (AAC), dipeptide composition (DC), pseudo amino acid composition (PseAAC), amphiphilic pseudo amino acid composition (APseAAC), composition (CTD), transition (CTD), distribution (CTD), secondary structure (SS), quasi-sequence order (QSO), and position-specific scoring matrix (PSSM)-based features, respectively. These reduced features have been comprehensively clarified in our previously published work, BPAGS [21].

We opted to employ the cross-validated feature evaluation (CVFE) technique [23] along with a hypergraph based method [24] to further distill features obtained via Pearson’s correlation analysis. The overall workflow in the CVFE algorithm is depicted in Fig. 2. At first, we randomly partitioned the dataset into *c* subsets and then we applied the XGBoost algorithm to identify the most important features for each subset. We tuned the hyperparameters of the XGBoost model via grid search. We performed feature selection for each subset of the data and then created an intersected feature set that comprises the common features found for all subsets. We repeat this procedure *e* times to generate *e* intersected feature sets. Finally, if any unique feature is found in at least (*p* × 100)% of all the intersected feature sets, we added this in our final selected feature set. The number of features in the feature subsets found through the CVFE method (with different combinations of *c, e*, and *p* values) is mentioned in Table 1. The compilation of significant features is available in supplementary tables S2-S5 provided in the **Supplementary Material**.

**Table 1:**
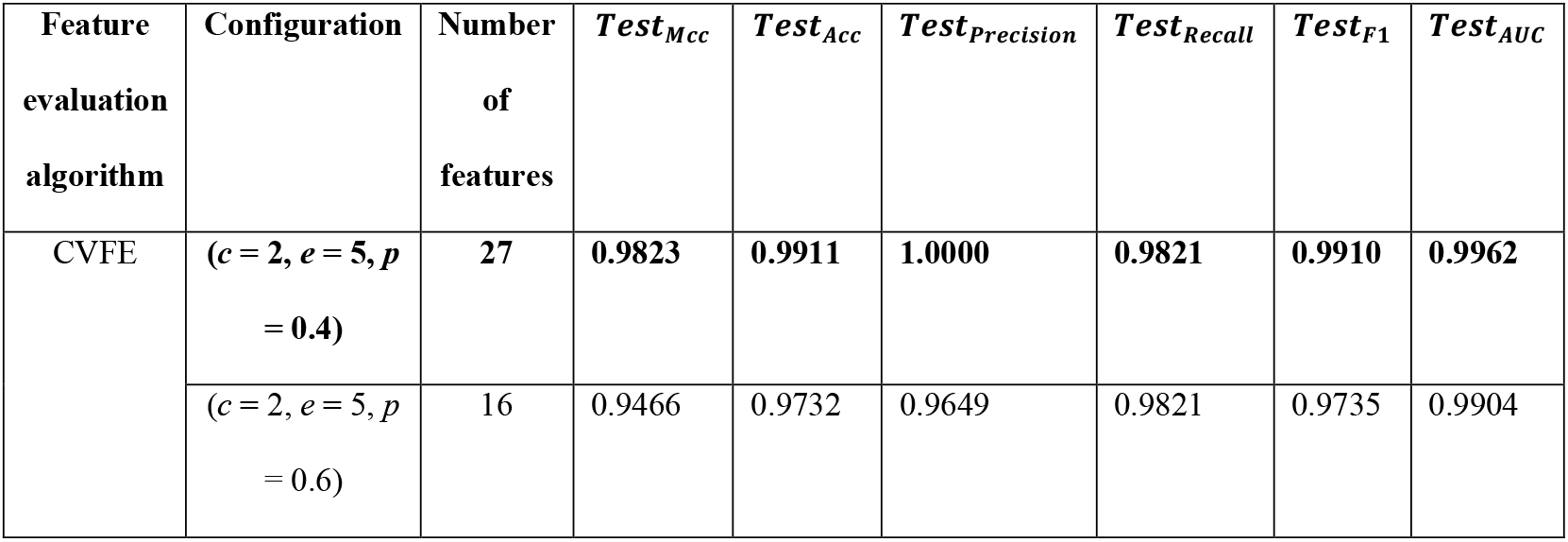
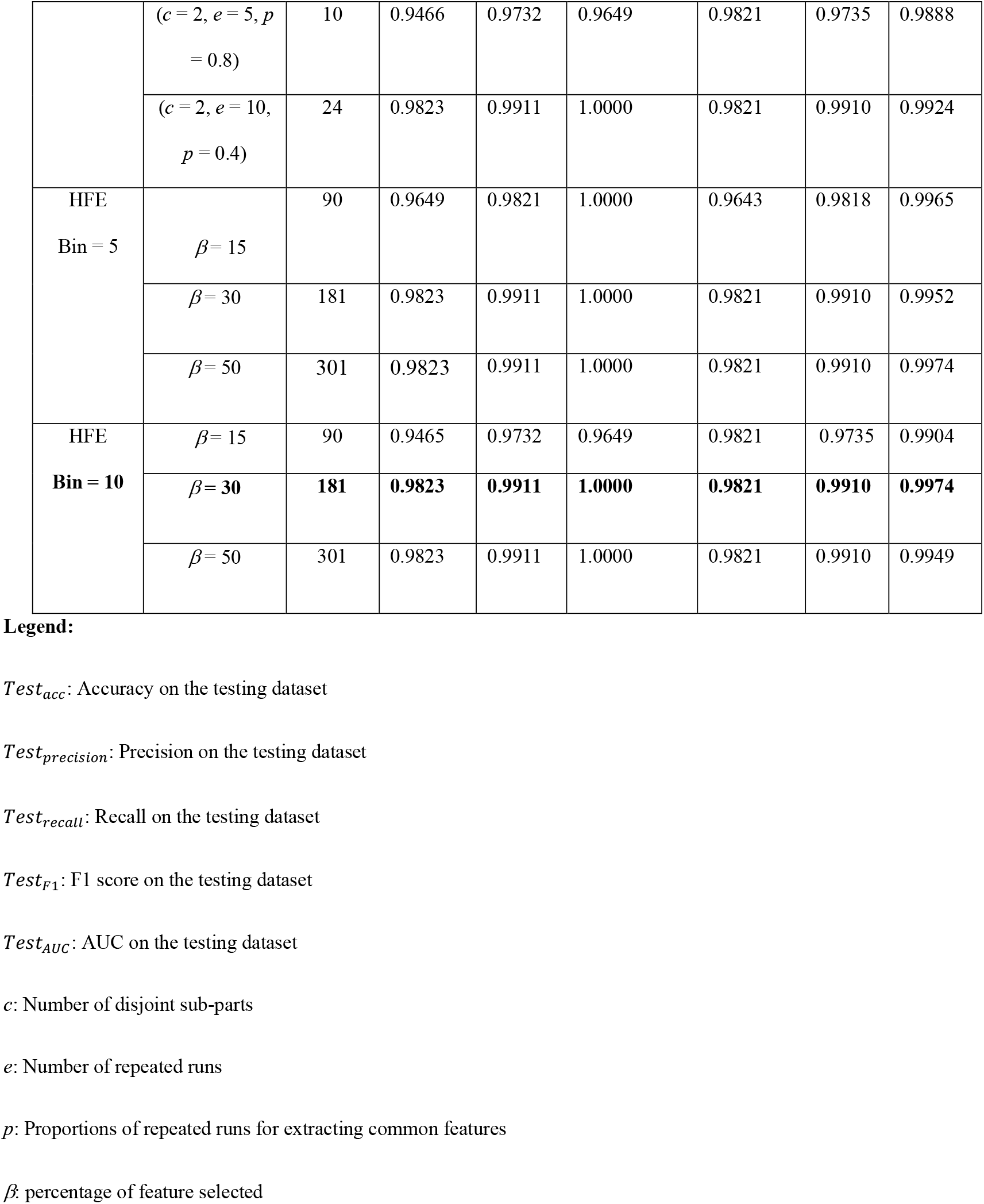
Number of features, MCC, accuracy, precision, recall, F1 and AUC values for testing data for various feature subsets. The most effective models for CVFE and HFE feature sets are highlighted in bold.

| Feature evaluation algorithm | Configuration | Number of features | $Test_{Mcc}$ | $Test_{Acc}$ | $Test_{precision}$ | $Test_{Recall}$ | $Test_{F1}$ | $Test_{AUC}$ |
| --- | --- | --- | --- | --- | --- | --- | --- | --- |
| CVFE | $(c = 2, e = 5, p = 0.4)$ | 27 | **0.9823** | **0.9911** | **1.0000** | **0.9821** | **0.9910** | **0.9962** |
| | $(c = 2, e = 5, p = 0.6)$ | 16 | 0.9466 | 0.9732 | 0.9649 | 0.9821 | 0.9735 | 0.9904 |
| Bacteriocin Detection Using Predictive Models |  |  |  |  |  |  |  |  |
| | $(c = 2, e = 5, p = 0.8)$ | 10 | 0.9466 | 0.9732 | 0.9649 | 0.9821 | 0.9735 | 0.9888 |
| | $(c = 2, e = 10, p = 0.4)$ | 24 | 0.9823 | 0.9911 | 1.0000 | 0.9821 | 0.9910 | 0.9924 |
| HFE<br>Bin = 5 | $\beta = 15$ | 90 | 0.9649 | 0.9821 | 1.0000 | 0.9643 | 0.9818 | 0.9965 |
| | $\beta = 30$ | 181 | 0.9823 | 0.9911 | 1.0000 | 0.9821 | 0.9910 | 0.9952 |
| | $\beta = 50$ | 301 | 0.9823 | 0.9911 | 1.0000 | 0.9821 | 0.9910 | 0.9974 |
| HFE<br>Bin = 10 | $\beta = 15$ | 90 | 0.9465 | 0.9732 | 0.9649 | 0.9821 | 0.9735 | 0.9904 |
| | $\beta = 30$ | 181 | 0.9823 | 0.9911 | 1.0000 | 0.9821 | 0.9910 | 0.9974 |
| | $\beta = 50$ | 301 | 0.9823 | 0.9911 | 1.0000 | 0.9821 | 0.9910 | 0.9949 |

**Figure 2:**
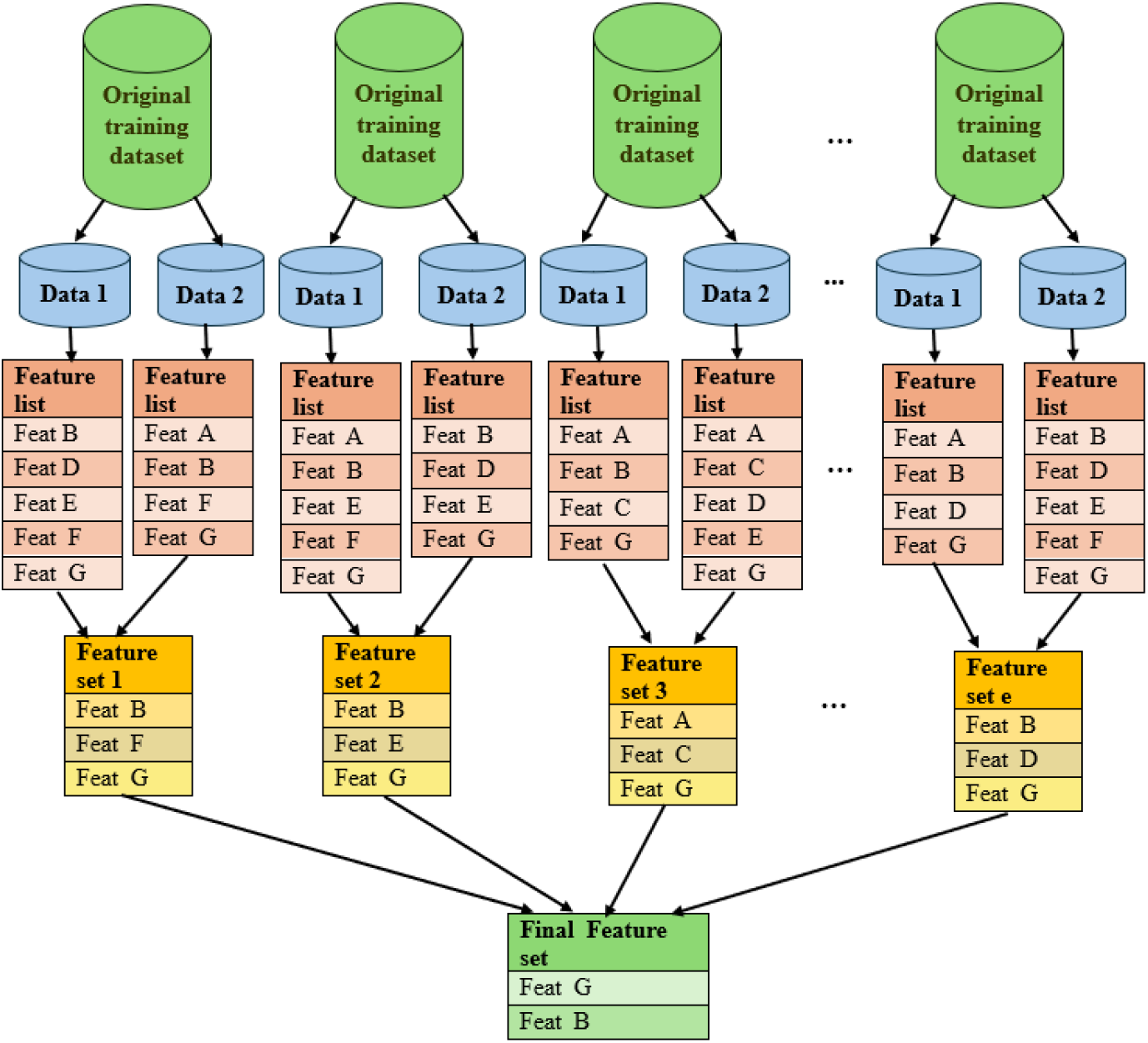
Depicting the procedure of selecting features using the CVFE approach.

The method of hypergraph-based feature evaluation (HFE) entails constructing a hypergraph using features identified through Pearson’s correlation analysis. Unlike a conventional graph denoted by G(*V, E*) with vertices (*V*) and edges (*E*), a hypergraph extends this notion by allowing edges to connect multiple vertices. Formally, a hypergraph is depicted as G(*V, E*), where *V* represents the vertices set and *E* indicates the edges, also known as hyperedges, which are subsets of *V*. Figure 3 illustrates both a basic graph and a hypergraph, and the hypergraph representation of our entire training dataset is depicted in Supplementary Figure S1 (**Supplementary Material**). In the HFE approach, we discretized the values for each feature into some uniform bins, and the significance/importance rates of the features were estimated based on the higher-order relationships in the training dataset. The relevant features are selected in the HFE by assigning ratings to hypergraph edges that correspond to feature values. The hypergraph-based importance rating is measured based on the hypergraph cut conductance minimization theory [24], which results in obtaining a vector of hypergraph-based ratings for all features. The values in the vector are sorted to get *z* top-ranked features. If the number of features is *m* then *z* = *β* × *m*, where the parameter value *β* is used to determine the percentage of features we want to select after HFE analysis. In our case, we considered bin = 5 and bin = 10, and the numbers of features selected for different β and bins are mentioned in Table 1. The lists of the reduced features using HFE for bin = 5 and bin = 10 are given in the Supplementary Tables S6 and S7 (**Supplementary Material)**.

**Figure 3:**
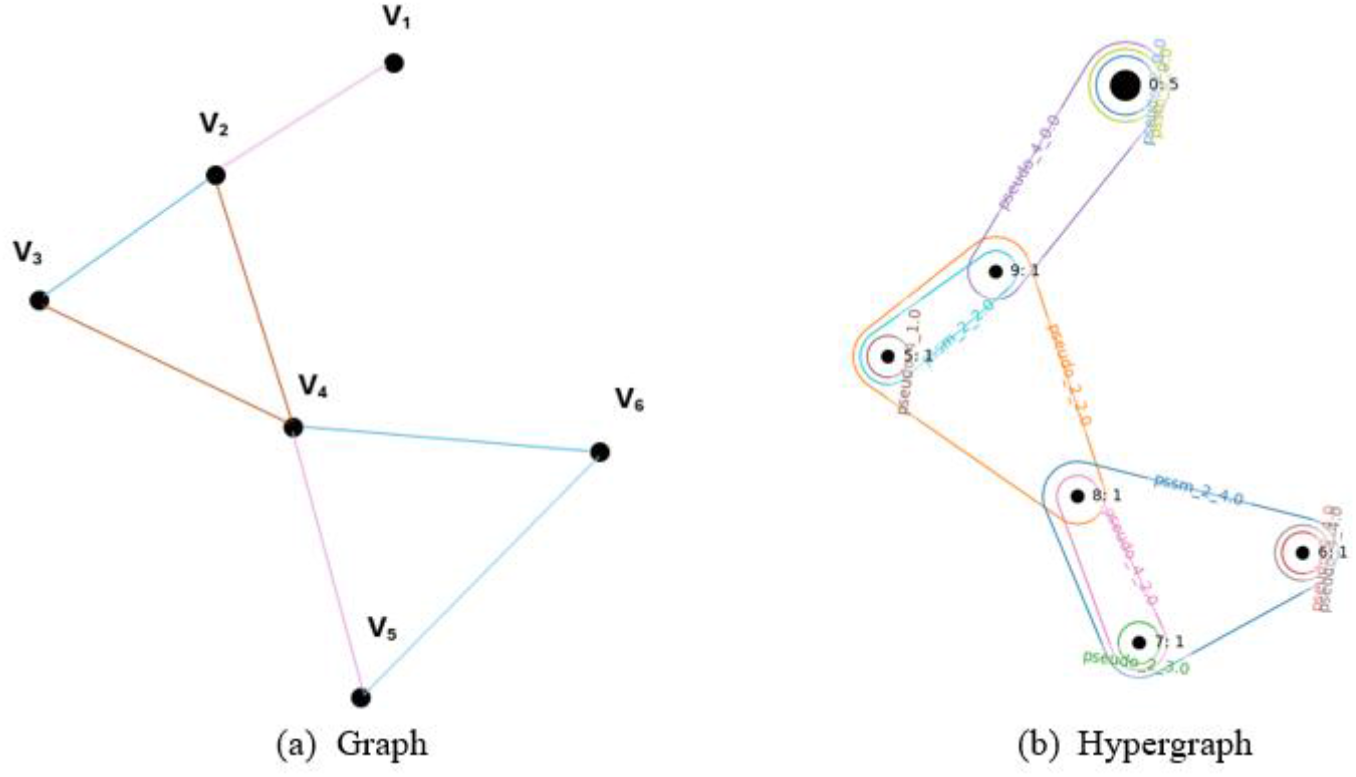
Illustrating the difference between a graph and a hypergraph.

### Web application

The CVFE and HFE methods were integrated into the web application along with previously implemented feature evaluation approaches [21], as illustrated in Figure 4. This machine learning-based web tool autonomously generates features for user-provided sequences, yielding classification and probability results. Details on data upload, binary classification, and probability estimation are outlined within the web application, and a user can download it from the web application. Users can download necessary files and augment training data with new protein sequences, enhancing prediction accuracy. The updated web application now provides users the facility of downloading the SHAP plot to inspect the impact of the top 10 features on the prediction outcomes. The web server can be publicly accessible at https://shiny.tricities.wsu.edu/bacteriocin-prediction/.

**Figure 4:**
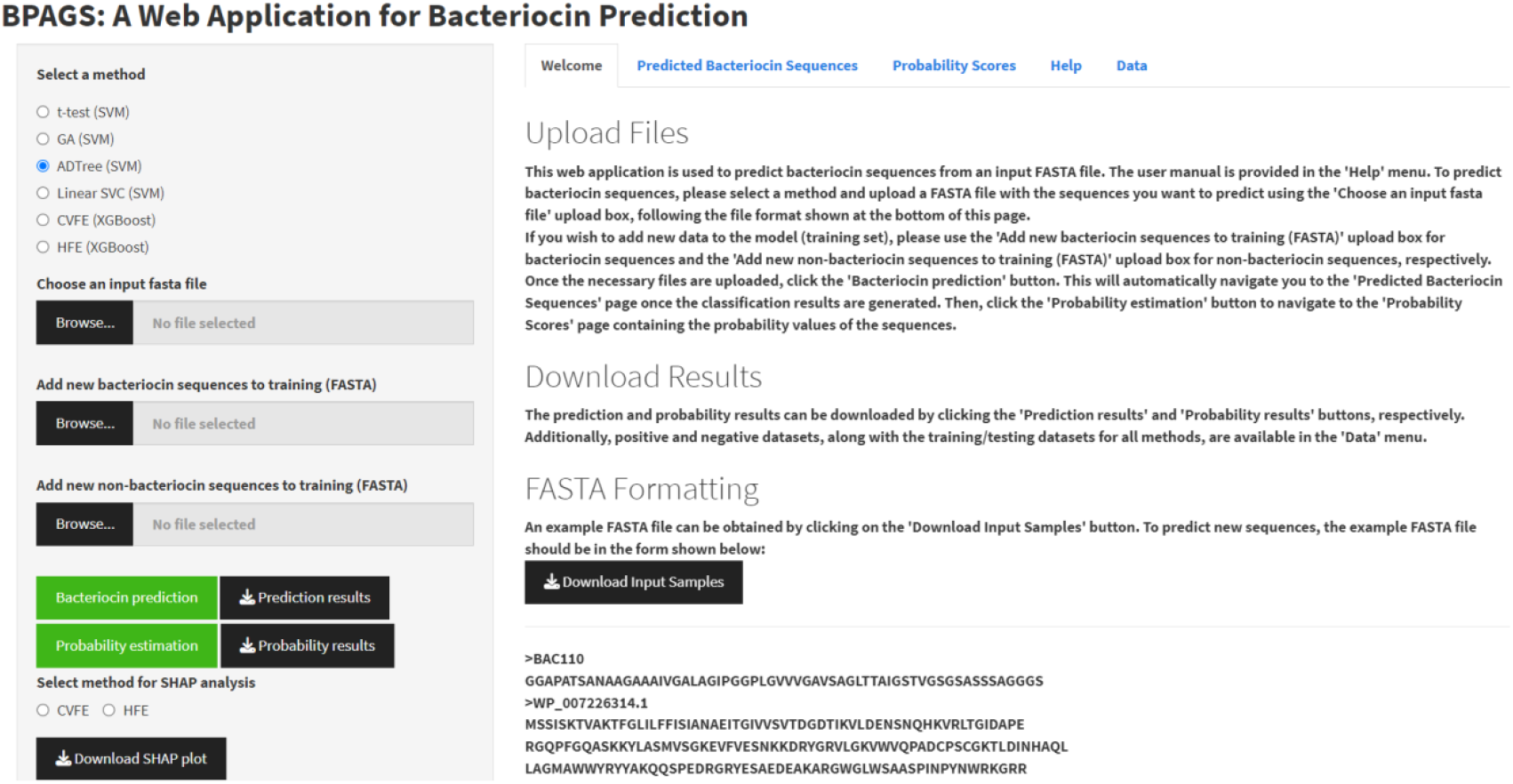
The web application for bacteriocin prediction.

### Code and data availability

The complete set of experimental data and accompanying scripts is available at https://github.com/suraiya14/cvfe_hfe.

## Results and discussion

After downsizing the feature collection through two distinct feature assessment techniques, we constructed separate predictive models utilizing the chosen features through the XGBoost [22] machine learning technique. By employing the SHAP (Shapley Additive Explanations) approach [25], we evaluated the significance of features and their contributions to the XGBoost models. SHAP values quantify the additional influence of each feature on the forecasts generated by the machine learning model.

### Model performance

We developed an XGBoost model by training it on various feature subsets derived from CVFE and HFE analyses alongside the training dataset. Evaluation of the model’s predictive accuracy on the testing dataset was conducted using equations 1 to 5, where TP, TN, FP, and FN denote true positives, true negatives, false positives, and false negatives, respectively. To assess classifier effectiveness, Accuracy assesses the proportion of accurately categorized instances compared to the total instances within the dataset. We utilized the Matthews correlation coefficient (MCC), a metric ranging from -1 to +1, with higher values indicating superior prediction capabilities.

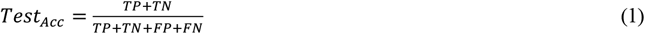

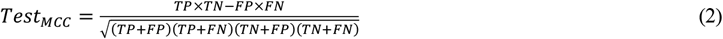

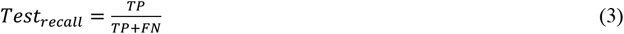

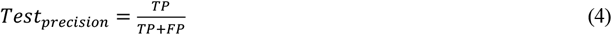

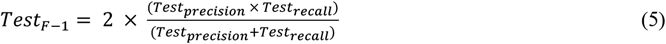

We also calculated recall and precision. Recall measures the proportion of correctly identified true positive instances, while precision evaluates the fraction of accurate positive predictions. The F1 score, a metric that accounts for both precision and recall, computes their harmonic mean, thus presenting a well-rounded assessment of the model’s efficacy. Furthermore, we determined the Area Under the Curve (AUC) to appraise the efficacy of binary classification models. A greater AUC value signifies better performance, where 1 denotes perfection and 0.5 indicates random chance.

Table 1 elaborates on the evaluation of XGBoost models across CVFE and HFE-based feature subsets, while Supplementary Figure S2 (**Supplementary Material**) illustrates the confusion matrices for all reduced feature sets. Overall, we obtained better prediction results for the HFE (bin = 10, *β* = 30) feature set compared to the CVFE (*c* = 2, *e* = 5, *p* = 0.4) feature set, with our best model able to identify 55 protein sequences.

### Selected features in the HFE Approach

The top-performing machine-learning outcome was achieved through implementing the XGBoost model, utilizing HFE method with bin = 10 and *β* = 30 parameters. The HFE selected 181 features from a pool of 701, predominantly focusing on distribution, amphiphilic pseudo amino acid, and dipeptide composition. For detailed insight into the chosen features on the training data, please refer to Supplementary Tables S7 provided in the **Supplementary material**.

### Feature contribution analysis

Figure 5 displays the ranking of the top 10 features based on their mean SHAP values for predicting bacteriocin presence for the best XGBoost model with hypergraph (bin = 10, *β* = 30) reduced feature sets. Each point on the plot represents a protein sequence, with overlapping points visualized through jittering to indicate their frequency. The *x*-axis indicates the influence of features on the model’s output, which is either a prediction of 1 (bacteriocin) or 0 (non-bacteriocin). The *y*-axis shows the mean |SHAP| values of the features. The color bar at the bottom of the figure represents the value of features where yellow and purple correspond to low and high values, respectively. The features in the plot are in ascending of importance determined based on the mean |SHAP| of the features. The detail descript of the feature can be found in [21, 27]. For instance, the dist_93 feature (i.e., group 1 (buried) of the CTD model [26] for solvent accessibility) is identified as the most important feature, and the next important feature is aac_5 (i.e., amino acid composition for Cysteine (C)). The solvent accessibility influences various aspects of bacteriocins, including their interaction with target bacterial membranes and their stability in the extracellular environment, and cysteine residues have been found as important properties in *bactofencin* bacteriocin [30, 31].

**Figure 5:**
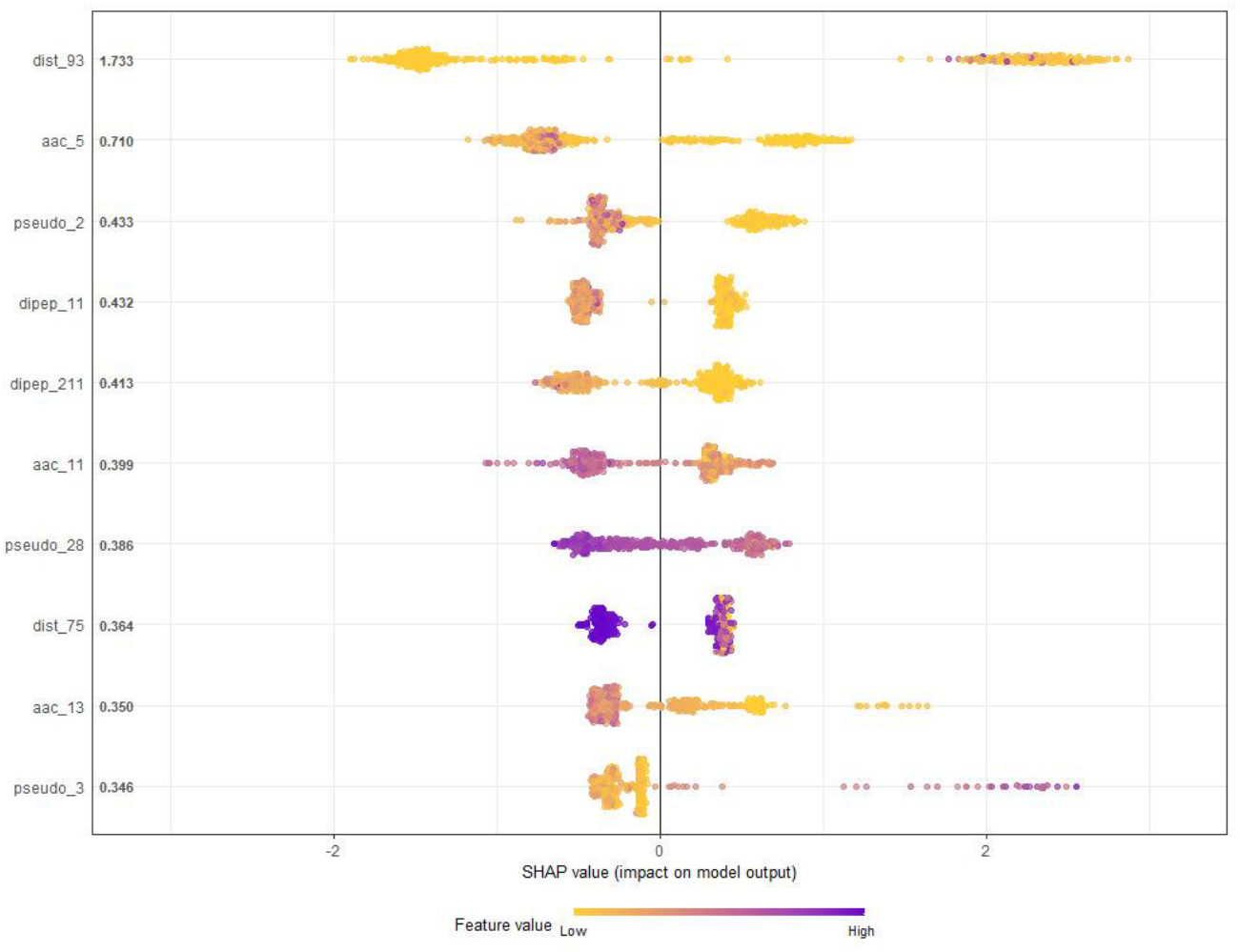
Feature importance (SHAP values) obtained from the XGBoost model for predicting bacteriocins.

A higher SHAP value indicates a higher likelihood of bacteriocin protein sequence. In Fig. 5, the SHAP values tend to be higher for increased dist_93 feature of a protein sequence, thereby showing a higher likelihood of being that sequence a bacteriocin. Also, the higher the aac_5 feature value is, the higher the chance of a sequence being classified as non-bacteriocin due to low SHAP value. We can analyze the impact/contribution of other features in the plot in a similar way. The feature importance plot based on the SHAP value for all features can be found in Supplementary Figure S3 (**Supplementary Material**).

### Performance comparison

The prediction performance of the XGBoost models built using CVFS-reduced feature sets and HFE-reduced feature sets was compared with the deep learning method RMSCNN [19], as well as our previously introduced BaPreS and BPAGS tools [20, 21]. RMSCNN, developed specifically for the detection of marine microbial bacteriocins through Convolutional Neural Networks (CNN), transforms protein sequences into numeric formats to facilitate feature acquisition and prediction. BaPreS and BPAGS automate feature generation and selection through correlation and *t*-test analyses, alternating decision tree, and genetic algorithm, employing Support Vector Machines (SVM) for prediction. Our XGBoost models demonstrated superior performance compared to RMSCNN and BaPreS, while providing comparable prediction results to BPAGS (refer to Table 2).

**Table 2:** Evaluation of the efficacy of models/tools in predicting bacteriocins.

| Method/tool | *Test<sub>acc</sub>* | *Test<sub>precision</sub>* | *Test<sub>recall</sub>* | *Test<sub>F1</sub>* | *Test<sub>AUC</sub>* |
| --- | --- | --- | --- | --- | --- |
| RMSCNN | 0.9375 | 0.9623 | 0.9107 | 0.9358 | 0.9818 |
| BaPreS | 0.9554 | 0.9636 | 0.9464 | 0.9550 | 0.9879 |
| BPAGS (ADTree) | 0.9911 | 0.9825 | 1.0000 | 0.9912 | 0.9984 |
| BPAGS (GA) | 0.9643 | 0.9643 | 0.9643 | 0.9643 | 0.9968 |
| BPAGS (Linear SVC) | 0.9732 | 0.9818 | 0.9643 | 0.9730 | 0.9990 |
| CVFE ($c = 2, e = 5, p = 0.4$) | 0.9911 | 1.0000 | 0.9821 | 0.9910 | 0.9962 |
| HFE (bin = 5, $\beta = 30$) | 0.9911 | 1.0000 | 0.9821 | 0.9910 | 0.9974 |

## Conclusion

The pursuit of new bacteriocins is essential for advancing the creation of fresh antibiotic treatments to counter the escalating threat of antibiotic resistance. This study introduces predictive models aimed at identifying novel bacteriocins. Our approach involves extracting diverse features from primary and secondary attributes of protein sequences, alongside sequence profiles. These features are then subjected to analysis using Pearson correlation coefficient, followed by CVFS and HFE feature evaluations. Subsequently, we employed the XGBoost machine learning algorithm using the selected feature sets. Our findings indicate that XGBoost demonstrates superior predictive capabilities, particularly when using the HFE-reduced feature set. Notably, the distribution feature of protein sequences emerged as the most significant, contributing to the optimal model. The efficacy of our most proficient model was evaluated by comparing it with both deep learning methods and tools we had previously created. The findings indicate that XGBoost demonstrated comparable or improved performance in comparison.

Our web application integrates both CVFS and HFE for feature evaluation, incorporating all necessary programs to automatically generate an optimal feature set. Users can now utilize CVFS and HFE alongside existing methods to predict bacteriocin presence in unseen testing data. Additionally, they can augment the training data with new bacteriocin and non-bacteriocin sequences and perform SHAP analyses, enhancing the predictive capability of the web tool. Presently, our model proficiently discerns singular bacteriocin protein sequences, and our goal is to improve its capability to identify protein clusters such as tailocins (bacteriocins resembling phage tails). Future improvements include the incorporation of granular molecular component-based features and protein-protein interaction networks-based features and the development of a robust feature selection algorithm to enhance prediction accuracy. We will maintain and update our machine learning-based web application as more bacteriocin sequences become available.

## Supporting information

The supplementary material for this article is provided herewith the manuscript. (DOCX)

## Authors’ contributions

SA: Conceptualization, data collection, formal analysis, software implementation, validation, visualization, and writing manuscript. JHM: Conceptualization, supervision, reviewing analyses, and editing the manuscript. All authors read and approved the final manuscript.

## Competing interests

The authors declare that they have no competing interests.

## Notes

### Competing Interest Statement

The authors have declared no competing interest.

## References

1. Braïek OB, Morandi S, Cremonesi P, Smaoui S, Hani K, Ghrairi T. Safety, potential biotechnological and probiotic properties of bacteriocinogenic Enterococcus lactis strains isolated from raw shrimps. Microbial pathogenesis. 2018;117:109–17.

2. Meade E, Slattery MA, Garvey M. Bacteriocins, potent antimicrobial peptides and the fight against multi drug resistant species: resistance is futile? Antibiotics. 2020;9(1):32.

3. Ren S, Yuan X, Liu F, Fang F, Iqbal HM, Zahran SA, et al. Bacteriocin from Lacticaseibacillus rhamnosus sp. A5: isolation, purification, characterization, and antibacterial evaluation for sustainable food processing. Sustainability. 2022;14(15):9571.

4. Khodaei M, Sh SN. Isolation and molecular identification of bacteriocin-producing enterococci with broad antibacterial activity from traditional dairy products in Kerman province of Iran. Korean Journal for Food Science of Animal Resources. 2018;38(1):172.

5. Riley MA, Wertz JE. Bacteriocins: evolution, ecology, and application. Annual Reviews in Microbiology. 2002;56(1):117–37.

6. Hamid MN, Friedberg I, editors. Bacteriocin detection with distributed biological sequence representation. ICML Computational Biology workshop; 2017.

7. Fields FR, Freed SD, Carothers KE, Hamid MN, Hammers DE, Ross JN, et al. Novel antimicrobial peptide discovery using machine learning and biophysical selection of minimal bacteriocin domains. Drug development research. 2020;81(1):43–51.

8. Zendo T, Nakayama J, Fujita K, Sonomoto K. Bacteriocin detection by liquid chromatography/mass spectrometry for rapid identification. Journal of Applied Microbiology. 2008;104(2):499–507.

9. Zhang J, Yang Y, Yang H, Bu Y, Yi H, Zhang L, et al. Purification and partial characterization of bacteriocin Lac-B23, a novel bacteriocin production by Lactobacillus plantarum J23, isolated from Chinese traditional fermented milk. Frontiers in microbiology. 2018;9:2165.

10. Desiderato CK, Sachsenmaier S, Ovchinnikov KV, Stohr J, Jacksch S, Desef DN, et al. Identification of potential probiotics producing bacteriocins active against Listeria monocytogenes by a combination of screening tools. International Journal of Molecular Sciences. 2021;22(16):8615.

11. Perez RH, Zendo T, Sonomoto K. Novel bacteriocins from lactic acid bacteria (LAB): various structures and applications. Microbial cell factories. 2014;13(1):1–13.

12. Boratyn GM, Camacho C, Cooper PS, Coulouris G, Fong A, Ma N, et al. BLAST: a more efficient report with usability improvements. Nucleic acids research. 2013;41(W1):W29–W33.

13. Hammami R, Zouhir A, Le Lay C, Ben Hamida J, Fliss I. BACTIBASE second release: a database and tool platform for bacteriocin characterization. Bmc Microbiology. 2010;10(1):1–5.

14. Van Heel AJ, de Jong A, Montalban-Lopez M, Kok J, Kuipers OP. BAGEL3: automated identification of genes encoding bacteriocins and (non-) bactericidal posttranslationally modified peptides. Nucleic acids research. 2013;41(W1):W448–W53.

15. Weber T, Blin K, Duddela S, Krug D, Kim HU, Bruccoleri R, et al. antiSMASH 3.0—a comprehensive resource for the genome mining of biosynthetic gene clusters. Nucleic acids research. 2015;43(W1):W237–W43.

16. Morton JT, Freed SD, Lee SW, Friedberg I. A large scale prediction of bacteriocin gene blocks suggests a wide functional spectrum for bacteriocins. BMC bioinformatics. 2015;16(1):1–9.

17. Mikolov T, Chen K, Corrado G, Dean J. Efficient estimation of word representations in vector space. arXiv preprint arXiv:13013781. 2013.

18. Hamid M-N, Friedberg I. Identifying antimicrobial peptides using word embedding with deep recurrent neural networks. Bioinformatics. 2019;35(12):2009–16.

19. Cui Z, Chen Z-H, Zhang Q-H, Gribova V, Filaretov VF, Huang D-S. Rmscnn: A random multi-scale convolutional neural network for marine microbial bacteriocins identification. IEEE/ACM Transactions on Computational Biology and Bioinformatics. 2021;19(6):3663–72.

20. Akhter S, Miller JH. BaPreS: a software tool for predicting bacteriocins using an optimal set of features. BMC bioinformatics. 2023;24(1):313.

21. Akhter S, Miller JH. BPAGS: a web application for bacteriocin prediction via feature evaluation using alternating decision tree, genetic algorithm, and linear support vector classifier. Frontiers in Bioinformatics. 2023;3.

22. Chen T, Guestrin C, editors. Xgboost: A scalable tree boosting system. Proceedings of the 22nd acm sigkdd international conference on knowledge discovery and data mining; 2016.

23. Yang M-R, Wu Y-W. A Cross-Validated Feature Selection (CVFS) approach for extracting the most parsimonious feature sets and discovering potential antimicrobial resistance (AMR) biomarkers. Computational and Structural Biotechnology Journal. 2023;21:769–79.

24. Misiorek P, Janowski S. Hypergraph-based importance assessment for binary classification data. Knowledge and Information Systems. 2023;65(4):1657–83.

25. Lundberg SM, Erion G, Chen H, DeGrave A, Prutkin JM, Nair B, et al. From local explanations to global understanding with explainable AI for trees. Nature machine intelligence. 2020;2(1):56–67.

26. Dubchak I, Muchnik I, Holbrook SR, Kim S-H. Prediction of protein folding class using global description of amino acid sequence. Proceedings of the National Academy of Sciences. 1995;92(19):8700–4.

27. Xiao N, Cao D-S, Zhu M-F, Xu Q-S. protr/ProtrWeb: R package and web server for generating various numerical representation schemes of protein sequences. Bioinformatics. 2015;31(11):1857–9.

28. Saini H, Raicar G, Lal SP, Dehzangi A, Imoto S, Sharma A. Protein fold recognition using genetic algorithm optimized voting scheme and profile bigram. Journal of Software. 2016;11(8):756–67.

29. Mohammadi A, Zahiri J, Mohammadi S, Khodarahmi M, Arab SS. PSSMCOOL: a comprehensive R package for generating evolutionary-based descriptors of protein sequences from PSSM profiles. Biology Methods and Protocols. 2022;7(1):bpac008.

30. Benítez-Chao DF, León-Buitimea A, Lerma-Escalera JA, Morones-Ramírez JR. Bacteriocins: An overview of antimicrobial, toxicity, and biosafety assessment by in vivo models. Frontiers in Microbiology. 2021;12:630695.

31. O’Connor PM, O’Shea EF, Cotter PD, Hill C, Ross RP. The potency of the broad spectrum bacteriocin, bactofencin A, against staphylococci is highly dependent on primary structure, N-terminal charge and disulphide formation. Scientific reports. 2018;8(1):11833.

