## Supplementary Material for "Bacteriocin Prediction Through Cross-Validation-Based and Hypergraph-Based Feature Evaluation Approaches"

### 1 Supplementary Data

#### 1.1 Training dataset

**Bacteriocin**

**----------------------------------------**

>BAC010

CRQSCSFGPFTFVCDGNTK

>BAC011

CANSCSYGPLTWSCDGNTK

>BAC014

CTFTLPGGGGVCTLTSECIC

>BAC015

GGAGHVPEYFVGIGTPISFYG

>BAC017

IASKFICTPGCAKTGSFNSYCC

>BAC023

GNGVLKTISHECNMNTWQFLFTCC

>BAC024

GNPKVAHCASQIGRSTAWGAVSGA

>BAC025

NRWWQGVVPTVSYECRMNSWQHVFTCC

>BAC030

GKNGVFKTISHECHLNTWAFLATCCS

>BAC033

KGGSGVIHTISHEVIYNSWNFVFTCCS

>BAC034

KGGSGVIHTISHECNMNSWQFVFTCCS

>BAC042

KGKGFWSWASKATSWLTGPQQPGSPLLKKHR

>BAC043

SASVLKTSIKVSKKYCKGVTLTCGCNITGGK

>BAC045

WKSESLCTPGCVTGALQTCFLQTLTCNCKISK

>BAC047

ITSISLCTPGCKTGALMGCNMKTATCHCSIHVSK

>BAC048

TAGPAIRASVKQCQKTLKATRLFTVSCKGKNGCK

>BAC055

SDCNINSNTAADVILCFNQVGSCALCSPTLVGGPVP

>BAC056

KYYGNGLSCSKKGCTVNWGQAFSCGVNRVATAGHGK

>BAC057

STPVLASVAVSMELLPTASVLYSDVAGCFKYSAKHHC

>BAC058

KYYGNGVHCTKSGCSVNWGEAFSAGVHRLANGGNGFW

>BAC060

TSYGNGVHCNKSKCWIDVSELETYKAGTVSNPKDILW

>BAC062

GLGKAQCAALWLQCASGGTIGCGGGAVACQNYRQFCR

>BAC065

ARSYGNGVYCNNKKCWVNRGEATQSIIGGMISGWASGLAGM

>BAC069

AISYGNGVYCNKEKCWVNKAENKQAITGIVIGGWASSLAGMGH

>BAC070

TKYYGNGVYCNSKKCWVDWGTAQGCIDVVIGQLGGGIPGKGKC

>BAC071

KNYGNGVHCTKKGCSVDWGYAWTNIANNSVMNGLTGGNAGWHN

>BAC072

VGIGGGGGGGGGGSCGGQGGGCGGCSNGCSGGNGGSGGSGSHI

>BAC074

QINWGSVVGHCIGGAIIGGAFSGGAAAGVGCLVGSGKAIINGL

>BAC080

MGAIAKLVAKFGWPIVKKYYKQIMQFIGEGWAINKIIDWIKKHI

>BAC081

KYYGNGVSCNKNGCTVDWSKAIGIIGNNAAANLTTGGAAGWNKG

>BAC085

ANCSCSTASDYCPILTFCTTGTACSYTPTGCGTGWVYCACNGNFY

>BAC087

SLQYVMSAGPYTWYKDTRTGKTICKQTIDTASYTFGVMAEGWGKTFH

>BAC089

MDKFEKISTSNLEKISGGDLTTKLWSSWGYYLGKKARWNLKHPYVQF

>BAC090

ILFSYLLFYVLKENSKREDKYQNIIEELTELLPKIKEDVEDIKEKLNK

>BAC091

VNYGNGVSCSKTKCSVNWGQAFQERYTAGINSFVSGVASGAGSIGRRP

>BAC095

NRWTNAYSAALGCAVPGVKYGKKLGGVWGAVIGGVGGAAVCGLAGYVRKG

>BAC097

MQKPEIISADLGLCAVNEFVALAAIPGGAATFAVCQMPNLDEIVSNAAYV

>BAC098

MKLPVQQVYSVYGGKDLPKGHSHSTMPFLSKLQFLTKIYLLDIHTQPFFI

>BAC100

DQMSDGVNYGKGSSLSKGGAKCGLGIVGGLATIPSGPLGWLAGAAGVINSCMK

>BAC102

KLTFIQSTAAGDLYYNTNTHKYVYQQTQNAFGAAANTIVNGWMGGAAGGFGLHH

>BAC103

MNNLNKFSTLGKSSLSQIEGGSVPTSVYTLGIKILWSAYKHRKTIEKSFNKGFYH

>BAC118

ATYYGNGVYCNKQKCWVDWSRARSEIIDRGVKAYVNGFTKVLGGIGGR

>BAC119

GETDPNTQLLNDLGNNMAWGAALGAPGGLGSAALGAAGGALQTVGQGLIDHGPVNVPIPV

LIGPSWNGSGSGYNSATSSSGSGS

>BAC122

ASIIKTTIKVSKAVCKTLTCICTGSCSNCK

>BAC123

VPGGCTYTRSNRDVIGTCKTGSGQFRIRLDCNNAPDKTSVWAKPKVMVSVHCLVGQPRSI

SFETK

>BAC124

KYYGNGVHCGKKTCYVDWGQATASIGKIIVNGWTQHGPWAHR

>BAC125

DIAPPGPNGDPKSVQIDDKYTGAEMYGEGDFRVGLGTDLTMYPPVYRESLGNGSGGWEFD

FTVCGSTACRFVDSNGDVKEDDKAKEMWWQEINFNDINQDLYSRNDSDWVGSTPADTQPE

FDYTDFALARDGVTLALTALNPAMGSLALGATYFLSDMVNWIASQHEDDSSLKRKWDYDG

LSGPLYADSSTYLLARDEMTSNSYESFTIDNIAVAFPEFPVRTKYYVTFTAPDDPSTQSI

STLEEEGIYRVPATE

>BAC126

MARPIADLIHFNSTTVTASGDVYYGPGGGTGIGPIARPIEHGLDSSTENGWQEFESYADV

GVDPRRYVPLQVKEKRREIELQFRDAEKKLEASVQAELDKADAALGPAKNLAPLDVINRS

LTIVGNALQQKNQKLLLNQKKITSLGAKNFLTRTAEEIGEQAVREGNINGPEAYMRFLDR

EMEGLTAAYNVKLFTEAISSLQIRMNTLTAAKASIEAAAANKAREQAAAEAKRKAEEQAR

QQAAIRAANTYAMPANGSVVATAAGRGLIQVAQGAASLAQAISDAIAVLGRVLASAPSVM

AVGFASLTYSSRTAEQWQDQTPDSVRYALGMDAAKLGLPPSVNLNAVAKASGTVDLPMRL

TNEARGNTTTLSVVSTDGVSVPKAVPVRMAAYNATTGLYEVTVPSTTAEAPPLILTWTPA

SPPGNQNPSSTTPVVPKPVPVYEGATLTPVKATPETYPGVITLPEDLIIGFPADSGIKPI

YVMFRDPRDVPGAATGKGQPVSGNWLGAASQGEGAPIPSQIADKLRGKTFKNWRDFREQF

WIAVANDPELSKQFNPGSLAVMRDGGAPYVRESEQAGGRIKIEIHHKVRVADGGGVYNMG

NLVAVTPKRHIEIHKGGK

>BAC128

GWWNSWGKCVAGTIGGAGTGGLGGAAAGSAVPVIGTGIGGAIGGVSGGLTGAATFC

>BAC131

MDKVTDNSPDVESTESTEGSFPTVGVDTGDTITATLATGTENVGGGGGAFGGASESSAAI

HATAKWSTAQLKKHQAEQAARAAAAEAALAKAKSQRDALTQRLKDIVNDALRANAARSPS

VTDLAHANNMAMQAEAERLRLAKAEQKAREEAEAAEKALREAERQRDEIARQQAETAHLL

AMAEAAEAEKNRQDSLDEEHRAVEVAEKKLAEAKAELAKAESDVQSKQAIVSRVAGELEN

AQKSVDVKVTGFPGW

>BAC132

MGSNGADNAHNNAFGGGKNPGIGNTSGAGSNGSASSNRGNSNGWSWSNKPHKNDGFHSDG

SYHITFHGDNNSKPKPGGNSGNRGNNGDGASAKVGEITITPDNSKPGRYISSNPEYSLLA

KLIDAESIKGTEVYTFHTRKGQYVKVTVPDSNIDKMRVDYVNWKGPKYNNKLVKRFVSQF

LLFRKEEKEKNEKEALLKASELVSGMGDKLGEYLGVKYKNVAKEVANDIKNFHGRNIRSY

NEAMASLNKVLANPK

>BAC134

MKHLNETTNVRILSQFDMDTGYQAVVQKGNVGSKYVYGLQLRKGATTILRGYRGSKINNP

ILELSGQAGGHTQTWEFAGDRKDINGEERAGQWFIGVKPSKIEGSKIIWAKQIARVDLRN

QMGPHYSNTDFPRLSYLNRAGSNPFAGNKMTHAEAAVSPDYTKFLIATVENNCIGHFTIY

NLDTINEKLDEKGNSEDVNLETVKYEDSFIIDNLYGDDNNSIVNSIQGYDLDNDGNIYIS

SQKAPDFDGSYYAHH

>BAC135

METLTVHAPSPSTNLPSYGNGAFSLSAPHVPGAGPLLVQVVYSFFQSPNMCLQALTQLED

YIKKHGASNPLTLQIISTNIGYFCNADRNLVLHPGISVYDAYHFAKPAPSQYDYRSMNMK

QMSGNVTTPIVALAHYLWGNGAERSVNIANIGLKISPMKINQIKDIIKSGVVGTFPVSTK

FTHATGDYNVITGAYLGNITLKTEGTLTISANGSWTYNGVVRSYDDKYDFNASTHRGIIG

ESLTRLGAMFSGKEY

>BAC136

MNKTHKMATLVIAAILAAGMTAPTAYADSPGNTRITASEQSVLTQILGHKPTQTEYNRYV

ETYGSVPTEADINAYIEASESEGSSSQTAAHDDSTSPGTSTEIYTQAAPARFSMFFLSGT

WITRSGVVSLSLKPRKGGIGNEGDERTWKTVYDKFHNAGQWTRYKNNGVDASMKKQYMCH

FKYGMVKTPWNLEPHKKAADVSPVKCN

>BAC141

MSWLNFLKYIAKYGKKAVSAAWKYKGKVLEWLNVGPTLEWVWQKLKKIAGL

>BAC142

ATYYGNGLYCNKEKCWVDWNQAKGEIGKIIVNGWVNHGPWAPRR

>BAC143

MAKEFGIPAAVAGTVLNVVEAGGWVTTIVSILTAVGSGGLSLLAAAGRESIKAYLKKEIK

KKGKRAVIAW

>BAC148

LVAYGIAQGTAEKVVSLINAGLTVGSIISILGGVTVGLSGVFTAVKAAIAKQGIKKAIQL

>BAC150

KTVNYGNGLYCNQKKCWVNWSETATTIVNNSIMNGLTGGNAGWHSGGRA

>BAC153

TTKNYGNGVCNSVNWCQCGNVWASCNLATGCAAWLCKLA

>BAC154

DIGGSRQGCVA

>BAC157

MMNATENQIFVETVSDQELEMLIGGAGRGWIKTLTKDCPNVISSICAGTIITACKNCA

>BAC160

MQTIKELNTMELQEIIGGENDHRMPYELNRPNNLSKGGAKCAAGILGAGLGAVGGGPGGF

ISAGISAVLGCM

>BAC161

AYPGNGVHCGKYSCTVDKQTAIGNIGNNAA

>BAC162

LAGYTGIASGTAKKVVDAIDKGAAAFVIISIISTVISAGALGAVSASADFIILTVKNYIS

RNLKAQAVIW

>BAC164

MSLLALVAGTLGVSQSIATTVVSIVLTGSTLISIILGITAILSGGVDAILEIGWSAFVAT

VKKIVAERGKAAAIAW

>BAC166

WFYQGMNIAIYANIGGVANIIGYTEAAVATLLGAVVAVAPVVP

>BAC170

MAGFLKVVQILAKYGSKAVQWAWANKGKILDWINAGQAIDWVVEKIKQILGIK

>BAC172

MSDPVRITNPGAESLGYDSDGHEIMAVDIYVNPPRVDVFHGTPPAWSSFGNKTIWGGNEW

VDDSPTRSDIEKRDKEITAYKNTLSAQQKENENKRTEAGKRLSAAIAAREKDENTLKTLR

AGNADAADITRQEFRLLQAELREYGFRTEIAGYDALRLHTESRMLFADADSLRISPREAR

SLIEQAEKRQKDAQNADKKAADMLAEYERRKGILDTRLSELEKNGGAALAVLDAQQARLL

GQQTRNDRAISEARNKLSSVTESLNTARNALTRAEQQLTQQKNTPDGKTIVSPEKFPGRS

STNHSIVVSGDPRFAGTIKITTSAVIDNRANLNYLLSHSGLDYKRNILNDRNPVVTEDVE

GDKKIYNAEVAEWDKLRQRLLDARNKITSAESAVNSARNNLSARTNEQKHANDALNALLK

EKENIRNQLSGINQKIAEEKRKQDELKATKDAINFTTEFLKSVSEKYGAKAEQLAREMAG

QAKGKKIRNVEEALKTYEKYRADINKKINAKDRAAIAAALESVKLSDISSNLNRFSRGLG

YAGKFTSLADWITEFGKAVRTENWRPLFVKTETIIAGNAATALVALVFSILTGSALGIIG

YGLLMAVTGALIDESLVEKANKFWGI

>BAC173

MSDPVRITNPGAESLGYDSDGHEIMAVDIYVNPPRVDVFHGTPPAWSSFGNKTIWGGNEW

VDDSPTRSDIEKRDKEITAYKNTLSAQQKENENKRTEAGKRLSAAIAAREKDENTLKTLR

AGNADAADITRQEFRLLQAELREYGFRTEIAGYDALRLHTESRMLFADADSLRISPREAR

SLIEQAEKRQKDAQNADKKAADMLAEYERRKGILDTRLSELEKNGGAALAVLDAQQARLL

GQQTRNDRAISEARNKLSSVTESLKTARNALTRAEQQLTQQKNTPDGKTIVSPEKFPGRS

STNHSIVVSGDPRFAGTIKITTSAVIDNRANLNYLLTHSGLDYKRNILNDRNPVVTEDVE

GDKKIYNAEVAEWDKLRQRLLDARNKITSAESAINSARNNVSARTNEQKHANDALNALLK

EKENIRSQLADINQKIAEEKRKRDEINMVKDAIKLTSDFYRTIYDEFGKQASELAKELAS

VSQGKQIKSVDDALNAFDKFRNNLNKKYNIQDRMAISKALEAINQVHMAENFKLFSKAFG

FTGKVIERYDVAVELQKAVKTDNWRPFFVKLESLAAGRAASAVTAWAFSVMLGTPVGILG

FAIIMAAVSALVNDKFIEQVNKLIGI

>BAC174

NRWYCNSAAGGVGGAAVCGLAGYVGEAKENIAGEVRKGWGMAGGFTHNKACKSFPGSGWA

SG

>BAC176

AGDPLADPNSQIVRQIMSNAAWGPPLVPERFRGMAVGAAGGVTQTVLQGAAAHMPVNVPI

PKVPMGPSWNGSKG

>BAC178

KGLGKLIGIDWLLGQAKDAVKQYKKDYKRWH

>BAC181

AVPAVRKTNETLD

>BAC182

GNGVVLTLTHECNLATWTKKLKCC

>BAC185

KPAWCWYTLAMCGAGYDSGTCDYMYSHCFGVKHSSGGGGSYHC

>BAC186

NETNNFAETQKEITTNSEATLTNEDYTKLTSEVKTIYTNLIQYDQTKNKFYVDEDKTEQY

YNYDDESIKGVYLMKDSLNDELNNNNSSNYSEIINQKISEIDYVLQGNDINNLIPSNTRV

KRSADFSWIQRCLEEAWGYAISLVTLKGIINLFKAGKFEAAAAKLASATAGRIAGMAALF

AFVATCGATTVS

>BAC187

PNWTKIGKCAGSIAWAIGSGLFGGAKLIKIKKYIAELGGLQKAAKLLVGATTWEEKLHAG

GYALINLAAELTGVAGIQANCF

>BAC189

SNDSLWYGVGQFMGKQANCITNHPVKHMIIPGYCLSKILG

>BAC190

IAPIIVAGLGYLVKDAWDHSDQIISGFKKGWNGGRRK

>BAC191

LIDHLGAPRWAVDTILGAIAVGNLASWVLALVPGPGWAVKAGLATAAAIVKHQGKAAAAA

W

>BAC192

LVATGMAAGVAKTIVNAVSAGMDIATALSLFSGAFTAAGGIMALIKKYAQKKLWKQLIAA

>BAC193

PNDGDTMTVSGGGGWVSNDDRKGGNDRDNGKGGSAVDFSKNPEKQAIVNPYLAIAIPMPV

YPLYGKLGFTINTTAIETELANVRAAINTKLATLSAVIGRSLPVVGRVFGVTAAGMWPSS

TAPSSLDSIYNQAHQQALAQLAAQQGVLNKGYNVTAMPAGFVSSLPVSEIKSLPTAPASL

LAQSVINTELSQRQLALTQPTTNAPVANIPVVKAEKTAMPGVYSAKIIAGEPAFQIKVDN

TKPALAQNPPKVKDDIQVSSFLSSPVADTHHAFIDFGSDHEPVYVSLSKIVTAEEEKKQV

EEAKRREQEWLLRHPITAAERKLTEIRQVISFAQQLKESSVATISEKTKTVAVYQEQVNT

AAKNRDNFYNQNRGLLSAGITGGPGYPIYLALWQTMNNFHQAYFRANNALEQESHVLNLA

RSDLAKAEQLLAENNRLQVETERTLAEEKEIKRNRVNVSTFGTVQTQLSKLLSDFYAVTS

LSQSVPSGALASFSYNPQGMIGSGKIVGKDVDVLFSIPVKDIPGYKSPINLDDLAKKNGS

LDLPIRLAFSDENGERVLRAFKADSLRIPSSVRGVAGSYDKNTGIFSAEIDGVSSRLVLE

NPAFPPTGNVGNTGNTAPDYKALLNTGVDVKPVDKITVTVTPVADPVDIDDYIIWLPTAS

GSGVEPIYVVFNSNPYGGTEKGKYSKRYYNPDKAGGPILELDWKNVKIDHAGVDNVKLHT

GRFKASVENKVMIERLENILNGQITATDTDKRFYTHELRELNRYRNLGIKDGEVPSSIQE

ESAVWNDTHTATLEDYKINEKEQPLYTDAALQAAYEQELKDALGGKHG

>BAC196

TITLSTCAILSKPLGNNGYLCTVTKECMPSCN

>BAC197

TTPATTSSWTCITAGVTVSASLCPTTKCTSRC

>BAC198

ATYTRPLDTGNITTGFNGYPGHVGVDYAVPVGTPVRAVANGTVKFAGNGANHPWMLWMAG

NCVLIQHADGMHTGYAHLSKISVSTDSTVKQGQIIGYTGATGQVTGPHLHFEMLPANPNW

QNGFSGRIDPTGYIANAPVFNGTTPTEPTTPTTNLKIYKVDDLQKINGIWQVRNNILVPT

DFTWVDNGIAADDVIEVTSNGTRTSDQVLQKGGYFVINPNNVKSVGTPMKGSGGLSWAQV

NFTTGGNVWLNTTSKDNLLYGK

>BAC199

KPAWCWYTLAMCGAGYDSGTCDYMYSHCFGIKHHSSGSSSYHC

>BAC200

KNYGNGVYCNKHKCSVDWATFSANIANNSVAMAGLTGGNAGNK

>BAC202

KYYGNGVSCNSHGCSVNWGQAWTCGVNHLANGGHGVC

>BAC203

MLAKIKAMIKKFPNPYTLAAKLTTYEINWYKQQYGRYPWERPVA

>BAC204

ATYYGNGLYCNKQKHYTWVDWNKASREIGKIIVNGWVQH

>BAC205

FVYGNGVTSILVQAQFLVNGQRRFFYTPDK

>BAC206

VNYGNGVSCSKTKCSVNWGIITHQAFRVTSGVASG

>BAC208

TNYGNGVGVPDAIMAGIIKLIFIFNIRQGYNFGKKAT

>BAC209

MFLVNQLGISKSLANTILGAIAVGNLASWLLALVPGPGWATKAALATAETIVKHEGKAAA

IAW

>BAC210

MAAFMKLIQFLATKGQKYVSLAWKHKGTILKWINAGQSFEWIYKQIKKLWA

>BAC211

ACQCPDAISGWTHTDYQCHGLENKMYRHVYAICMNGTQVYCRTEWGSSC

>BAC212

AIKLVQSPNGNFAASFVLDGTKWIFKSKYYDSSKGYWVGIYEVWDRK

>BAC214

VTTSIPCTVMVSAAVCPTLVCSNKCGGRG

>BAC216

GNAACVIGCIGSCVISEGIGSLVGTAFTLG

>BAC218

ATPATPTVAQFVIQGSTICLVC

>BAC219

IGGALGNALNGLGTWANMMNGGGFVNQWQVYANKGKINQYRPY

>BAC222

VTSWSLCTPGCTSPGGGSNCSFCC

>BAC223

MKTILRFVAGYDIASHKKKTGGYPWERGKA

>BAC224

IVWLANKFGVHLTNHLTNSILNAVSNGSSLGSAFAVIAGVTLPGWAVAAVGALGATAA

>BAC225

VFHAYSARGNYYGNCPANWPSCRNNYKSAGGK

>BAC229

LTANLGISSYAAKKVIDIINTGSAVATIIALVTAVVGGGLITAGIVATAKSLIKKYGAKY

AAAW

>WP_061432710.1

MKNPTLLPKLTAPVERPAVTSSDLKQASSVDAAWLNGDNNWSTPFAGVNAAWLNGDNNWS

TPFAGVNAAWLNGDNNWSTPFAADGAE

>CAX48972.1

MASILELQDLEVERASSAADSNASVWECCSTGSWVPFTCC

>YP_003491235.1

MGPVVVFDCMTADFLNDDPNNAELSALEMEELESWGAWDGEATS

>WP_013079675.1

MTKKNATQAPRLVRVGDAHRLTQGAFVGQPEAVNPLGREIQG

>ACR33052.1

MSALAIEKSWKDVDLRDGATSHPAGLGFGELTFEDLREDRTIYAASSGWVCTLTIECGTV

ICAC

>sp|Q09T02.1|MICA_CLAMM

MNDILETETPVMVSPRWDMLLDAGEDTSPSVQTQIDAEFRRVVSPYMSSSGWLCTLTIEC

GTIICACR

>NP_391616.1

MKKAVIVENKGCATCSIGAACLVDGPIPDFEIAGATGLFGLWG

>sp|O87236.1|LANA1_LACLL

MNKNEIETQPVTWLEEVSDQNFDEDVFGACSTNTFSLSDYWGNNGAWCTLTHECMAWCK

>sp|O87237.1|LANA2_LACLL

MKEKNMKKNDTIELQLGKYLEDDMIELAEGDESHGGTTPATPAISILSAYISTNTCPTTK

CTRAC

>sp|O88038.1|LANSB_STRCO

MNLFDLQSMETPKEEAMGDVETGSRASLLLCGDSSLSITTCN

>AAK33966.1

MNNTIKDFDLDLKTNKKDTATPYVGSRYLCTPGSCWKLVCFTTTVK

>YP_444120.1

MRTLTLNELDSVSGGASGRDIAMAIGTLSGQFVAGGIGAAAGGVAGGAIYDYASTHKPNP

AMSPSGLGGTIKQKPEGIPSEAWNYAAGRLCNWSPNNLSDVCL

>AAL73241.1

MSNTQLLEVLGTETFDVQENLFTFDTTDTIVAESNDDPDTRFKSWSFCTPGCAKTGSFNS

YCC

>sp|Q2QBT0.1|LANNU_STRUB

MNNEDFNLDLIKISKENNSGASPRITSKSLCTPGCKTGILMTCPLKTATCGCHFG

>ANP43731.1

MLDVIKNRKKIEEKLELPEILLEEVEEHSAMGGINTWNTTATSTSIIISETFGNKGKVCT

YTVECVNNCRG

>CAA84399.1

MVTKYGRNLGLSKVELFAIWAVLVVALLLATANIYWIADQFGIHLATGTARKLLDAVASG

ASLGTAFAAILGVTLPAWALAAAGALGATAA

>CAA63706.1

MTNAFQALDEVTDAELDAILGGGSGVIPTISHECHMNSFQFVFTCCS

>sp|Q52053|Q52053_9ZZZZ

MLNKENQENYYSNKLELVGPSFEELSLEEMEAIQGSGDVQAETTPACFTIGLGVGALFSA

KFC

>BAD72777.1

MKEIQKAGLQEELSILMDDANNLEQLTAGIGTTVVNSTFSIVLGNKGYICTVTVECMRNC

SK

>ANP43734.1

MLKEEKLEKITGLIPESELEEHLSGESSGAGTPAITTAISAIIAATAQSPCPTSACSKSC

NK

>AAT87775.1

MERRMSFMKNSKDILTNVIEEVSEKELMEVAGGKKGSGWFATITDDCPNSVFVCC

>BAD05046.1

MSTKDFNLDLVSVSKTDSGASTRITSISLCTPGCKTGVLMGCNLKTATCNCSVHVSK

>AAN86036.1

MFLVAGALGVQTAAATTIVNVILNAGTLVTVLGIIASIASGGAGTLMTIGWATFKATVQK

LAKQSMARAIAY

>AAL15569.1

MSKFDDFDLDVVKVSKQDSKITPQWKSESVCTPGCVTGVLQTCFLQTITCNCHISK

>AAL15567.1

MTNMSKFDDFDLDVVKVSKQDSKITPQVLSKSLCTPGCITGPLQTCYLCFPTFAKC

>AAK32702.1

MNKDLNALTNPIDEKELEQILGGGDGVFRTISHECAMNTWMFIFTCCS

>ARW80050.1

MSMTMTLQQAVVDDEFRSVLLADPAAFGLSVESLPGAVERQDHEAIEAFTEAVVASEIYA

CASTCSFGPFTIACDGTTK

>BAB04173.1

MTNLLKEWKMPLERTHNNSNPAGDIFQELEDQDILAGVNGACAWYNISCRLGNKGAYCTL

TVECMPSCN

>NP_478384.1

MKNELGKFLEENELELGKFSESDMLEITDDEVYAAGTPLALLGGAATGVIGYISNQTCPT

TACTRAC

>NP_478383.1

MKSSFLEKDIEEQVTWFEEVSEQEFDDDIFGACSTNTFSLSDYWGNKGNWCTATHECMSW

CK

>NP_834755.1

MSEIKKALNTLEIEDFDAIEMVDVDAMPENEALEIMGASCTTCVCTCSCCTT

>AHJ59549.1

MSKGYKFTKEELVEAWKDPQVREKLKDLPKHPSGKALNELSEEELAEIQGASDVQPETTP

LCVGVIIGLTTSIKICK

>WP_015792833.1

MTEEMTLLDLQGMEQTETDSWGGSGHGGGGDSGLSVTGCNGHSGISLLCDL

>CAG43551.1

MEKVLDLDVQVKGNNNTNDSAGDERITSHLFCSFGCEKTGSFNSFCC

>AAC69560.1

MSKKQIMSNCISIALLIALIPNIYFIADKMGIQLAPAWYQDIVNWVSAGGTLTTGFAIIV

GVTVPAWIAEAAAAFGIASA

>WP_067999479.1

MSAQGKDPNEIRRRFEELPMEVFQLDGSGLPIESLTDGHGMTEVGASCTSCVCICSCCT

>EFI65094.1

MTNEEIIVAWKNPKVRGKNMPSHPSGVGFQELSINEMAQVTGGAVEQRATPATPATPWLI

KASYVVSGAGVSFVASYITVN

>EFI65095.1

MTNEEIIVAWKNPKVRGKNMPSHPSGVGFQELSINEMAQVTGGAVEQRATPTLATPLTPH

TPYATYVVSGGVVSAISGIFSNNKTCLG

>BAN83916.1

MTEKTQITDVQAFEDLVAKVQEMDGPAQASSTVAALAGLDAAELQNFLEEKSGISPDEEA

QGSVMAAAASIALHC

>WP_013079674.1

MTKTHRLIRLGDAQRLTQGTLTPGLPEDFLPGHYMPG

>SED43766.1

MTEQSEQTPTEYIPPMLVEVGEFTEDTLGNWHGTSPDWFFNYYW

>EGD17355.1

MDTSNNDARTTALDQDLIVLGVASLDTQGGPLAGEEMGGITTLGISQD

>EDY58505.1

MLISTTNGQGTPMTSTDELYEAPELIEIGDYAELTRCVWGGDCTDFLGCGTAWICV

>EFE76491.1

MQKSVGHNGRQPRRREGVMKQQKQQKKAYVKPSMFQQGDFSKKTAGYFVGSYKEYWSRRI

I

>WP_043998581.1

MDKKNILPHQGKPVLRTTNGKLPSHLAELSEEALGGAGMDASFFPCSYDGADASFFPVCS

YDGADASFFPCSYDDGDA

>CAP64339.1

MRITPMDKKNLLPNQGAPVIRGISGKLPSHLAELSEEALGGNGAEASATVSICAFDGAEA

SFTGCMCAFDGAEASITGCICAFDGAEASITGCICAFDGDEA

>sp|Q07642.1|LANSB_STRGR

MALLDLQAMDTPAEDSFGELRTGSQVSLLVCEYSSLSVVLCTP

>NP_604414.1

MISSHQKTLTDKELALISGGKTHYPTNAWKSLWKGFWESLRYTDGF

>NP_345049.1

MNTKMLSQLEVMDTEMLAKVEGGYSSTDCQNALITGVTTGIITGGTGAGLATLGVAGLAG

AFVGAHIGAIGGGLTCLGGMVGDKLGLSW

>AAB91455.1

MNTITICKFDVLDAELLSTVEGGYSGKDCLKDMGGYALAGAGSGALWGAPAGGVGALPGA

FVGAHVGAIAGGFACMGGMIGNKFN

>ZP_04066356.1

MRTMEEQIFNSMIQQGAFAALFVWMLFTTQKKNEQREEQYQKVIEKNQQVIEEQAKAFSS

LSKDLSDVKQKILGNGDEK

>ZP_04066940.1

MRTMEEQIFNSMIQQGAFAALFVWMLFTTQKKNEQREEQYQKVIEKNQDVITKQAEAFGD

LSKDVSEIKQKILGSGDVQ

>NP_345050.1

MDTKMMSQFSVMDTEMLACVEGGGCNWGDFAKAGVGGGAARGLQLGIKTGTWQGAATGAA

GGAILGGVAYAATCWW

>NP_345056.1

MDTKIMEQFHEMDITMLSSIEGGKNNWQTNVLEGGGAAFGGWGLGTAICAASGVGAPFMG

ACGYIGAKFGVDLWAGVTGATGGF

>NP_345057.1

MNTYCNINETMLSEVYGGNSGGAAVVAALGCAAGGVKYGRLLGPWGAAIGGIGGAVVCGY

LAYTATS

>NP_345058.1

MDTKMMSQFAVMDNEMLACVEGGDIDWGRKISCAAGVAYGAIDGCATTV

>YP_140101.1

MATQTIENFNTLDLETLASVEGGGCSWGGFAKQGVATGVGNGLRLGIKTRTWQGAVAGAA

GGAIVGGVGYGATCWW

>AAG29818.1

MNTKTFEQFDVMTDEALSTVEGGGKGYCKPVYYAANGYSCRYSNGEWGYVVTKGAFQATT

DVIANGWVSSLGGGYFGKP

>AAC95138.1

MHKVKKLNNQELQQIVGGYSSKDCLKDIGKGIGAGTVAGAAGGGLAAGLGAIPGAFVGAH

FGVIGGSAACIGGLLGN

>AAC95139.1

MKKELLNKNEMSRIIGGKINWGNVGGSCVGGAVIGGALGGLGGAGGGCITGAIGSIWDQW

>AAG28763.1

MKKIEKLTEKEMANIIGGKYYGNGVTCGKHSCSVDWGKATTCIINNGAMAWATGGHQGTH

KC

>NP_297555.1

MRELTLTEIDNVSGADLGSRLSAAIVGGVAAFFAGSIWGGTRGGDGGGILGVGSIGQGVG

MVYGGIAGAIGGAIAGFVLDKDVIYSYTNGFMSSIFNGTFAK

>CAA90906.1

MIKREKNRTISSLGYEEISNHKLQEIQGGKGILGKLGVVQAGVDFVSGVWAGIKQSAKDH

PNA

>AAZ29031.1

MKKQILKGLVIVVCLSGATFFSTPQQASAAAPKITQKQKNCVNGQLGGMLAGALGGPGGV

VLGGIGGAIAGGCFN

>AAZ29032.1

MKIKWYWESLIETLIFIIVLLVFFYRSSGFSLKNLVLGSLFYLIAIGLFNYKKINK

>ZP_03980216.1

MTNFGTKVDAATRSYDNGIYCNNSKCWVNWGEAKENIAGIVISGWASGLAGMGH

>AAF44686.1

MKHLKILSIKQTQLIYGGTTHSGKYYGNGVYCTKNKCTVDWAKATTCIAGMSIGGFLGGA

IPGKC

>NP_863263.1

MKNIKNASNIKVIEDNELKAITGGGPGKWLPWLQPAYDFVTGLAKGIGKEGNKNKWKNV

>AAD28234.1

MQNVKELSTKEMKQIIGGENDHRMPNELNRPNNLSKGGAKCGAAIAGGLFGIPKGPLAWA

AGLANVYSKCN

>AAQ95741.1

MKKLTSKEMAQVVGGKYYGNGVSCNKKGCSVDWGKAIGIIGNNSAANLATGGAAGWKS

>BAA07120.1

MISMISSHQKTLTDKELALISGGKTYYGTNGVHCTKKSLWGKVRLKNVIPGTLCRKQSLP

IKQDLKILLGWATGAFGKTFH

>BAA82353.1

MKNFNTLSFETLANIVGGRNNWAANIGGVGGATVAGWALGNAVCGPACGFVGAHYVPIAW

AGVTAATGGFGKIRK

>NP_964623.1

MKLNDKELSKIVGGNRWGDTVLSAASGAGTGIKACKSFGPWGMAICGVGGAAIGGYFGYT

HN

>NP_542216.1

MDNLNKFKKLSDNKLQATIGGGMSGYIQGIPDFLKGYLHGISAANKHKKGRLGY

>NP_542217.1

MESNKLEKFANISNKDLNKITGGGFWGGLGYIAGRVGAAYGHAQASANNHHSPING

>YP_288875.1

MNALKRTCATLLISAGLTAGAVGVAAAAVEYVGGGIWDHGLTSSIVYSDYYHGSVCHGST

AVGTKTVRASAPAGYWSLADAPRAIANNQAYWRTTC

>NP_862432.1

MKTKSLVLALSAVTLFSAGGIVAQAEGTWQHGYGVSSAYSNYHHGSKTHSATVVNNNTGR

QGKDTQRAGVWAKATVGRNLTEKASFYYNFW

>ACR43769.1

MKELSEKELRECVGGGTWDDIGQGIGRVAYWVGKAMGNMSDVNQASRINRKKKH

>NP_268769.1

MIKFAEEIQKEELFHIIGGYSATDCKNHLIGGITSGAIAGGVGAGMATLGVGGVAGAFAG

AHVGAIAGGLTCVGGMLFNGK

>ACR43770.1

MKNNNNFFKGMEIIEDQELVSITGGKKWGWLAWVDPAYEFIKGFGKGAIKEGNKDKWKNI

>AAT72009.2

MNKTKSEHIKQQALDLFTRLQFLLQKHDTIEPYQYVLDILETGISKTKHNQQTPERQARV

VYNKIASQALVDKLHFTAEENKVLAAINELAHSQKGWGEFN

>NP_784211.1

MKIQIKGMKQLSNKEMQKIVGGKSSAYSLQMGATAIKQVKKLFKKWGW

>NP_784217.1

MLQFEKLQYSRLPQKKLAKISGGFNRGGYNFGKSVRHVVDAIGSVAGIRGILKSIR

>NP_784216.1

MKKFLVLRDRELNAISGGVFHAYSARGVRNNYKSAVGPADWVISAVRGFIHG

>NP_784205.1

MTVNKMIKDLDVVDAFAPISNNKLNGVVGGGAWKNFWSSLRKGFYDGEAGRAIRR

>ZP_04015571.1

MKIKLTVLNEFEELTADAEKNISGGRRSRKNGIGYAIGYAFGAVERAVLGGSRDYNK

>NP_784207.1

MKSLDKIAGLGIEMAEKDLTTVEGGKNYSKTWWYKSLTLLGKVAEGTSSAWHGLG

>AAG02566.1

MTKTSRRKNAIANYLEPVDEKSINESFGAGDPEARSGIPCTIGAAVAASIAVCPTTKCSK

RCGKRKK

>AAX99121.1

MKKKFVSSCIASTILFGTLLGVTYKAEAATVHVAGGVWSHGIGKHYVWSHYSHNKRNHGS

TAVGKYSSFSGVARPGVQSKASAPKAWGGNKTFYSLH

>NP_345048.1

MNTKMMEQFSVMDNEELEIVSGGRGNLGSAIGGCIGAVLLAAATGPITGGAATLICVGSG

IMSSL

>NP_720757.1

MNTKMMEQFETMDAETLSHVTGGGLYDGANGYAYRDSQGHWAYKVTKTPAQALTDVVVNS

WASGAASFAAYA

>YP_279852.1

MILFFMIFCTSSRLQRDKFKNYEKKLFDMEIKKLETFHQMTIEKLAKVEGGKNNWQANVS

GVIAAGSAGAAIGFPVCGVACGYIGAKTAITLWAGVTGATGGF

>AAP44569.1

MEAIKKLDLQAMKGIVGGKYYGNGLSCNKSGCSVDWSKAISIIGNNAVANLTTGGAAGWK

S

>AAP44566.1

MKNVQSLSKEELVLVVGGYTAKQCLQAIGSWGIAGTGAGAAGGPAGAFVGAHVGVIAGSA

VCIGGFLGQ

>AAP44567.2

MKTANIKLLTNQEMIEIFGGKTNWGSVVGSCVAGGLVGALGGTPISIGAGCLVGAGQDWI

SQK

>AAY68489.1

METAVAYYKDGVPYDDKGQVIITLLNGNPDGSGSGSGGGGGTGGSKSESSAAIHATAKWS

TAQLKKTQAEQAARAKAAAEAQAKAKANRDALTQHLKDIVNEALRHNSTHPEVIDLAHAN

NAAMQAEAERLRLAKAEEKARKEAEAAEKAFQEAEQRRKEIEKEQAETERQLKLAEDEEK

RLAALSEEARAVEVAQKNLAAAQSELAKVDEEINTLNTRLSSSIHARDAETNTLSGKRNE

LDQASAKYKELDERVKLLSPRANDPLQSRPFFEATRLRARAGDEMEEKQKQVTASETRLN

QISSEINGIQEAISQANNKRSTAVSRIHDAEDNLKTAQTNLLNSQIKDAVDATVSFYQTL

SEKYGEKYSKMAQELADKSKGKKISNVNEALAAFEKYKDVLNKKFSKADRDAIFNALEAV

KYEDWAKHLDQFAKYLKITGHVSFGYDVVSDILKIKDTGDWKPLFLTLEKKAVDAGVSYV

VVLLFSVLAGTTLGIWGIAIVTGILCAFIDKNKLNTINEVLGI

>YP_025360.1

MSGGDGKGHNSGAHDSGGSINGTSGKGGPDSGGGYWDNHPHITITGGREVGQGGAGINWG

GGSGHGNGGGSVAIQEYNTSKYPNTGGFPPLGDASWLLNPPKWSVIEVKSENSAWRSYIT

HVQGHVYKLTFDGTGKLIDTAYVNYEPSDDTRWSPLKSFKYNKGTAEKQVRDAINNEKEA

VKDAVKFTADFYKEVFKVYGEKAEKLAKLLADQAKGKKVRNVEDALKSYEKYKTNINKKI

NAKDREAIAKALESMDVGKAAKNIAKFSKGLGWVGPAIDITDWFTELYKAVETDNWRSFY

VKTETIAVGLAATHVAALAFSAVLGGPVGILGYGLIMAGVGALVNETIVDEANKVIGL

>sp|Q47502.1|CEAK_ECOLX

MAKELSGYGPTAGESMGGTGANLNQQGGNNNSNSGVHWGGGSGHGNNGGQGNSNSSGSTS

TVMKTGESYLTPWGDVVINNDGLPVMNGIVMTEENSTLVDNPFGGVSRVLNSLISDMPSL

FAESSGNNNNNTASVNTAPTNAQVSDMDKSSKVVSNVINEKQKQKNKIATQISEKQKKIE

EMKKVFKHHSYHGITDLERDVDELQKKSNQLDADISKLNSYKNTLQSKIGDVNKQKEAEE

KARENAEVAEHETLNEEKQAVAEAEKRLAEAKAELAKAESDVQSKQATVSRVAGELENAQ

KSVDVKVTGFPGWRDVQKKLQRQLEAKQAEYSAVENELKNAVSFRDGKAAEVKEAEQKLK

EAQDALEKSQIKDAVDTMVGFYQYITEQYGEKYAKIAQDLAEKSKGKKIQGVDEALAAFE

KYKNVLDKKFSKVDRDAIFNALESVNYDELSKNLTKISKSLKITSRVSFLYDVGSDFKNA

IETGNWRPLFVTLEKSAVDVGVAKIVALMFSFIVGVPLGFWGIAIVTGIVSSYIGDDELS

KLNELLGI

>AAD35867.1

MVNMEFLKRSFAPLTEKQWQEIDNRAREIFKTQLYGRKFVDVEGPYGWEYAAHPLGEVEV

LSDENEVVKWGLRKSLPLIELRATFTLDLWELDNLERGKPNVDLSSLEETVRKVAEFEDE

VIFRGCEKSGVKGLLSFEERKIECGSTPKDLLEAIVRALSIFSKDGIEGPYTLVINTDRW

INFLKEEAGHYPLEKRVEECLRGGKIITTPRIEDALVVSERGGDFKLILGQDLSIGYEDR

EKDAVRLFITETFTSRLSTRRP

>CAA44310.1

MPGFNYGGKGDGTNWSSERGTGPEPGGGDKGHSGDRDRGGAGVGNSPEQQQIAAIQNDPA

LRMKLEAVIKAARRINPDAKLHIESVSPSGTLSLSATGLTADQAKHIGLGGLVMGVNAKG

VTVAIGDIETGHARKPSPPGKGGNNGLNAGQIGASSLGSFVTDSHRDRPVSGWHGNGKTG

EFSTTRTTGSYYGFHHLKVEKQDGLATYSLYYKANKNRPAFIAVVRGDNLNAMEVKYANG

KPVKSPGSVKTIVKEFVEYQNAELKAIKDGVSLAAGINKDIAEKIGAKYAKLAKDLEAGI

QGKYIRNVQDAEKTYEQLTKGLNKKLKAQDKAAIVAWLKMIDAEQYARNARVLGKVFTGV

DWAIKGADLVNAAIEGFSTGNWKAFRNQLEALGLSIGAGYTLSAIAAFFAPTLVSSTVGI

FAFAYLFGWATSYIDAERAGELEKWVADL

>CAA72509.1

MPGFNYGGHGDGTGWSSERGDGPAPGGGMQGNGGGHSGNNDSGSNSVSQQISAIQNDQKL

KQKVVNMLIAARKMNPDAKMILGSIAPSGVMQVTIEGVTSTQARQLGLGGLVMGYNASGV

IGAVGEIDTGHRLNASGASTPGSETSVDSFVNGQKPAEEWHAVAKDSWTGAGPVNTGLVN

NAIKSVRIIKKGYVTGVLTPEEVMNKAEYKAMRQAFDSLPLAKQGEAVRQIVAAWSLAYQ

DFPVNLKKEMGRVTERIVDAINLALILNQTESRLSESQKNVDVANQIISDTVKAINDVNK

KIAEKRNQQVSLTDLMNKKQKEVEDLKKIFKNHSYHRIRDAQREYDDARNKYALLASDIN

ALQAQVSGLTARKQQAEQNKAAAEKAKADAAAKAAAEKAAAEAKAKAEAEKARKEAEEKA

NDEKAVLTKASEIIISVGDKAGEYLGDKYKVLSREIADNIKNFQGKTIRSYDEAMASVNK

LMANPDLKINAADRDAIVNAWKAFDAEDMGNKFAALGKTFKAADYVMKANNVREKSIEGY

QTGNWGPLMLEIESWVLSGIASAVALSFFSAIFGTFAMLGVFSTSLAGILAVILAGLVGA

LIDDNFVDKLNNEIIRPAY

>NP_061654.1

MVCFKSQRGMNMPGFNYGGYGDGTGWSSESGGPAPGGGMHGNSGGQRGDNANSSNSVSQQ

ISAIQNDQKLKQKVVNMLIAARKMNPEAKMILGSIAPSGVMQVTIEGVTSTQAKQLGLGG

LVMGYNASGVIGAVGEIDTGHRLNASGASTPGSETSVESFVNGQKPAGEWHAVAKDSWTG

AGPVNVGLVNNAIKSVRIIKKGYVTGVLLPEEVMNKAEYKAMRQAFDSLPLAKQGEAVRQ

IVAAWSLAYQDFPVNLKKDMGRVTERVVDAVNLALILNQISSGMSASQKDVDAANRIINE

TVKAINDVNLKIAEKKKQQVPLLSLMKQKQKEVEELKKVFKNHSYHRIRDAQRAYDDARN

KNDLLVSDINALQAQVSGLTARKQQAEKNKAAAEKAKADAKAKAEAEKAAAEAKAKAEAE

KARKEAEAKANDEKAVLTKASEIIISVGDKVGEYLGDKYKALSREIAGNIKNFQGKTIRS

YDEAIASVNKLMANPDLKINAADRDVIVNAWKAFDAEDMGNKFAALGKTFKAADYVMKAN

NVREKSIEGYQTGNWGPLMREVESWVVSGIASAVALAIFSATLGAYLLAVGASAAVVGII

GIIIASFIGALIDDKFIDRLNNEIIRPAY

>sp|P04480.1|CEA_CITFR

MPGFNYGGKGDGTGWSSERGSGPEPGGGSHGNSGGHDRGDSSNVGNESVTVMKPGDSYNT

PWGKVIINAAGQPTMNGTVMTADNSSMVPYGRGFTRVLNSLVNNPVSPAGQNGGKSPVQT

AVENYLMVQSGNLPPGYWLSNGKVMTEVREERTSGGGGKNGNERTWTVKVPREVPQLTAS

YNEGMRIRQEAADRARAEANARALAEEEARAIASGKSKAEFDAGKRVEAAQAAINTAQLN

VNNLSGAVSAANQVITQKQAEMTPLKNELAAANQRVQETLKFINDPIRSRIHFNMRSGLI

RAQHNVDTKQNEINAAVANRDALNSQLSQANNILQNARNEKSAADAALSAATAQRLQAEA

ALRAAAEAAEKARQRQAEEAERQRQAMEVAEKAKDERELLEKTSELIAGMGDKIGEHLGD

KYKAIAKDIADNIKNFQGKTIRSFDDAMASLNKITANPAMKINKADRDALVNAWKHVDAQ

DMANKLGNLSKAFKVADVVMKVEKVREKSIEGYETGNWGPLMLEVESWVLSGIASSVALG

IFSATLGAYALSLGVPAIAVGIAGILLAAVVGALIDDKFADALNNEIIRPAH

>sp|P05819.3|CEAB_ECOLX

MSDNEGSVPTEGIDYGDTMVVWPSTGRIPGGDVKPGGSSGLAPSMPPGWGDYSPQGIALV

QSVLFPGIIRRIILDKELEEGDWSGWSVSVHSPWGNEKVSAARTVLENGLRGGLPEPSRP

AAVSFARLEPASGNEQKIIRLMVTQQLEQVTDIPASQLPAAGNNVPVKYRLTDLMQNGTQ

YMAIIGGIPMTVPVVDAVPVPDRSRPGTNIKDVYSAPVSPNLPDLVLSVGQMNTPVRSNP

EIQEDGVISETGNYVEAGYTMSSNNHDVIVRFPEGSGVSPLYISAVEILDSNSLSQRQEA

ENNAKDDFRVKKEQENDEKTVLTKTSEVIISVGDKVGEYLGDKYKALSREIAENINNFQG

KTIRSYDDAMSSINKLMANPSLKINATDKEAIVNAWKAFNAEDMGNKFAALGKTFKAADY

AIKANNIREKSIEGYQTGNWGPLMLEVESWVISGMASAVALSLFSLTLGSALIAFGLSAT

VVGFVGVVIAGAIGAFIDDKFVDELNHKIIK

>sp|P17998.1|CEAD_ECOLX

MSDYEGSGPTEGIDYGHSMVVWPSTGLISGGDVKPGGSSGIAPSMPPGWGDYSPQGIALV

QSVLFPGIIRRIILDKELEEGDWSGWSVSVHSPWGNEKVSAARTVLENGLRGGLPEPSRP

AAVSFARLEPASGNEQKIIRLMVTQQLEQVTDIPASQLPAAGNNVPVKYRLMDLMQNGTQ

YMAIIGGIPMTVPVVDAVPVPDRSRPGTNIKDVYSAPVSPNLPDLVLSVGQMNTPVLSNP

EIQEEGVIAETGNYVEAGYTMSSNNHDVIVRFPEGSDVSPLYISTVEILDSNGLSQRQEA

ENKAKDDFRVKKEEAVARAEAEKAKAELFSKAGVNQPPVYTQEMMERANSVMNEQGALVL

NNTASSVQLAMTGTGVWTAAGDIAGNISKFFSNALEKVTIPEVSPLLMRISLGALWFHSE

EAGAGSDIVPGRNLEAMFSLSAQMLAGQGVVIEPGATSVNLPVRGQLINSNGQLALDLLK

TGNESIPAAVPVLNAVRDTATGLDKITLPAVVGAPSRTILVNPVPQPSVPTDTGNHQPVP

VTPVHTGTEVKSVEMPVTTITPVSDVGGLRDFIYWRPDAAGTGVEAVYVMLNDPLDSGRF

SRKQLDKKYKHAGDFGISDTKKNRETLTKFRDAIEEHLSDKDTVEKGTYRREKGSKVYFN

PNTMNVVIIKSNGEFLSGWKINPDADNGRIYLETGEL

>sp|P09883.4|CEA9_ECOLX

MSGGDGRGHNTGAHSTSGNINGGPTGIGVSGGASDGSGWSSENNPWGGGSGSGIHWGGGS

GRGNGGGNGNSGGGSGTGGNLSAVAAPVAFGFPALSTPGAGGLAVSISASELSAAIAGII

AKLKKVNLKFTPFGVVLSSLIPSEIAKDDPNMMSKIVTSLPADDITESPVSSLPLDKATV

NVNVRVVDDVKDERQNISVVSGVPMSVPVVDAKPTERPGVFTASIPGAPVLNISVNDSTP

AVQTLSPGVTNNTDKDVRPAGFTQGGNTRDAVIRFPKDSGHNAVYVSVSDVLSPDQVKQR

QDEENRRQQEWDATHPVEAAERNYERARAELNQANEDVARNQERQAKAVQVYNSRKSELD

AANKTLADAIAEIKQFNRFAHDPMAGGHRMWQMAGLKAQRAQTDVNNKQAAFDAAAKEKS

DADAALSAAQERRKQKENKEKDAKDKLDKESKRNKPGKATGKGKPVGDKWLDDAGKDSGA

PIPDRIADKLRDKEFKSFDDFRKAVWEEVSKDPELSKNLNPSNKSSVSKGYSPFTPKNQQ

VGGRKVYELHHDKPISQGGEVYDMDNIRVTTPKRHIDIHRGK

>AAG29099.1

MKNILLSILGVLSIVVSLAFSSYSVNAASNEWSWPLGKPYAGRYEEGQQFGNTAFNRGGT

YFHDGFDFGSAIYGNGSVYAVHDGKILYAGWDPVGGGSLGAFIVLQAGNTNVIYQEFSRN

VGDIKVSTGQTVKKGQLIGKFTSSHLHLGMTKKEWRSAHSSWNKDDGTWFNPIPILQGGS

TPTPPNPGPKNFTTNVRYGLRVLGGSWLPEVTNFNNTNDGFAGYPNRQHDMLYIKVDKGQ

MKYRVHTAQSGWLPWVSKGDKSDTVNGAAGMPGQAIDGVQLNYITPKGEKLSQAYYRSQT

TKRSGWLKVSADNGSIPGLDSYAGIFGEPLDRLQIGISQSNPF

>AAT85004.1

MADNQPVPLTPAPPGMVSLGVNENGEEEMTVIGGDGSGTGFSGNEAPIIPGSGSLQADLG

KKSLTRLQAESSAAIHATAKWTTENLAKTQAAQAERAKAAMLSQQAAKAKQAKLTLHLKD

VVDRALQNNKTRPTVIDLAHQNNQQMAAMAEFIGRQKAIEEARKKAEREAKRAEEAYQAA

LRAQEEEQRKQAEIERKLQEARKQEAAAKAKAEADRIAAEKAEAEARAKAEAERRKAEEA

RKALFAKAGIKDTPVYTLEKTKAATTLFLTPGVRLLNRAPAMIQLSALAAEINGVLTTAA

SAVMTATAEFSGWIASALWRGVAGVATASTVGPMVAAASTLFFSPRAGGGSDSKVPGRDI

EMLAAQARLFTAGKLSIEPGMKSVNLPVRGFISSETDGRQSLMLVKTGSDGVPSTVPVLD

AVRDSTTGLDKITVPAMSGAPSRTILVNPVPIGPAAPWHTGNSGPVPVTPVHTGTEVKQA

DSIVTTTLPIADIPPLQDFIYWQPDASGTGVEPIYVMTSQPRKGVKDYGHDYHPAPKTEE

IKGLGELIESRKKTPKQGGGGRRDRWVGDKGRKIYEWDSQHGELEGYRASDGSHLGAFDP

NTGKQLKGPDPKRNIKKYL

>AAL73547.1

MKTNNVTGTMKKVISTLAATGCMFSMAAAIPANSTIGSAVLGNAVVADAAVISVNTVVDA

KNGNADLVQGKFYKSPSQNYVLVFQNDGNLVIYHYNKTTDKAYSPIWSSQTENRGGTKCV

LQGDGNFVIYRSDGKPIWNTQTNGKKGAYLTISDEGEIKITSRNYNYATTWSSKNNHGYS

INQGPIITDPVDGQLSPHFHSREFACDCGNTHTIDQNLINKLEQLYTKLNCSKIIVNSGY

RDPNCSVAVGGGYDDAHTRGLAADVVCYDKNGNVIPCLTVAWAAEQIGFTGIGLMYGGAI

HLDVRTTSNYKNGHWFGDERKEYKNDYISTFKNYVPHKA

>AAT90329.1

MSDITYNPEDYNNGIPPEPGLVWKPGGSFPNGSYVPGSWGWPTRGYDVPPLPGDTEMLTV

TPKGTPADTWPKRPDIKEWYVPGEKPFDPSTGNGWVPDVDGYAESLPAGIPAVVQAAISK

VKGAPLKGGMSAVDIWKLKPATEYPGRFNSTDPAFSWFPVRALTDTDISAMPVAPETVPV

HTRILDNVHDGVQFVSAVFAGSMQYNLPVVKAQATAGSDYYTIGRLPGIMSAFTFSFYTK

GTPQDSRFFRDTVKAGGDLREAGFTVGANTSDFIIWFPQGSGLEPLYFSMTMNMPAGPLQ

RRQEAENKARAEADRLRAEAEAKIRAEAEARAKAEAERKALFAKAGIQDTPVYTPEMVKA

ANAALSAGGSMALSRAPGMIQHSAAGVGTLPFNSSLAGWEAGALWRGVDVLARIAPVASA

VATVATVLTLVRAALDIPAAGEGSDRVPGRNIDMLAAQASLYTAMKTNIQPGMKTVDLPV

RGYISYDGNGRQSVNLVRTGTGGVSATVPVLSAVRDKTTGLDKITVPAVAGAPSRTILIN

PVPVGPATPSHTGSSTPVPVTPVHTGTDVKQADSIVTTTLPAADIPALQDFIYWQPDATG

TGVEPIYVMLSDPLDSGKYTRRQLQKKYKHAIDFGITDTKINGETLTKFRDAIEAHLSDK

DTFEKGTYRRDKGSKVYFNPKTMNAVIIQANGDFLSGWKINPAADNGRIYLETGDL

>NP_889019.1

MNNLYRDLAPISAAAWAQIEEEVARTFKRSVAGRRVVDVKDPGGFGLAAVGTGHLRGIAA

PQKGVDAKLREVKALVELTVPFELQRDEIDAVERGANDADWQPAKDAATELAYAEDRAIF

DGYKAAGIVGIREGSSNSRLELPTDAADYPAAVGRALEQLRLAGVDGPYSVLLGADAYTA

LSEGSDDGYPTIDHIKRIVSGDIIWAPALNGGCVLSTRGGDFELHLGQDLSIGYQSHTDK

VVRLYLRETLTFLMLTSEASVPVAPKG

>CAE09438.1

MDILRRENAQFPASIWSAIEKEAGLVFGKHLTGRKVVDFKGGLGIGFSSLPTGRVISSKE

KLGEASVGVRMNTPVIELKIPFSFPESEVEAILREANAFDISSIEKAAKKVCVAENELVF

YGLKKEGIEGLIPSIPHKPIKAKGDEILPAVAEGIKELVNSEIEGPYALLIQPQYFGKLF

GVAGNSGYPLTLKLAELLQGNNIIVAPALKSGALLVSLRGGDYELYSGMDIGVGYSEKKS

TNHELFFFETLTFRINTPEASIAIEW

>YP_426062.1

MNDLMRDLAPISAKAWAEIETEARGTLTVTLAARKVVDFKGPLGWDASSVSLGRTEALAE

EPKAAGSAAVVTVRKRAVQPLIELCVPFTLKRAELEAIARGASDADLDPVIEAARAIAIA

EDRAVFHGFAAGGITGIGEASAEHALDLPADLADFPGVLVRALAVLRDRGVDGPYALVLG

RTVYQQLMETTTPGGYPVLQHVRRLFEGPLIWAPGVDGAMLISQRGGDFELTVGRDFSIG

YHDHDAQSVHLYLQESMTFRCLGPEAAVPLRGLSQAATKA

>YP_366690.1

MNNLHRELAPISSSAWEQIEEEVARTFKRSVAGRRVVDVDGPEGPELSAVGTGHLVEVAA

PREQVNARLREVRTIVELTVPFELSRDAIDSVERGARDADWQPAKDAAQRLAFAEDGAIF

DGYAAASIVGIREGTSNNKLTLPADVSAYPDAISDALEALRLAGVDGPYSVVLGSDAYTA

LSEARDQGYPVLGHIKRIVSGEIIWAPAISGGCVLSTRGGDYELHLGEDVSIGYTSHTDK

VVRLYLRETFTFLMLTSEASVAVAPQANTTA

>sp|Q45296.1|LIN18_BRELN

MNNLYRELAPIPGPAWAEIEEEARRTFKRNIAGRRIVDVAGPTGFETSAVTTGHIRDVQS

ETSGLQVKQRIVQEYIELRTPFTVTRQAIDDVARGSGDSDWQPVKDAATTIAMAEDRAIL

HGLDAAGIGGIVPGSSNAAVAIPDAVEDFADAVAQALSVLRTVGVDGPYSLLLSSAEYTK

VSESTDHGYPIREHLSRQLGAGEIIWAPALEGALLVSTRGGDYELHLGQDLSIGYYSHDS

ETVELYLQETFGFLALTDESSVPLSL

>CAA90860.1

MSDTMVVNGSGGVPAFLFSGSTLSSYRPNFEANSITIALPHYVDLPGRSNFKLMYIMGFP

IDTEMEKDSEYSNKIRQESKISKTEGTVSYEQKITVETGQEKDGVKVYRVMVLEGTIAES

IEHLDKKENEDILNNNRNRIVLADNTVINFDNISQLKEFLRRSVNIVDHDIFSSNGFEGF

NPTSHFPSNPSSDYFNSTGVTFGSGVDLGQRSKQDLLNDGVPQYIADRLDGYNMLRGKEA

YDKVRTAPLTLSDNEAHLLSNIYIDKFSHKIEGLFNDANIGLRFSDLPLRTRTALVSIGY

QKGFKLSRTAPTVWNKVIAKDWNGLVNAFNNIVDGMSDRRKREGALVQKDIDSGLLK

>YP_050090.1

MFTDEIIWHDVITKYSVNNLSQDMLNDPSETMFVLGDVYKEQALEYYGYLRSELLKSKEL

ISNAEKSLIIALESRVKAEQDKKSADQKLKDEQEKDKGKAPELKLDDKIREQLGNRGWTE

QDVRDTVSKGAKGSAEDKCSPKKTPPDFLGRNDPASVYGEFGKYIVVNDRTGEVVQFSDK

SDPEWVDDSRINWGDKNE

>AAM95702.1

MAGRTRIPFNGVGTSVLPAYQTLSAGQYLLSPNQRFKLLLQGDGNLVIQDNGATVWVANE

QQPFSSTIPLRNKKAPLAFYVQYGAFLDDYSRRRVWLTDNSTFTSNDQWNRTHLVLQDDG

NIVLVDSLALWNGTPAIPLVPGAIDSLLLAPGSELVQGVVYGAGASKLVFQGDGNLVAYG

PNGAATWNAGTQGKGAVRAVFQGDGNLVVYGAGNAVLWHSHTGGHASAVLRLQANGSIAI

LDEKPVWARFGFQPTYRHIRKINPDQKPIDIWTWHF

>prf||1912296A

MSDVFDLGSMTTVATATGQYSFYTPPPPTPIPYLTYIARPGINKFDLPEGAKIKDLIKRY

QYIGSQIPAAIMIRGVQEEIKKSTNTALANVGAIVDGELAYLASQKKEKLNPAEATPLQM

ASAEKAAAVELLASKQKELADARTIANAFFGYDPLTVNYVNVMNEIYGRREDKDFSFDNW

SKSYSAAQKIRLIEAKISVLNSRSSALDGKVAELTRLQRLEDAQHAAEAARQTEAERLAQ

EQRQAEARRQAEEARRQAEAQRQAELQRLAEAEAKRVAEAEKKRQDEINARLQAIVVSES

EAKRIEEIYKRLEEQDKISNPTVTTPPAVDAGSRVDDALAHTGTRVTSGGETGATGGSGR

DVDTGTGQGGITARPVDVGSVSIPDRRDPKIPDQPRRDLGSLVPTFPDFPTFPSFPGVGV

PAAAKPLIPAGGGAASVSRTLKTAVDLLSVARKTPGAMLGQVAAVVATMAVSSFWPKLNN

GERQASFAIPVAELSPPLAVDWQAIAAAKGTVDLPYRLKTLNVDGSIQIIAVPTEPGSAA

VPVRALTLDSASGTYKYTTTGPGGGTILVTPDTPPGQIDPSSSTPAVPRGPLIMPGTLLI

PKEPQIESYPELDQREFNDGIYVYPEDSGIPPLYIVYRDPRDEPGVATGNGQPVTGNWLA

GASQGDGVPIPSQIADQLRGKEFKSWRDFREQFWMAVSKDPSALENLSPSNRYFVSQGLA

PYAVPEEHLGSKEKFEIHHVVPLESGGALYNIDNLVIVTPKRHSEIHKELKLKRKEK

>AAA23073.1

RFAHDPMAGGHRMWQMAGLKAQRAQTDVNNKQAAFDAAAKEKSDADAALSAAQERRKQKE

NKEKDAKDKLDKESKRNKPGKATGKGKPVGDKWLDDAGKDSGAPIPDRIADKLRDKEFKN

FDDFRRKFWEEVSKDPELSKQFNPGNKKRLSQGLAPRARNKDTVGGRRSFELHHDKPISQ

DGGVYDMDNLRITTPKRHIDIHRGQ

**Non-bacteriocin**

**--------------------------------------------------------------**

>WP_001030800.1

MNKDSTQTWGLKRDITPCFGARLVQEGHRLHFLADRAGFTGSFSEVQTLQLDEAFPHFVA

HLELMLLSCELNPRYAHCVTLYRNGLTGEADTLGSHGYVYIAILNRHGFNRHLRVI

>WP_050443533.1

MNFEQMKAVYEMVKAIYNKEERLVIGKEKLHLTHGINKNSFADFYRAFQKMLDGELHTRG

ISTDLRDFYLSQIYEDYGTKKLETALNAYMDFIIYYEKKHNNIKKKNERKIYQKHYELIK

HQSPERKGRVKVVEFYEGEFEQVFITKHERNTEARNKCIQAKGVKCVVCDFDFEKTYGEL

GKGFIHVHHINPISTKDGNYAINIENELVPVCPNCHAMLHRRKDKILSIEELKRIFHNK

>WP_159117600.1

MAKDTPSMAKNKIKRCLWAIYDQHPKKSEVDSLWTYFESKCAYCGVEIERSSRTGHVDHL

IPSAEGGSNSIHNHVLACARCNGDEKREEDWLTFLSKKSGKSSIFEQRRSNIEEWLSLMP

PNGTNTALKSEVEKVVDKALKDFDSAVAQVRSLINVNDRG

>WP_130071231.1

MSRNPYYIKMINSQRWKNLRCDKLRANPVCEVCEANGLSTLATEVHHKSPVESVSHELGM

KHLMFDRTNLQSLCHACHSEIHRRVFSHSKEAIQANNRRATERFADKFLK

>WP_169167565.1

MKVLKLSAQGVPQSWITLEQAVIHSAAGDVRWVAGSEVAVFHGGHNAVTGLQSVIAVNSI

IGTRGVSRINPFELKPGLANNKLFARDRNVCAYCGGHFDEHDLTREHIVPLAQKGADQWM

NVVTACRPCNHRKGPRTPEQARMPLLYAPYVPSLWEDFILRNRRILADQMEFLAAHLPRS

SRLLN

>WP_160213701.1

MALFLLEWWRMAKPFSDAFYHSKAWGRAREDALKRDSYLCQRCLAGGEITPATMVHHIEE

LTPANIDNPDITCGLDNLVSLCDLCHKKTHGWARAGATRQGLAFDADGNLICLAE

>WP_160212293.1

MPSNNVRYRNWKARTEQRRRILRECDGRVCPFCGRPMDASLDWWTDPADGRRKRHPYSIE

VDEIVPVSKGGSPIDPANLQGAHRICNQRAGAKNRRPKPRGDVTGGGLPASREW

>WP_159494819.1

MLVLRLNKAGMPQEWIDVEHAAKLYSQEKVLFELGSDAITLKGGWNHEGLQSQLTLSSII

ACDGKVTDMSGKVALTNRFLFRRDSYLCLYCGQKFSPKQLTRDHIIPRSRGGKDTWTNVA

TACQRCNHAKAAKTPEEANMPLLAVPFRPNIYERFYLMNRRILSDQMAFLKGHFSHKRNW

TCLD

>WP_142428358.1

MIDVTSKQARAKFYGSSEWRRLRQQCLERDHYECQWCKQEGKLTTQYDSVLEVDHIKELE

HYPQHALDIDNLRTLCKDCHNKRHGRFNYRESKRKRKWDDEWW

>WP_142426822.1

MIEVTTKTDRAKFYSSSQWKKLRLKALERDHYECQWCKEQGKVTTINDAILEVDHIKELE

YHPEFATDIDNLRTLCKECHNKRHSRMNYRGAERKKKFDDEWWGD

>WP_142422844.1

MTDEFYRWLLQLIREDRLVKFYQSPKWRRLREKAMKRDHYECQECRRLGKYHRVENVHHI

KEVKDRPDLALDLDNLICLCVEHHNEVHGRYLTALDKQEKKIESFANFDASERW

>WP_142422626.1

MGKKWNTEMFSEFVNSTYPDFEVRGEYVSSKNNILIYHKKCDREFSVIARNFKTRGTCSL

CNGKFKSNTSEFKDKVNTLTNDEYEVIGEYVTCKDKIELAHKKCGTIYFATPDDFINGGT

RCPRCFGNNRKTSKRFKNEVFNLFKNEYIVLGEYKNNKTPLLMKHDSKKCNHEFMVSPDA

FLRGSHCNKCGTEKRSGENHYKYNFSLTEEDRMARDMQNGEIRKWRDKIYLRDDYTCQVC

RIKGYKLNAHHLNSWDFYERERFDTDNGITLCEDCHRKFHKKYGYGHNTKKQFTLYLEEN

KPTTSIL

>WP_121698255.1

MSERRISEKELILPTLYLAVCNGGRITTSELIKQLTAMMRPSGIDAEILSGRNDTYFSQK

VRNLRSHNTLVAPGYAIYDDKGYAITQLGRDFVEARMDSLRYLLSSDFDYEDVRGHLDDV

TDGKVIPYDELVSEGETITMTATSHERSRKLRDAAVAHYTQDGVLKCCCCGFDFGSFYGD

KYGSSCIEIHHIKPIFMYEGRSEEQTIEEALDNLMPVCPNCHRAIHRNHVMRDELPDFIA

AIKASRKS

>WP_081722951.1

MERGSPHTWEVVDVATSRTGTAQYKHWRKRVLIAARDAGIAQCPHCGVRLDYTRGLQPNS

AEPDHILPVRWGGKNTLENGRVLCRRCNQSRGDGTRPKVKPRRAASVDVDW

>WP_001372261.1

MIEKICEVIDGEYVCDIDISVEEWKILLRDKKVFDDKSIAALKKWFIEPDHSCTCFDIGK

KYDLHSMSANGVINGLGGRVQKQLGRFEVKGVGKIASGTKFITVMKSREIKGNPKRNLWT

IREELVQAIKELDFFSTNESSSIDFYSDNDLITALEESNHFDVTQTFEYSEKAKPKKAAI

EVKNGLSYPRSKSVSKNALNKADYKCEINCDHPTFRRRNSPLNYTEPHHIVPMSKQDYFE

NSLDVEENIISLCCNCHKQIHLGKGFEDMLRKIYAERKDVLKKAGIEILLEDLILFYKME

GN

>WP_102372778.1

MSGNSRNAAQPEFREGSRCEVTLDRYERSEAARKACIAAHGATCAICGFDFSHTYGPTFA

GIIQVHHIVPLHVTGKEHEVDPMHDLIPVCPNCHVALHSKPDGTYLPDELRALMR

>WP_160214295.1

MFGFSVTLYLFIAGMGAGLYIASCLVEREMERARPPRDMQLLHQKALIISLLLVCAGSAF

LILDLTVPQKMYLVFKRPFGSVISFGAWLIALLTLMLAVRNGFWRIFATSRSPLIRFLKA

ATFLLACGVTLYTGFFLIGLKAISFWESLLVVALFAISSLSSGIACFSVLAAFSLRRSTT

PPVVCKADQVDTFLLAAEIIVLGIFVVSQLFGDAASAASSSRLISGELAWAFWLMLVGIG

LLFPFGLSVFGRANHKLSPLVVKGISACIGCFFLRYCIMEAGVRSFSLA

>WP_160213053.1

MTYEPIWGPIIAWYLFLAGLGGGAFVTSVFIRFRHPECTRLIRTGRIIAPAVVIIGLCLL

MFDAHAGFMHPLRFALLLTNFGSVMTWGVVFLAAFVVLALVALLLDLLKKPVWQWLDIAG

MVMGLCVAIYTGCLLGVCQGFPLWNNALLPVLFLVSAVSTGMAAVLLAGVFVAPEEFNAV

VSLKKFHFWLPVVEIALVMALLFITASNPSPAGWNSVVTLLCGDWAVAFWVLFIAVGLVI

PIALECWMLWIATPVVEESRTGQMISGFSDLGVLVGGFVLRLMIVSAALPITIVQPWIF

>WP_160213014.1

MQLLTKNKLLTGIFAVLGVAGIAAWAYQLAGGLGVTGMSNANSWGLYIAMFMFFVGLSAG

GLIVASSASVFHTTEYKKVAMPAIILSTVCICCAGAFVLIDLGGIQRIWHLFASPNVASP

LVWDICVITLYLVINVLYLRFMHKGAERAVSVLSRFALPTAVLVHSVTAWIFGLQIAKEG

WFSAIMAPIFVASAMDSGLALLLIVLIALNAAKLFETPKKLIASLAGLLATCIAIDGFLI

FCEVITMAYPGAEGAATLAVMVSGPTAPFFWAEVVGGLLVPFLMLVFAKNRQNTALVTVA

SVLVVAGVLCKRLWLLLTAFVIPNIVGAPGIMSDAWMMGGSYAPTAIEFLIVLGVPSLGA

LAFMAIGSKLLVPATAKEHAPARSGAAADLDLEAQVA

>WP_160213011.1

MSDLIAAYLFCAGAGSGAAFLAAVFECFVRAGAFRRARFADRRQAVSMRAVALSVYGAAL

VLLAFGMLCLVFDLGRPDLALKLFLRPNLTLSTFGAFALAVLALALMVLVALRLGRENQG

AIRRRIDGLSRAVVIVASAAVMAYAGLLLGQADGMPLFETPWLAVLFVASALASGLAVVM

LAVAVAGNGHVEAVRYLKRRLTVRLDVALIVLEAVAAGLYLAAIALGPAGTVALAPLLTG

AQGGLFVGGFGLGGLTVPLVLDLLQWRRPLPGWAYGLAAVATLLGALALRFALVQAAGPL

LSWAPVA

>WP_160212856.1

MLDTFVTVYLFLGGCAAAVVLVTCAWSLAVRAACGRRQPAPPVFGRLRVRCLLAGFVLLV

LAVLCLLLDLGRPQLFWLLFARPTSSLISIGSFLLMATLLVSGFLLGASVPGAPRSSRRV

LCSAEVVCCALSAGVMLYTGLYMACLEAVPLWNNPALPVLFALSSLSSGLSVVLIAASFA

DDRFLLAADCRRLRLAHAVSLAGEMVAVGAYLALAWGDGFARPGLEALLSPNDLGSWFVV

GFLGLGVALPLGAEVFAAMARRPMEAIPLDALCIIGGLVLRFCVVIAA

>WP_160212813.1

MFDALVIAYLFLGGTGAALGGLLGLLTLGQVLGLGESGRGQHLNGLSSGQHRRFFGFGNV

LAAAVCLLGAVCLLFDMERPDKVLVLLTSPNTSLVAMGAYSLGAVLLLSALAGVLHLHRR

TLPPAAGALLCAAQLVAACVTMTYTALLLMGFRAVAFFQTWALVGLFFCSSASCGLALAT

LTGMVLRVFPLERGHERTAAAEAALSLLEGLFLVLFMIHAHYAAPQAFAQLATGPLAWAF

WTVVVGCGVASPVAAWALPRFVRGYRLHGGLSPALVLLAGFALRFCVIAVV

>WP_160212742.1

MDFNEGGREAASAAGRSAERAKEKAQAWGGAALNAAIGVSGVLAVLGIVLWGIQLSGGMV

QTAMRNLDSWGLYITMFMFFVGLSAGGLIISSVPKAFGIKGFGGISKVAVYSSIACTVAA

IGFVVVDLGQPMRLWELFVYSNLGSPLMWDIIVLGTYLILSCVYLWAQVQSEKGKVSAAA

LRVISVIALVCAVLVHSVTAWIFGLQVGREMWHTALLAPWFVSSALVCGTGLVMLVCMGL

SKAGYLEFSRENLVKLAKLLGAFVCVDLYFFGCDLLTEAFPAAGGMEVVTMLVSGPLAPF

FWVEIIGCILCAAVCFVPSLRKPGLLAVGAVLAIAGIFCKRVQLLVGGFQLTNLDMPGPV

TSLSVTNWESGFSGAYSGLVYWPTPLEFGVALGVVALAVFIFCLGVKFLPLRPKED

>WP_068921157.1

MSVQADSVVAHRWALRSGVYRATAANGDLMLAAWPHTAMLGHASPQLLALLDALAEGPVP

VDEPGMSATLDRLRAGGWLSRTVSCAGRDLYTVTPLAAPTEAPAPAGELRLSRFAVLRNT

PEGLVLEMPGSWCDIRVHDPAVAALLADPSGDAGLPADAAAAVRADLVAAGMLVAEEEER

EPFERRQWSTHELWFHERSRLGNRGWFGGAHFGGTFWARGVHEPPPARPSPYPGEAVPLA

RPDLATLRRTDPTLTTVLEDRESVRDHDDDAPITAEQLGEFLYRCARVRLLRTIEGFEYS

SKPYPGGGSAYELEVYPIVRLAADLTAGMYHYDAHDHLLRPVQPLGHPSVRRLLKVATES

SVTKAPPQVLLVISARVGRIMWKYEAMGYALMLKHVGVLQQTMYAVATAMGLAPCALGSG

DDLAFTGATDRDRLTECAVGEFMIGSRRKELATWQL

>WP_121705446.1

MKRKTALLVAACAALMALGGCQKVNEAPTTEAAPQAETGAATKKDPAGEKAEEGKGVNKV

AYITAQRLGDDGPVDMVYRGIKAGCDEAGIEVHVVEAKKGEYEESMQAMVSEGYNLIFAV

FPELIDSVKAVSQQNPDVSFIHAICATKGDNLEGICCYEQQSSFVMGVLAAMTTKNNKVA

FVGGVDNPDTHRYLDGYKEGIEYVNPEIEVQTSWIGSFEDPAKAKELALVHYQNGADVLW

GSGGKSALGLYEAAKEMGEGYYVMGCTDDNNGRLPGQVLASHYEAWDTAAKDLVIDWNDG

IFEPGLKVLTLENGYAYCKLADESQCEIPQEVRDKVEEVTEQIKSGEIVVKSMPTYEEVI

ATLE

>WP_000146146.1

MASGDLVRYVITVMLHEDTLTEINELNNYLTRDGFLLTMTDDDGNIHELGTNTFGLISTQ

SEEEIRELVSGLTHSATGKDPEITITTWEEWNSNRK

>WP_007792748.1

MSEVTRYVVTVKFHEKSLTDINELNNHLTRGGFQLTLADDDGKIHELGTNTFGLVSALSE

KEVAELAEGLGEAALDQKPQVTVTTFENWLRDNDTV

>WP_188061558.1

MRPVKKLAVTSAVAMLSLGMTACGSNSTNNNSSAPAGNSGASGSSGGSAAGALKVGLAYD

VGGRGDHSFNDSAAKGLDEAKAEFGIKPTEVAATNGENDAARVSRLQQLAQSGNQAIVAV

GFSYAAAIGKVAKQFPNVKFAIIDDASPDSKGDNIDQITFTEEQGSYLAGAAAALKSKSG

HIGFVGGVEVPLIKKFQAGYVAGAKKVNPNIKIDSTYLTQAPDFSGFADPAKGKTAAQGM

FQNGADIVYHAAGKSGDGVFDAAKAAGSGKWAIGVDSDQAQTAPAGVRPIILTSMLKGVD

VGVKSFLKKVHDGNFKGGNSVYALKDGGVSLATTGGHIDDIKAKLDELKKGIEDGSIKVP

SA

>WP_083706534.1

MTRAPVIGWTMLAGAALAASAGSLPIQLGFAILAIGILGMAHGASDLAIVAPGRRPLFLF

LYVSVSLICLAWWTGYPEIALPLFLAASAIHFGVEDAPHGSLPERAARGISLVATPAILH

REGYGDILAFAAGHGISTTVLFLLIAAGAVATALVLIMAIRRRDGRLLIGTGALLVLPPL

IGFSIGFLVLHALPQTDQRREEIGCVSHRAYFRAVAPILLAALLIAAAVGAFFVYREGTG

VRALFAGIAALAMPHLLVTPWFEGRAGRPVAYSCPAISGRAQHPQT

>WP_076714729.1

MDWMAFLIGFTIMSLASLAIYAKGSKTSPSLHHTLLHAAVPFIAATAYLAMAFGIGTLIN

IDGSVTYLARYADWSVTTPILLASLVLLAFHERGKMGEVGGYLTAIIVLDVLMIVTGLIS

SLALVPVLKWVWYLWSCAAFVGVLYLLWVPLRAMAAERGEALGTAYRKNVVFLTVIWFLY

PIVFLVGPEGLKIISDPTSVWAILIMDVLAKVVYAFYAAANLKTALHDHRA

>WP_056438944.1

MARLSIFPLAGAILFPGMPLPLHIFEPRYRALVSDAMARDRRIGMVQPSGEGDTPSLYQM

GCVGRIAEVEAMEDGRYNLVLEGVSLFRIVRELEVTTPFRQVEAELLPVIDEDLLSLGRR

ASLEQESKRFADLQGYAVDWDAVGRLDDESLVNGIAQIAPFDVAAKQALLEAPDLEQRAE

LIIQLMQFFGRHDGEDRVTLQ

>WP_056438392.1

MRIDLTPYRRSTIGFDRLFDLLEANSRAASAENYPPFNLERLADDRYRITLAVAGFARDE

IEITAQQNMLLVTGKKDDKAGSPNFLHVGIANRSFERRFELADFVFVEDARLNDGLLVID

LVREVPEAMKPKTIAIKTGQPLAAVEHHAGEADEAKAA

>WP_162547883.1

MKNILLALAASAAAIVGVAAPAAAQDKTKVCFVHVGSKTDGGWTQAHDIGRQQLQEHFGD

KIETPYLENVPEGPDAERAIERMARSGCALVFTTSFGFMDATLKVAEKFPDVKFEHATGY

KTAANVATYNSRFYEGRFINGQIAGKMSKTGVAGYIASFPIPEVVAGINAFLHGARTVNP

EFKLKVIWVNTWFDPGKEADAAKALFDQGVDVLTQHTDTTAPMQVAEERGLKAFGQASDM

IAAGPTAQLSAIVDTWAPYYIKRTQAVIDGTWSSAQTFDGLKDGILSMAPYTNMPDDVKA

MAMDTEAKIKSGELKPFSGPINKQDGTPWLKEGESADDGTILGMNFYIEGVDDKLPQ

>WP_015068113.1

MTEAVKTPYPRTFSHIGISVPDLEAAVKFYTEVLGWYLIMKPTEIVEDDSAIGEMCTDVF

GAGWGKFRIAHLSTGDRVGVEIFEFSNQENPENNFEYWKTGIFHFCVQDPDVEGLAEKIV

AAGGKKRMKAPRYYYPGEKPYRMIYMEDPFGNILEIYSHSYELHYASGAYE

>WP_169634528.1

MTNLQKRSLLQTVALTAVAVAALVGCGKKEEVAPAAAPGAEAPAKSEPLKIAFMYVSPVG

DGGWTYQHELGRRAIQEKFGDRIETSFVESVPESADSERVMRDMAGQGSKLVFATSFGYQ

EFVQKAAADLKDVKFEHATGYKTAGNVATYDTKTFEGAYLAGIVAGGMTKTKTIGVVASV

PIPEVVRNINSFVLGAQSVDPAIKAKVVWVNEWFSPPKESEAATSLINGGVDVMYQNTNS

PAVLKTAQERGVRAFGKDGDMSAFAPQAHLGSAVIDWTPYYTKVTQDTLDGKWEGGSFWW

GVKEGAMDLVKIADDVPQEIKDRVAKAKAGMKDESFHVWTGPIQDNAGKEVLPAGKVGDN

AFLTGIDFYVHGVEGKVPGAK

>WP_159120456.1

MEHYLSLFVRSIFVENMALSLFLGMCTFLAVSKKVKTAMGLGVAVIVVLGISVPVNQIIY

VNILAPGALAWAGFPEADLSFLNFLTFIGVIAALVQILEMSLDKFFPALYNALGIFLPLI

TVNCAIFGGVAFAVQREYNLTESVVYGVGSGMGWAIAIVLLAAVREKLKYADMPDGVRGL

GSVFMIAGLMALGFQSFTGIQL

>WP_159120259.1

MDLATVIGMLGAIGFIVMAMILGGSLSMFIDVQSILIVFGGTLFVILSQFTLGQFFGAGK

IAGKAFMFKIESPEELIEKIVEMADAARKGGFLALEEAEISNEFMQKGVDMLVDGHDIEV

VRETLSKDISMTSERHDFGASFFKGMGDIAPAMGMIGTLIGLVAMLSNMDDPKAIGPAMA

VALLTTLYGAFFANVICLPIAFKLSVRAGEEKLNQSLVLDGIVGIADGQNPRVIEGVLKN

YLAASKRGSAEEE

>WP_159120252.1

MSLERFPLFPLSAHLLPEGRMALRIFEPRYVRMVKQACAENSGFVMCMLNSNGDKETNKH

IHKIGTYAQVVDFDMLDDGLLGIKVAGSHLVEVSNIEAEKDGLRTGDCKTIPQWQCDLAP

QQIAPMDERLKEIFGSYEELAALYESPKFDNPNWVLNRWLELLPVDGSQKQHFLAQRECT

SLLNYLSGLIG

>WP_105932012.1

MRTIDLSPLYRSFIGSDHLASLIDAASRAEKQSTYPPYNIELLGDDKYRVTMAIAGFSKD

DVSIQVEENTLTITGTKKAETEDKESKERKFLHKGISERNFERKFQLGDHVKVLAADMEN

GLLHIDMERVIPEAKKPRQIEIGSRLLENQ

>WP_105930623.1

MTQEETQIKSIPAKAYSVLEEWMNSITHGLGLIAAIIGLVFMVYRADNPLALTTAVIYGS

TLILMFLSSTLYHAISHDKAKGWLKLFDHSAIYLLIAGTYTPLLLVSIGGVLGITMTAVI

WCLAIGGVAFKLVAQHRFPKVSVMTYLLMGWIALGLIYPLYLALPGAGLWLLVAGGLCFS

LGVCFYVAKKVKYTHAIWHLFVIGGCSCHYFSIYYFVF

>WP_018697113.1

MESLNIFIRSIFIDNMVFAFFFGMCSYIAVSKSVKTALGLGAAVTFVMVMTVPLNYLLYE

FVLKAGALSWAGLPDVNLDFLTFIVFIATIAAFVQLVEMAVEKFSPTLYSQLGIFLPLIA

VNCAIMGGSLFMQQKVDALELTSLWQSIVYGLGSGLGWWLAIVMMAAIREKTTYSQIPAA

LKGPGIAFIITGLMGIAFMIFSGIQF

>WP_005856736.1

MFILMPVIFVLGILAIALEDKIKINKAAIALFMAISMWMILMFDAYNIFVERSSTIFQEF

LTQNPEMASLPPHEQFINFISNRAIVYHLGNVSETLFFVMCSMLIVDIVDKHGGFRAVTG

YIRTPNKRKLLWYISFATFFFSALLDNLAAAIVIMAVLRKLVPDRTDRLKYACMVIIAAN

AGGSWSPIGDVTTILLWVGKNISAMHQISHVFIPALVNMLVPLTIAHFWLFKKGSTLRVL

SEEEQGDEYIPEIPNRSRRMIFVIGVLSLALVPVFQMVTNLPPFLGVLLGLVILWFYTDL

MYSKLHMHESQKLRISQLLPNIDLATIFFFLGILMAVGALETSGQLGIMSAFLDKHVHEP

YLISFVIGALSSCVDNVALVAATMGMYPIVEQVADLSPYAQFFVSDGGFWTFLAYCAVTG

GSILIIGSATGVTVMGLEKIDFMYYTKRFSILALIGYCCGAGVYMLLFA

>WP_169169348.1

MNALVGRGGLFDEFFKDVNPGFYVRPLHGDPLPTPGQMKVDVKENDSGYTVCAEVPGVPK

EDIQVSVEGNVVSLRAEVRQQDQQTEGEKVLRSERYFGAVARSFQLPADIDAAQCKAKYD

NGVLTLTLPKKQGGNAQRLSIE

>WP_159490586.1

MTEVKQQTKTISAKAYSVLEEWLNSITHGIGCIAAIVGLIFMLYRAEDKLALTTAAIYGS

TLILVFLSSTLYHAISHQKAKGWLKLFDHSAIYLLIAGTYTPLLLVSIGGVLGITMTAII

WSLAMGGVAFKLIAQHRFPKVSVMTYLLMGWIALGLIYPLYLALPGAGLWLLVAGGLCFS

IGVCFYVAKKVKYTHAIWHLFVIGGCSCHYFSIYYYVV

>WP_142437480.1

MIKKISRYEMLNEILNCVTHGLGFILSIIALIALTTKAANLKSSIHVIAYLIFGIAQVLL

FFSSTIYHSLMFTKFKRVFQIIDHSSIYLLIAGSYTPYCLLAIGGTFGWGLYSFIWTCAI

AGIVYKNITMSKENKIPKYSMITYVLMGVFAILIIEPLYKSIGLTGVLLLVSGGLFYFLG

TYFYRSKNMNFSHPIWHIFVILGATYIYFSIFLTT

>WP_131622846.1

MNTLLIIILVVLMGLVVVSLVRGIVAFLQSHKADIDAGGQRQQDMQLLQNKMMFNRIKYQ

ALAIVVVAIIISIAR

>WP_121708026.1

MVGNKENKKRMEGWDITSLEAEGLHKCIRPGVQDCRAVSVYRLNLKQGSRFTLESGELEM

NPVLIRGRAKLSGAGLDGELEKLDSFYIPGDTGVGLEALEDCVFYIGAAPCEGYGKPFVR

KFDLSLPLGDIHQIHGHGVGQREVFFTLNHQVEASRLICGLTWGANGAWTSWPPHQHEKD

LEEVYCYFDMDAPRFGFHISYLKSGEVEDIVAHTVRSGSMVLAPAGYHPTVASPGTRNTY

FWILAAHSHASRRYDLAVLDPVYADT

>WP_121707743.1

MMFKVKDPGSALTHFIAMLLALAAATPLLVKAARSPEQTHILALTIFIISMVLLYAASTV

YHTLDISPKVNQILRKADHMMIFILIAGTYTPVCMLVLGDYTGWMLLALVWGIAFFGILI

NALWITCPKWFSSLIYIAMGWVCILAFGKIIAALPASAFGWLLAGGIIYTIGGVIYALKL

PLFNSRFKNFGSHEIFHLFVMGGSLCHYIMMYAFVA

>WP_106897292.1

MGTYMREPINGLTHLFGAILSFVGLLAMVIKASTTADSTLTIISVIIFGISMTLLYAASA

TYHLVVAKAHVIAFLRRLDHSMIFVLIAGTYTPLCLISLNGMTGWVLFTIISAIAVAGVS

FKLIWFHAPRWLSTALYIAMGWIVVFFSSSLAPVLGTNGMALLIIGGLIYTVGAFIYWLK

PKFMNFKHFGFHEIFHIFILLGSLFHFLCVYLYVL

>WP_016292657.1

MQITIREPGSAITHFIGMMMAIIATAPLLVKAAMEPGAASLASLAVFMLSMILLYGASAT

YHSVNFSERAIKIFRKIDHMMIFVLIAGSYTPVCMITLGGKLGYTLLAVVWGIAILGMSI

KALWITCPKWFSSIIYIAMGWVCVAVFGPLWRTLPASAFLWLLTGGIIYTIGGIIYALKL

PLFNSQHTHFGSHEIFHLFVMGGSICHFIFMYLYVA

>WP_032850333.1

MENKRYNNVEEWANTLSHGAGILLGVIAGYFLLAKAAAGAEPKWAVACVTVYLFGMLSSY

VSSTWYHGSRPGKLKELLRKFDHGAIYLHIAGTYTPFTLLVMRHAGGWGWGIFSFVWLSA

IVGFILSFKKLKEHSNLETACYIAMGACILVAMKPLMDHLAEMGAGPAFWWLIGGGVSYI

IGAVFYSLRKPYMHATFHLFCLGGSIGHIIAIWLIL

>WP_121698083.1

MKHKAIFIRDDNGTEVSAQSPVIVSASRGTDIPAFYADWFFRRLEKGYVRWRNPFSGQDS

YVSFENTRFIVFWSKNPAPLLPYLPMLKERGIGCYIHFTLNDYEAEGLEQNVPPLSQRIE

TFRRAVEALGRGAVVWRFDPLILTDKINIDTLLEKIAHIANALTGYTEKLVFSFADIESY

KKVSRNLRQSCINYREWDEESMCEFASRLSTKNHDNWNLRLATCAERIDLSEYGIGHNRC

IDPELISRLTPHDAILQNFLYNAKTDNGQRKACGCILSKDIGAYNTCPHGCLYCYANTSS

ASAFANYKEFATNPLTDLII

>WP_149888968.1

MSLKLVAAVAAVASAFALSACGDKKEAAPAKPATPAAPAAQTEAAEPLKVGFVYVAPIAD

VGYTKQHDIGRIYAIDKVGKDKVTTTFVENVPETADAERVIRQMVADGNKLIFGTSFGYM

NYMQKLAKEYPDVKFEHATGYKTAPNMTNYNIRFYEGRYLAGMLAGGATKSNIIGYVAPF

PIPEVLQGINAFTLGAKSVNPNIQVKVIWTNAWYDPPKDTDSAKTLLGQGADILTQHTNT

SAVASAAEAAGKMVIPYNSDMKSVAPNAQIAALVLNWGPYYAKKIQQTIDGKWDPTPVWM

HYKDGAMSLEGVRTDKIPADIVKKMEEVKAKIESGEFHPFTGPIKTNDGKEAAKAGEVLK

DNQLQTMNYYVDGVIGKVPN

>WP_010714134.1

MTKNKLFHLTDKSNINSIRKHGLVGVKKAKDLLRRETGTDHTLFNQQTYSILNTYGWKGY

DLRCATFMFEEDSCLGYELLQLMNDDPYILEIEIDRLNKDKLFVFNTEIASHLLNYSKQD

QHRLAKFYWNTAIPYNTYVQNKDKVNDAFQLGNIMYQAEYVYFGEISPKYIKNYERGKKY

DSGRY

>WP_007273123.1

MLMRTDPFRDLDRIAQQVFGTPARPAAMSMDAWRNGDTFEVEFDLPGIKPDSIDLDVERN

VVTVKAERPALDKDLEMLASERPRGVFSRQLVLGENLDTENIEASYEAGVLRLRIPVAEK

AKPRKISITAPSQDREAIDP

>WP_006681612.1

MREVVEYLEDRGVEHLVHFTPITNLGGIKKRGILPRNEIDGFPDIVFEALDEVRLDERTD

MSCFSISFPNFLMMYRYRTKLWSREEDVALLFIPISVLSDLEYDQVVFCPSNAASRECRR

TDPQDLLGLAAVEKLFVEEMTTRSGVVFSRQSEDLPDFLTTNPQAEIQIAATIPWEKVSF

VVVNDYETKQSLLQSGIHRNVYAKWEVNNDSSLNVFKYPSYWRAWVDAAGDLHG

>WP_121699091.1

MENLNIFIRSIFVDNMIFAYFLGMCSFLAVSKNVKTALGLGAAVTFMLVISLPINYLLET

YVLRAGALQWLGPEYADVDLSFLSLIMFIAVIASLTQLVEMAVEKFSPSLYSSLGIFLPL

IAVNCAILGGSLFMQQRDFPDVWTACCAGAGWGLGWLLAIVAIAAIRERLQEYSNIPKPL

RGVGITFILTGLMGIAFMSFLGIKL

>WP_121699089.1

MNKQSNTYTIIYIVALVIIVGTALAFTALSLKPLQTANADADKMKQILASVHIAPAKSDI

ITDFDKYITDRFVIDAEGKRVEGDAFAINVSAQSKLPQAERKLPVYECTLTPGDVKYILP

VYGAGLWGPIWGYVAVNSDGSSIYGAYFAHQGETPGLGAEIEKPAFSDQFTGLNLFKEGA

FKPVNVIKAGQAPMNGEDYVDAISGGTITSKGVASMLDNCLSGYKTFLESLTNKGQ

>WP_167508517.1

MSGSSRLAKFRTAQADKQNWQCFYCGFPMWEGDLALPSEHRRLPIGLLDRFLCTAEHLEP

KMNGGKNRPDNLVAACRFCNQTRHKMRDVLSPAAYQQHVRRRIRARKWHPIECHRLFG

>WP_160582050.1

MPYKPKRPCSYPGCPKLTDGRYCGEHQKIVTAHYNKHERDPASKKRYGRAWKRIRDRYIA

AHPLCEECRKAGKVTPAEEVHHIRPLSKGGTHAEGNLMALCKPCHSEITAREGGRWERRR

>WP_135855810.1

MIEFIDDTEPWRPTTSGERLLFLLLANHKEARFLSEFLAGSGLRFARRHSDWSNFLVGAS

LLHLQQRLIKLGNDVREKSPSDRMDQLKELRARTCDIVQLTPIEYEGDFGELVQEVLVSA

EQMHKEPPQNLKRSVLRSSPSCYSCGRNFGSVYENDEDAKEGLRATADHVWPRALGGDST

EDNLLPACTSCNSTKGHLATWHMAWLQPIVFSDVDGEHAPPPVPREVQMALHMRAATSYA

RANGTTLRDAFLAIGPRDRPEKIDSEQGYDFFNMRVHNETRTMVKWIPG

>WP_120438463.1

MPGWNLKNGELQKCQISEDEYWSLFNFVFSDACMKRNTYKFGLIKSIMDNLFNCTQDDYG

NYRLSYSAIFEKFTINYWNLVLKYHLKQMRSDGRTEVSKIESILLAASEENDLIKTLDFN

SLSNSDRSKVVRQVSIACRKNVIGALYNDMEGKLYGFDLKEKGIMLGERAYDFMLKYKTE

LEKLNYYAWAKFMEKINEDDVLVKVLDKLELSTPKRDDLSVYREVLYEEFEACNCFYCGK

KLSLSNRGIHVDHFIPWSYVKDDKLWNFVLSCPKCNERKNNKIPSKKYLEIMLKRNEYMK

GVVDEFVEIEFKNYDSSQFMRLWKYAQLSGMKQFQQEFF

>WP_120438225.1

MESYILHVRSVKESMSREVINTYQLEEILPKIIDYEEENYYNNIGVDSNLEKDLLEDLQM

LDNKYKEITKSVIIRYQKIVEHIKQTRGRKCQICQYSFIMDNGNEYCEAHHIQYLSKNGS

QSSDNVILLCPNHHRMFHYAHDAVFVDDLVDGKRKVLIENVEHLIDFS

>WP_140970856.1

MKEYKTKQQKRKFYDSGEWKSIREQVKKRDNYECQECKRNGRVQTDTNEYSESAKRKKIQ

LVVHHIKELEYHPELALEKDNLETACVDCHNKEHGRFFEKKPNKWENDEKW

>WP_169253183.1

MKTLVLNAGYEPLSIVPFTRAVVLVLTGKATVLAAEDIPVRSEHMSLDQPSVILLTRYVR

PPSNRRVSLSRRGVLRRDGHRCAYCSKPAYTVDHVLPRSRGGANTWENLVACCRECNNRK

GNRTLGEIGWKLSFLPQEPRLGQLWMRGIDKPVEKWRPFLEYSSAA

>WP_083793542.1

MMRLFNKRQRRILAWVAGGQCTICQRPLNSNFHADHVLAHSKGGATTTDNGQALCAPCNL

KKGAK

>WP_040799872.1

MAVAQTRRARAARRRKRRVDAADNDLTAEQWKELKREWGGCAYCTATDTVLQKDCVQPIS

RGGSYTVGNVVPACGSCNASKSNSEVTSWMRRKRLDERAFLTQYVHVRQALGLV

>WP_007230052.1

MVELEIEVGDEIKNEDLVRMFGCGPQGGMRRSHATNTLVLTSKHVDNVYDDRWVADVFHY

TGMGLEGDQSLTYSQNKTLAQSASNGVEVHLFEVFKPKFYTYMGAVELAAEPYVENQKDQ

NGLERKVYVFPLRPISGGQPTLPFKKIESAIHNKQKMAHKLSDQELLNRAASKSSAGASR

SVETKYYERDPWISEYAKRRAGGKCQLCESDAPFISKAGEPYLETHHIEWLANGGEDSIS

NTVALCPNCHRKMHNIADNNDVVKLKSRNASYE

>WP_007225374.1

MSTFLLTWSPDKWGYENLQEYLDARKSEEFVQRWSSGRTKKIPIGSRVFLTKQGKGNKGI

FGSGHVTKEPAEEPHFNEEQLKLGKKALFVMVNFDQLYDPQSEIPITHSELQAFDSKVWD

SQSSGITIPEETASKLEQLWLERTGAVEISYADEVPKDNSLKEGAAKKIWVNAYERNPDA

RERCIRKWGLNCVVCNFHFEQCYGHLGKRYIHVHHLKPLAEIQKEYEVNPEEDLRPVCPN

CHSMLHRNKNSVLSIEELQTLVNMYSR

>WP_169253656.1

MSTSEYDSYRPPEPEGGRKKRRRGGRGGPSGPDGPRGRRGGRGGGWKNRGADGNREMPMV

EDVEFTSYYGRPIVKAPPWGDEISAYLFLGGLAGGSSLLGYGAQLTDRPGLRIASRMTAI

AATGIGGVALVADLGRPERFLNMMRVVKVSSPMSLGVWILSGFGVGSGVTFAIELDRITG

EKLLPLGPLRKVLHGLETPAAVESAFFATPLAAYTAVLLGATAVPTWNAAGRNGLPYVFV

SSASMAAGGAAMALAPVGQTGPARLLALAGTAGEAYAMSAMKKRMHPAEVDPMDDGEPGH

KLHRAEKLLIAGTIGTAVAEVGARVFAKKLGGGWKTRAVLRGLSVVSGAALAAASAYTRF

GVLEAGIESTKDPRHVVEPQRARLEERRARGITDDSITTGR

>WP_141265030.1

MLVKALTGSKRYWGWITLLLVLIGTGFTCYLWQLDKGLTITGMSRDVSWGLYISQFTFLV

GVAASAVMVVLPYYLHNVKAFGRITILGEFLAVAALIMCLLFVLVDVGKPMRILNMIFYP

TPNSMFFWDMIALNGYLLLNIIAGWHALEAEYKAVPPPAWTKVLVYISIPWAVSIHTVTA

FLYAGLPGRHYWLTAIMAARFLASAFASGPALLILLCYIIKRVSKFDPGREAIQKLAAIV

TYATIVSTFFIGLEFFTAFYSQVPAHGIYTLKYLFAGLDGHSRLVSWMWAFAILVVFALV

LLINPGTRTRDSYLQLACAAVFVSMWIEKGIGLVIGGFVPNPFERVTEYVPTLPEILIAL

GVWATGFLVLTFLYKIAISVKEETV

>WP_160582917.1

MKLKIKDPGSALTHFIGMVLAILAATPLLVRAAHTPGPLHIAALAVFICSMILLYTASTV

YHTFDISESVNRLLRKIDHMMIFILIAGTYTPVCLIVLGNPAGYRLLALVWGIAVLGILI

NALWINCPKWFSSCVYIAMGWVCVTAFREIVAALSPAAFGWLLTGGIIYTIGGVIYALKL

PIFNSRHKNFGSHEIFHLFVMGGSFCHYMMMYGYIAA

>WP_160582365.1

MLMPSIFGEDLFDEMMGFPFDNRFFARRNPVYGKAATAVMKTDVKDMDGMYEISMDLPGF

SKENINAELKDGYLTVNATTSVNQDDSSEGRYIRRERYCGSMSRSFYVGDAVKKEDIKAR

FENGILALTIPKVPEQPKVETPNYIAIEG

>WP_160581716.1

MIISASRRTDLPACYPDWLFQRLKEEYVLVRNPMNAHQISRIDLSPKVVDGIVLWTKNPL

PLFRHLNELEKYSYYVQFTLTPYGPEAEPGLPSKNRVMIPAFCRLSREIGRERVVWRYDP

IFLSNVYTMEYHTKYFRVLASRLGEYTEKCTVSFLDLYQSTARNGRPLGIHTETGEQQLE

LMERFAEIAEEWGITIDTCAEQGDFGRFHVGRASCIDKDRLERIRGCQLKVKKDPNQRPE

CGCAASIDIGTYDTCRIGCLYCYANHHRDTVFKNSQRHNPASPLLFGEIGENDRIMERRM

ESCMDGQMSFHDWHG

>WP_135902375.1

MRSPEYAPLFRSTVGFDRLFDMLENSVRTDWPPYDIEKKGENEYRITMAVAGFSQEDVEL

TQHGPELTVTGQKSTAENGVQFLHRGLASRNFKQVFRLADHVKVANATLENGLLSIELVR

EIPEELKPRRISITSTATADPQPQISQDVKPGRKVA

>WP_135898580.1

MRHVDFSPLYRSTVGFDRLFTMLDSLAQPDGAQTYPPYNIERTGEDSYRISMAVAGFSDD

EISIEAHRNVLTVKGERKDEGTGEGSELLYRGIASRAFERRFQLADHVDVVGAALKNGLL

FVDLKRNIPEELKPRKIAINSAPAKAKQIEAKTAA

>WP_135873972.1

MPAQTDAAADWSGWRLAHAMAARFVRWAVDTEGSSRSSALIRIGLAMLFWSRWAGELLLY

MDQSPAGLFLAANFFVATTLLFVGYQSRVAAVWTGAVGLAMYHYFGFQLGREPWTHHHTY

LLAVSALLIALTPCGASYSLDRYLAVMRAERMGLPPPAERGNLLGLRLIVVQLSVLYFFA

AFDKSGYAFSSGARIEAIFLWYYAGSDYPAIPGLAWLATIVSLAVVALEYSLAFGLPFRA

TRRYLLLPGLAFHAIIYVTLPVYTFSATMVLLYLAYFDADAVDRVIARLQGIGPTAGEET

S

>WP_120424976.1

MTLKIKDPGSALTHFIGMLLALFAATPLLIKAARSPEQTHVLALTIFIISMILLYAASTT

YHTLDISPKVNQILRKVDHMMIFILIAGTYTPVCMLVLGDRTGWALLGLVWGIALAGITI

NALWITCPKWFSSLIYIAMGWVCVLAFGKITAALPKSAFGWLLAGGIIYTIGGIIYALKL

PLFNSRYKHFGSHEIFHLFVMGGSLCHYIMMYAFVA

>WP_135901799.1

MDDVPGNAARYRTGLAVRRSVLGDPHVDRAEIAATDFDQPFQELITEAAWGTVWARPGFS

KRERSIVTLALLAALGHDEEVAMHVRATANTGASRSDICEAFLHVAIYAGVPAANRAFKI

AKEVFSEMDGGKAVHAR

>WP_135901798.1

MPFVTLGGITLHHRYVEANGKTPAVVFINSLGTDFRIWDQLLSELDGEMPLLVYDKRGHG

LSDIGDIRSIDDHVDDLIGLIDHFGLDRLVLCGLSVGGMIAQGLYARRPEIVAAMILCDT

AHKIGTAESWNARIATVQANGIRAVADAVLKVWFTPPFHSERQPELDGYWNMLTRQALPG

YIGTCMAVRDADFTETARRIAVPTLCVVGDQDGSTPPDLVRSLADLIPGARFEIISDAGH

IPCVEHPAALVALIRDFVASLPSGETHG

>WP_135895629.1

MQRSLADSRVKFRHLQCFLAVAQFGSVQRAAGSLSITQPAVSKTVAELEDILGVKLFERG

RHGAVPTREGQLFMPHASACVSALRQGVDLLARAEGAAAATLEVGVLPTVAGALIPPVLK

RFASLWPRVIVRLATGANPELLERLKAGTIEFAIGRLADPERMVGLSFEQLFSEPLVAVV

RAGHPLDVTSGLPPAALQDFPVVLPPFGTLIRQSADSLLSAWGVLPLSAFVEVLSVSTGR

ALTLENDAVWFVPLSAVEYELTHGMFVRLPLPFAGTDEPVGLIRRSDTQPSPVGRAFIDA

VREVAQARMAASGGKAAGKVARKRGRGRTPAS

>WP_135891186.1

MPYAAVNGTELHYRIDGGRHGNAPWVILSNSLGSDLSMWTPQVAALSKHFRVLRYDTRGH

GHSEAPKGPYTIEHLAGDVLGLMDTLKIARAHFCGVSMGGLTGVALAARHASRFERVVLA

NTAARIGSPEVWVPRAARARTEGMLALADAVLPRWFTADYIEREPVVLAMVRDVFVHTDK

EGYALNGEAIDATDLRPETHGIKLPVLVISGTHDVAATPAQGRELAQAIPGARYVELDAS

HISNIEKADAFTKTVIDFLTESK

>WP_135872108.1

MAESRIRFRHLQAFLEVARQGSVARAADFLHVSAPAVTKTLRELEEALGVPVVERDGRGI

RVTRLGEIFLGHAGSAISALKRGVDSVRQDGALNRDPIRIGALPTVSARVMPLAMTLFLE

ENTGAALKIVTGENAVLLEQLRVGALDLVVGRLAAPEHMTGFFFEHLYSEQVLFVVRAGH

PLAEPGTDIFARLDEFPVLMPTRESVIRPFVDRLFITNGMTAPATEIETVSDSFGRSFMR

QSNAVWIISAGVVANEIASGAFVALPVDTEETKGPVGLTMRTDTAPSPAFTILLKTIREA

ARHHA

>WP_135855689.1

MHFLRCGETVIHYRVKGLDSGKPVIAFINSLGTDFRIWDAVTEVLGDDYAYVLHDKRGHG

LSDIGRPPYSIDDHAGDLIALLDHLGVKSAVIWGLSVGGLIAQGLYARRPDLVRALVLSN

TAHKIGTADMWNARIDKISADGLGSLVDPVMERWFTPAFRTPDNAAYAGARNMLAQQPEA

GYSGTCAAIRDADFTAAAGRIAVPTLCVVGDQDGSTPPELVKSLADLIPASRFVTIAGCG

HIPCLEQPLAYAQAACIFLKTLPEN

>WP_120446043.1

MAKGSAEKRRNMKKLLEEMEKNQELKAKIKELDENPKSTTKDYIQVAAENGLELTEADFQ

PAGSVGELADDELEAVAGGKDACTCVVGGGGQAYDEKTCVCVLGGGGEFTDGEARCICVA

IGTGQHG

>WP_120446041.1

MKRLLEEMEKNQELKAKIEELDKNPESTPKDYIRVAAEYGIEIKEEDFKPAQGELTDDEL

DAVAGGEPCACVFGGGGTANQSDDTCACVFGGGGEYSDGSCRCACVGGGAGDGHHYVPVS

DLFR

>WP_169251451.1

MDLTASVLAAPAAPADDARVLVVLPSLGTSAAALWQQAATELTALSPETTVIGIDLPGHG

RSAPIDVPAGTSVAPRITMSDLAEAVLATLDRVLPDVAAPGAPVDLAGDSIGGATALQLA

LDHGDRFGRIAVFCTGAKIGEATAWEERAQTVATSGTPTQVIGSAQRWFGEGFMDREPDA

SAALLHSLQDADRFSYAALCYALADFDVRARLPEITRPLLAVAGSQDQPTPATKLAEIAG

DVPGALLEVIDGAAHLVPAEAPVVTAGLLADFLQGKGLGAGAGGTGGAGATAASDSPADL

PTASGPREASRDEVREAGMTVRRQVLSDAHVDRANAKVDDFTSDFQDLITRYAWGEIWTR

PGLERRMRSAITLTAMIAGGHEAELAMHVKAALRNGLTRDEIKEVLLQSAIYCSVPSANT

AFSVASRALAEYEAEENAD

>WP_007228512.1

MKRTLLLLLAISIAACATGTDPAVTASAIDDFQPDARPTAKTMVYECDDAEFITRVGPGE

MALWFEDRYLILSQVRAASGTKYQEGDVVFWSKGDQVIFSAAGVRYANCQINHVRAPWED

ARRRGVDFRAVGNEPGWHLEVRGDQHLLFVGDYGATKIMFSNVHTTEDREQLHYLSQDDD

NSINVTVIESACIDTMKGDQFPYNVQVQLNDRNYQGCGRTLDHPWE

>WP_086005900.1

MSARLARFVKDPKHRLELLQDIHAASAQADLPPELVLSLIEVESHFDRFAISRVGAQGMM

QVMPFWKNEIGRPDDNLTLNKTNFAYGCRILQFYLQREKGDLHKGLARYNGSVGRRVYSD

KVYRAWNDHWRTEPLDWGD

>WP_040812715.1

MKQRVVLIEDELALAASYQDFLQRAGYDVTIYRTAESAVKGVEDCHPDLILLDIGLGHDP

EAGFELCRTLRARDALVPIVFLTARDEEVDVISGLRLGADDYLTKDISRSHLLARISGLL

RRVVALRNPENQEQVLKRGDLELNSERLTTTWGKSQIALTYTEFWMLYVMAKNPGHIKSR

EQLMEAARVVLDDSTVTSHIRRIRRKFETADKQFKHLETAYGLGYRWRGDA

>WP_007226915.1

MKNIQRSIGKSITALIMVLLGSSMIYQAGALMLEDTVDGKDNLYSSEWGHWFTMPGDGAL

AAYAPQSTAASAIVDSSNNAYDFSSWDYLDIVVTGSVTDAGSYETDAGGCTDPTASCWFG

DGQFRYQDVYSVIGIWSSSADEISWLDTIYDNWVDAVFTVGSDASIEVPEIEGAYLFLAE

NDGFFADNSGFYTATITTSVPEPASALLMLTALLGLFAIRRQRRAL

>WP_078486412.1

MRINIKLAAGFPLSRLESTFHGIETTRDENNTHTVRLTKSEVTADRDFELTWSPVPGNEP

HAALFSEQWAGDNYSLMMVIPPHQEGQGGALPREMVFVVDTSGSMHGASMGQAKAALKMA

LSRLAPDDRFNIIQFNSSTQALFGRAVGASPRNLARAEDYVDSLTASGGTEMLPALRRAL

TGEKELDRLRQVVFMTDGSVGNEAQLFEVIEQKLGASRLFTVGIGSAPNSFFMTRAARLG

RGSFTYIGKVSEVRSKMKALFNKLESPVLADVEIDWGEDVQVDMWPRRIPDLYMGEPLVL

AVKGEVDGKTVVIRGRSGDKPWQQRVTLHGGGSRGGIRLLWARKKIADLMDQKARGRTED

EVRHEVLVVALGHKLVSKYTSLVAVDKTPSRPLDQALIGKNVPVQLPKGWSAEKVFGSMP

QTATPALLNLLLGVLAMIGSWMVSVFGKRNRKNVADEVRHLNGEMYR

>WP_007229091.1

MQQHIAIVEDEAAIAANYRDHLQRQGFRVSLFADRDSAADAFAIQLPDLAIIDVGLGKEM

EGGFELCRDLRARAPGIPIVFLTARDSELDIISGFRLGADDYLTKGISQAQLTARINALF

RRVKALQKPEQEKHLVVQGALELNKERMTANWRGLPLELSVTEFWMVHTLALHPGHVKNR

QQLMDCANVVLDDNTITSHIKRIRRKFQALDADFGSIDTAYGVGYRWKG

>WP_082785960.1

ANIEVRLAKGVDKSTIASPYHQIKLDEPHHGIINVSLTNSVVANRDFVLQWRAKQGMSPM

ALVFNQQGKTHGDGASEDNVSENRQSDDHYSLVMVLPPKTDEHALSTLPRELILVIDTSG

SMAGDSIVQAKSALLYALNGLKAEDSFNIIEFNSELTQLSPTSLPANQTHLARARQFIHR

LQADGGTEMALALNAALPRGINRLSESSQSLRQVIFMTDGSVGNEQALFDLIRYQIGESR

LFTVGIGSAPNSHFMQRAAELGRGTFTYIGNVDEVEQKISLLLSKIQYPVLTDINVRFDD

GGVPDYWPSPIPDLYRGEPVVVSLKRSEREPQELVISGRQGHKNWQQSLSLKDSHGGAIT

EPDAGLDLLWARKQIAALELSKNGANDDKVKQQVTALSMNYHLVSPYTSLVAVDLTPIDS

SAMTRDAVVRQHLPLGWKPFGVLPQTATSSRFDMLLGAVTLILALLLAGSMLRQRRKERA

VILAIPYKQTL

>WP_007229132.1

MPDLMARALLPLLILLALQQLGSAGLIKAKAGLAPLMLAKAWEQSLASQGRPVKPWPWAD

TWPVAKLQVPSMGISQFVLAGDTGNALAFGPGHNLASAALGAAGPAMIGGHRDTHFQFLQ

HLRKGQRIVLQLPDGLLRHYRVKQMTVDTASGDMLWPNVGEQLLLVTCYPFDAFVTGGSE

RFVVTAEPESLPLQTLDALGGEPQRILL

>WP_007230524.1

MEDNIFQLQYALDTFYFLICGALVMWMAAGFAMLEAGLVRAKNTTEILLKNVALYAVSCT

MYMICGYMIMYGGDLFLSSITGDGVAGAEEAATYAPSADFFFQVVFVATAMSIVSGAVAE

RMKLWAFLAFAVVMTGFIYPMEGSWTWGGNAVFGMYTLGDLGFSDFAGSGIVHMAGAAAA

LAGVILLGARKGKYGPQGQINAIPGANLPLATLGTFILWMGWFGFNGGSVLATASVESAN

SVAVVFMNTNAAAAGGLIAALLVAKIMFGKADLTMALNGALAGLVAITAEPSTPTALQST

LFGGIGGALVVFSIVTLDKLKIDDPVGAISVHGVVGLLGLLLVPLTNDGSSFSGQLIGAV

TIFGWVFVTSLIVWGVLKAVMGIRVSEEEEYEGVDLAECGMEAYPEFTTK

>WP_007224114.1

MENEIFQLQYALDTFYFLICGALVMWMAAGFSMLEAGLVRSKNTTEILTKNVALYSISCI

MYMVVGYSIMYGGGDLTFFLDGIVGDGVTGAEEPATYAPSADFFFQVVFVATAMSIVSGA

VAERMKLWAFLAFAVVMTGVIYPMEGAWTWGGEAVFGMYTLGDLGFSDFAGSGIVHLAGA

SAALAGVIMLGARKGKYGPQGQTNAIPGANLPLATLGTFILWMGWFGFNGGSVLATASVE

SANSVAVVFMNTNAAAAGGLVAALIVARVLFGKADLTMALNGALAGLVAITAEPSTPTAL

QATLFGAFGGVLVVFSILSLDKLKIDDPVGAISVHGVVGLLGLLLVPITNGENSSFSGQL

IGAATIFVWVFGTSLIVWGVIKALVGIRVTEEEEYEGVDLSECGMEAYPEFITSK

>WP_007225392.1

MTSWDIAAIAAVLLVGVPHGGFDGAVARRLGWSKGIGGWLGFHLGYLALAAGVVWLWVQW

PVVSLAIFLAISALHFGHSDIADVPAPTTGSPSNRWLPLIAHGGLVSIAIPSLQPLAVQP

IFALLVGDEGAVMLLQAIRTLFLPWLLSFAGYAIYAVINPVWRKSLNSLIILLIVVFLMP

PLISFALYFCLWHSRGHTLRTWHRISAGSERRRSAIEAIIYSVMAWTAALVFFLYAEASL

SASLLQLTFIGLAALTVPHMLLVDLADKLNPQRPLP

>WP_140969888.1

MLGKLGELNEGYNVLTEMNGQCSDMLMDIGIYKMSNGKEELLFDNKNETAVLLLEGTIRL

EWEGMEQVIQRQSVFEENPWCLHVSKNVKVTITALSDSEVLVQKTDNDQEFASKLYTPNE

CQSVVAGDGVWEGTAQRVIRTIFDYNNAAYSNMVVGEVISYPGRWSSYPPHHHDQPEVYY

YRFNKPQGFGCAMVGEDAYRVVHNSFITIPGELDHPQATAPGYAMYFCWMIRHLENNPWN

DRIMEEDHKWLLEPNAKIWPEKE

>WP_140969579.1

MTEKMTRMTQFVKEEIANAITHGIGAILSIPALIILIIHASKHGTASAVVAFTVYGVSMF

LLYLFSTLLHSIHHPKVEKLFTILDHSAIYLLIAGTYTPFLLITLRGTLGWTLLAIIWTL

AIGGIVFKIFFVRRFIKASTLCYIIMGWLIIVAIKPLYENLTGHGFSLLLAGGILYSVGA

IFFLWEKLPFNHAIWHLFVLGGSTMMFFCVLFYVLPTA

>WP_040822867.1

MDKEMKILIVDDFSTMRRIIKNLLRDLGFTNMAEADDGSTALPMLRNGDFDFVVTDWNMP

GMSGFDLLKAVRADEKLKTLPVLMVTAEAKRDQIIAAAQAGVNGYIVKPFTAAVLKEKID

RIFERVGN

>WP_007224228.1

MARVLIVDDSPTETHRMTKILDKHGYEVITADSGEDGVAKAKETLPDVVLMDIVMPGLNG

FQATRQLSKNASTSHIPVIIVTTKDQETDRLWGQRQGAKGYLTKPIEDSALLNTISDVLG

>WP_007224227.1

MEGNFENLKVMVIDDSKTIRRTAETLLKKVGCEVITATDGFDALAKIADTKPNIIFVDIM

MPRLDGYQTCALIKNNSEFKQIPVIMLSSKDGLFDKAKGRIVGSDEYLTKPFSKNELIGA

IEAHVG

>WP_040541578.1

MALEYILPGTVSQESPAVMHTLEFKEEPGRVLLVLPTWVGDVVMATPFVRALFMRFPDAE

ITLLMNHHLYPLLEGSPWVQHCEFWAPRKKTAEAKQQQRELLNRLKARRFDLAVMLPNSL

RSAWLCFRAGAKRRVGFSRDGRGLLLTDKVEVPNRVAGGYQPLPLCDYFAVLGDALGMEH

PGDRLALFLTDQANDAVQSRLLKDGVLPEQPLVVLCPGANFGASKCWDPKRFAAVADRLV

NRHNAAIAISPGPGEEPLAEAIRDNMDAPSFLLTQPCLTLGELKSLIVRADLLLGNDTGP

RHFGRALDTPRVTVFGPTEQRWTETSHGDETIVNVDVPCGPCHKKVCPLDEQVCMTQVTV

EMVSEACEEQLSASC

>WP_169253931.1

MIAASALVLSACGSGEGGESGSDYLACMVSDSGGWDDQSFNQSGREGMENAKKNLGIEEK

LAESQGDADFGPNVDNMVQQGCNLTFGVGFLLEDTIQEAAEANPDLNFALIDSTFSDADG

KPVTIDNAKAVVFNTAEAAYLAGYVAAATSESGKVGTFGGIQIPSVTIFMDGFADGVDKF

NEDNKKDVKLLGWNKEKQDGSFSGDFENQGQGQELTKQLISQGADVIMPVAGPVGLGAAA

AAKEAGDVNLVWVDSDGYESTEYGDIILTSVVKQISQAVEDTIKEGTEDNFSNEPYVGTL

ENEGVGLAPYHDFEDKVPEDVKKDVEELKKQIIDGSLVVESESTPK

>WP_169253893.1

MNMLAMRDHSSDELRKKLLKRDLMPEAIDVLIEKLKNSRLLNDEEFAHRFARAQRENRKL

SRSVLKRELSKKGISPELASEAVADIDGEEELAREVAEKKAASTRRLDYAVRERRILGML

ARRGFPSAICIKVTRDVLTDD

>WP_169253863.1

MIPLALAALLIFFLFNSRRKQKARAEQIKSGLVPGATVMTTFGVFGTVLSIDEENNQVTI

ESGPGTVLRVHRQAIGQIENNQAAAPVDAPGADAPAADVDAAADDEKPAITDAELDAMNE

RKRAEKDTADDETAEDISADESAAKTEDADAVAEAEDVDIDAAETDADSAAADDDSTDST

DSDSDKKN

>WP_169253834.1

MGLDDHIFNRLLKERIIWLGSEVRDDNANAICAQMMLLAAEDPDADISLYINSPGGSVTA

GMAIYDTMQYIKPDVSTVAMGMAASMGQFLLSSGTPGKRYATPHARILMHQPLGGIGGTA

TDIKIQAELILHMKKQMAELTAQQTGKSLEQILKDNDRDHWFTAEEALEYGFIDKMVTRA

SDVENN

>WP_169252912.1

MKHLITGRRVTWAAIGMVGALALSGCGGAGGSSAGGGGDEVNVLMVNNPQMQDLQKLTAD

HFTKDTGIKVNYTVLPENEVRAKIGQEFAAQAGNYDVASLSNYEIPTYANNKWLTPMDEG

VADAEGFDQDDILAPMAESMTVDDQIYGEPFYGEGSFLMYRTDIFDKAGVEMPEKPTWDE

VAKLAAKVDGAEKGTKGICLRGLPGWGEMFAPLTTVVNTFGGTWFDEDWNAQVDSKEFKE

ATNFYVDLIKDHGESGAPQAGYTECLTNLQQGKVAMWYDSTAATGTLEADDSPVKGKIGY

VAAPVKETDSSSWLYTWAWGIQAAGKNQDAAKQFVAWASSKDYEETVAEELGWQHVPAGK

RTSTYENPEYQKAAEPFYQQTEDAINSADPESPGVQPRPTLGVQFVTIPEFADLATGISE

DVSSAIAGRTTADKALEKGQKEAEKVGDKYKK

>WP_169252886.1

MSLLGEPLSILALGLFNGFALCVFVLSLPPLRRPTLSSRIAPYLRDQESLVDIYAPPTPR

ADGFWGLAKSWLVSSTLWVTSRITTDATLRLRIDRLGGNATIERFRISQVLSILVGMIVA

GGLAGALSAQRGFSPIVTVVLIISGGVAGHVFNDWRLSQAIARHESRVLAEFPTVAELLA

LSITAGEGIVEALERVCRTCSGDLIDELRAALAATRTGTPLVEALDTMATRIAIPEIVQF

VDGLAVSMARGTPLAEVLRSQAADVREQSRRRLLELSGRKEIGMLVPVVVFVLPVTVIFA

VFPSLTVLDLSP

>WP_169252885.1

MSYSQFALLSGACLGAGLFLIWTSLWQRPQSKRTQSRWVRELDDMLTGAGFPRLRPAHLL

LISVAAFCIVTVVSTVLTGSWAIALCFGLFASWLPHRALQHRARSRQVMRRELWPETLDH

LNSGVRAGLSLPEALSSLAHRGPEPLRPLFEVFAEEYRASGSFALALERFRQVSADPVAD

RIVVALSVTRQVGGSDLGTMLRALAQFVRDDARTRNELSARQQWTVNGARLAVAAPWVVL

AFLSTRPETAVAYNSRTGLILLAAGFVVSLLAYQAMKRIGRLPAEPRVIEGSSLSAAHRR

FADNAGMSSGFTADSDDSADGRAA

>WP_169252597.1

MVTSAPRPPKLSLGDLVPGYLGDFGPEEFLEGMEFTDLDLAEADATQATFLDCRMTNVNF

GDAEAQIDLSGVRISGTEITDCRADTWTIPRGNLLHTDVSGTRIGAGVAYDSVWEKVRFT

NCRISYLNLRESKLTDVEFRDCKIDEIDLDRAKASRVAFPGSSVGVFQCEGATLGNVDIR

GLEPHKISGVHSLRGAIIDDTQLMLFAELFASELGISVE

>WP_169252584.1

MSTFTKVLDRWLTVTTAAAIIVMMLHTVTHALARSIFHAPIYGTNEIVEYWYLPIVALLG

IPAAQLQKEHITVTMAIERAKPATAALFTVFACILGALVSAAFAWFGLMKALENTAIGST

ADVSAVITWPVYYLVPIVFVLLVVLYILDAVAILRRRRTETGEEQ

>WP_169252403.1

MNWKSLVNRAARIGVREGLRYLRQSQSKKNSKGASQPTDARRPDTARSGGGGAASAGTAD

RSASSGQGGSGSYPGDYTGSITVSYSPDLDGDADPGEVVWGWVPFEEDHTQGKDRPSLIV

GRDGRWVLALMLTSKDHIPGGVGEVRQDRHATWMNIGTGDWDSQGRPSELRLDRIIRLDP

DSIRREGAIMPRDVFDRVAEHISG

>WP_169252333.1

MDILSSTLAAGGNAAAASESPIPVWFMITSGIIVVAILVFDLLLVVKRPHTPSMREASIW

VAFYVALALVFAGALFAIGDAQHGSEFLTGWLLEYSLSIDNLFVFIIIMGSFSVPRKYQQ

EVLMVGIIIAIVFRGIFILAGAAIISAFVEVFFIFGIFLLIVAYRQAFSSEDEGDGENGL

IRFLRKRINVVDEYHGNKLRVTLDDNKKYWTPMFIVFIAIGSTDVMFALDSIPAIFGVTQ

NAFLVFTANVFALMGLRQLYFLLGGLVDKLVYLHYGIAAILGFIGVKLIIHALHSSDWEF

LSWGHSIPEVPTWLSLTFIVFAMVVATLASLMKMKKDGISFRDMSSEESEGA

>WP_169252137.1

MNFMPGSTASNMPQSRYVMPQFEERTPYGFKRQDPYTKLFDDRVVFLGAQVDDTSADDVM

AQLLVLESQDPDRDITLYINSPGGSFTAMTAIYDTMQYVKPEIQTVCLGQAASAAAVLLA

AGTPGKRLALPNARVLIHQPAMQGQGQGQASDLEIQAAEVLRMREWLEATLAKHSNKEAT

QISNDIERDLFLTAAQAKDYGLVDQVLSSRKDAN

>WP_169251947.1

MNASLTDSSPAVPDGSGPVGSAGADDLPPADDHRIARRRSALRAELTKLAGQVKPKLRGW

FHAGAFPLSMIGGLALVIISPTIESRIAAAIFAVTGMLLFGTSAVYHRGRWRTRARLVLR

RLDHANIFLITAGTYTPLAVLMLTTDQAILLLSVLWGAAALGVAFRTIFTTAPRWLFVPI

YVGFGVAGVGYIPQIWATNFAVGLLVVLGGVCYIAGAVIYGIKRPNPSPKWLGFHEIFHI

LTILGYGCHLAALIIAAVAAY

>WP_169251583.1

MIVLTAALIAAGIVLAAPSDPGARLRELLHRAEDGESVPKGVRAVFRRGGHTADDQAERL

WAIAAVENCAHLLKVGMTPQAVMTTLSRHNDALAPISRAISLGEEPGRAIATRSSALSEA

AAHVLTGMAAVWTVSERSGAPAAEMILRYAAAQRDALDAERERRIAMAGPRSTVRVLSWL

PLIGVGLGLLIGVRPLELISGLPGQLSIGAGLLLYGAGRWWMRTMMLRAQR

>WP_169251255.1

MNGKQARTGRGRIAWTAAALALLAIAAVLITAGLLTDHSPPQPDADSAAEQSESRTTPSA

DAQPTDPSTPRTEQSSDGDQQAGMEASAPTRLRIPAIDVDTSLMDLGLTADDELEVPPLG

KDAPAGWYKRSPTPGEVGPSLIVGHVDSASEGPAVFYRLGALEPGDTVSVTREDGSEAEF

TIDDVTDYGKDSFPDYRVYGNTEDPEIRLITCGGEFDDDTGHYEDNIVVTGHLSDGG

>WP_169251032.1

MNIIDKYLSKVENVLAGGCLIAATALAVFAVLLRNITGDVLFWSEEAVIYLIIFSTFFGA

VVALRHNDHVAVDIMPTLLKGKAKKFFVVLGGLATLVYAGFIAYLSWALITEPFSRTTIT

PALKLPLWVVELSLAVGMTLFFIRAAEMTVRALRTPAAELDKDVLAEEAAAVGIAVEDIA

IVEDGRDRANRADGNDGGDRLGDGETDEKGDDR

>WP_169251018.1

MTRTLRRPRTLAAAAGIAALTLLATACGGGGGGGGEEGEQSADFITIATGGSSGVYYQVG

ATMSEILADELGADTSVQSTGASVENLTLIQDGGAELAFTQGDAVDQALAGEGAFDGKQI

DSLPVVANLYDQYVQLVTIEGSGIDSIEDIKGKSVSVGDQNSGVELNARTVVDAYGLSYE

DFSADYLPYAEAIDQMRNGQLDAAFVTSGLPNSAVTDLATSDDVKVVPFTGDGREKLLSE

HDYFGEGEIPAGTYGDSKAAETLTIPNLLVVSPSLSDDAVYDITKTLFDSIDKIQSSHNA

AKDITVDNAQDVPVSELAPGAKQYFDEQG

>WP_169250942.1

MRVIRFFEDWVVIAAFMVIVIVTFVNVLSRYIFKASLAFSEEITINLLVVLTMMGAVVGI

RLGAHLGFTYLVENTKAGTRRALLFTGTGLIVVFLAVLLIWGGEMTIAQGVRGRATPSLG

IPQWLFTLSIPLAGLLGIIRSIQALRTALGEDTSAEAVVDRLAGEATPTVDSSEFSADVK

GGRK

>WP_169250805.1

MSTSHSADQQVGHSAGRSADDSAAAGDQPLPSRRELRRREAEAAAAQGEPTAVYSDAPPV

YEQPQPTHAQPTHAQTTQAMPAVPQGHGGQSTQGRLGEDQPQNRPVTRRRRRRQPAEKEP

LGPIRGTVRTFGELCITAGLVLILFVVWQLWWTDIQANRDNEVLADELTQDWANQDPNEL

PDDPDEPVVAEPVGKNEAFGIFYIPRFGDDYYRTVAEGVDLEPVLNRMGVGRYPNSAMPG

EVGNFSVAGHRVTYGKPFNQIANLRPGDEIIVQTKDGFYTYTFRNFDIILPDAVEVLSPV

PDAPDFKGKDRILTMTACNPMFSARERYVAYAELTDWTPAGNGAPDSIKDSKAYDKVSKN

GGA

>WP_169253925.1

MRARTTQIAALAAVGVLTLSACSVTAPEALEEERLGCLVSAPAGFDDHSAGALTLEETEL

ARGAGVFSGTSSQRVSGGSATSAALDRMHGHDCALTTVIGPGGADELADFAAAHPDDLFL

GVSPGTDDLPKNVLSMDFDLVPPAYIAGFTAATASETGKVGIVVSHGFPQADRILAAFDA

GVDLYNKEKDEDLPKAESYHPSSDRAASANGSPRAIDDTRDAGKDYFERSFDADVDVLVP

FGSAAAMGVVTSADEKRTDLTAATEDPESGDDGPSLPKVIWYGTAGGFSKAIVATVEPNV

RRGLRTMFPDWPQSRNPDEVAPAPKTEPEEIGGFEVSERRYEGTIDNGGVRIVAEDGFLS

RVSDAGRGITDLRERIKSGEIDPEKG

>WP_169252724.1

MRQVEPSVELLAKPDIDWEAMRTYLDEVGGTSWADRVEDAESPDAEDLLEFSGRMCYRSW

EPGLNPNVSRVRTDSAQYLGNVVKSQHGSVLEHANFTFVLHNVSRVLTHELIRHRAGSAF

SQESLRYVRLTDIPFWFPEWAREDPELMERSLQVLDTLEEHQKWMAEHFELDEEGTKFSH

KKHMTSFMRRFAPEGLATGIVYTANLRSLRHVIEMRTAKGAEEEIRLIFNKIGEVMREEA

PAVFADYEVVDGEWIPGTRKA

>WP_169252371.1

MSPITDEFSSDDGLTRAEVRAKKRRRMLRRRRTTTIVIICILVFGIGGFFGVRAAGGVFD

DLFGPKGDYEGQGTNEVSIEIAPGSSARTVANQLVEAGVIMNSEPFLDEIERREATIQAG

TWTMREKMSSEAAVEALINPIAPPKITVAEGKQVEEIKAIMIESGMNADEVDKAIDDKTP

KDYGLDIEAPSLEGYLYPATYDLNKQKTAEDIVQEMVDKTETELDELGIESKDANRILTL

ASLVEKESPGDPEVRSKVARTFLNRISEKSKTGGLLQSDATVAYIHGARSDLTTTKKERQ

SDSPYNTYKKKGLPPGPINSPSRGAVEAALEPADGDWQFFVATNPDTGETKFADNYEDHK

KNVEIYRKWLREHRKDNG

>WP_169252360.1

MRIVLGVAGGIAAYKAAHIIRRLRELDHSVKVVPTANALKFIGAPTLEALSGQTVTTDVF

DEIDTVNHVRIGQDAELVIIAPATADLIAKIAAGRADDLLTASVLTTTAEVVVAPAMHTE

MWLNPATVANIATLRSRGIHVLDPAVGRLTGPDSGPGRLPEPEEIVDFALSVKDGTADDS

ADALNRPSGALSGRRIVISAGGTREPLDPVRFLGNRSSGKQGIALAKAAHAAGASVELVA

ANIDTGLLSGLPADITVTPVESTLELQEAMHTAQARADAIIMAAAVADYRPAETADSKMK

KSGDDGLTLRLTQNPDILRGLVAERSGQTGLRRQIIVGFAAETGDSDTTALDYARAKFER

KGVDLLVFNDVSDDRAFGHDDTMVQIISSDRGDIVVGEEFHGSKDHVSQKVIAAVSDQIT

HVESTT

>WP_169252217.1

MFSSAEELVNFISDNDVKFVDVRFCDLPGVVQHFNLPAASYGTEEITEGLLFDGSSITGF

QGIHESDMKLLADVTSAYIDPFREAKTLVVTHSIVDPFTDEPYSRDPRQVAAKAEAYLQS

TGIADTAFYGAEAEFYLFDSIQYENTPGNSFYRIDSEEAAWNTGADEAGGNLGYKTPHKS

GYFPVSPQDHFADIRDEMSLTLEQIGFEMERAHHEVGTAGQQEINYKFNTLQHAADQLLD

FKYVIKNTAFANGKSATFMPKPMFDDNGSGMHCHQSLWKNGEPLFYDENGYGGLSDLARW

YIGGLIEHAGAVLAFTNPTINSYRRLVPGYEAPVNLVYSARNRSAAIRIPVTGSSPKAKR

LEFRVPDPSSNPYLAFSAQLMAGLDGIRNRIEPPEPIDKDLYELPPEEAKDIKLVPGTLD

EALIELEKDHDFLTAGDVFTPDLIETWIRIKRENELDVARLRPTPTEFELYYAL

>WP_169251582.1

MSEWLDASLVADVRKTLLDKPGPVTTAAVAEAVQRTGRVLGSSALLELVTRLSAQLSGAG

PLQSVLEVPGTTDVFVNGPREIFADTGGGPRLLDLSLGSEEEVRSLAVRLAALGGRRLDD

SSPYVDVRLPDGVRMNAIVPPISGDTTTISFRVPKRSGFLYSQLCDSGFVPDEISPLIAE

AVTSRANILISGGTGTGKTVLLGALLSLVETIQRIVIVEDSRELIVTHPHTVQLAARQAN

VEGGGEVTLTDLVRNALRMRPDRLVVGECRGAEVRDMLTALNTGHEGGCATIHANTAEAV

PSRVAALGALAQMSPDAVYSQFSTAIDLVIHLRRCGSERGITELAIPTRAGAEAVVMEPV

WGRPSPSTTGTVNRRALARFEAVLEQKSAQ

>WP_169251567.1

MSVNIGVVGATGQVGGVMLDLLADDPGFEIGSLRLFASARSAGKTIDFKGQPITIEDAAE

ADPSGLEIALFSAGGATSKAQAERFAAAGVTVVDNSSAWRSDPEVPLVVSEVNPDDLDEP

PKGIIANPNCTTMAAMPVLKALHDKAGLTRLIVATYQAVSGSGLSGVEELAGQLEAGLPD

ARKLATDGTAVNLPEPNNYVEPIAFNVLPMAGSVVDDGQNETDEEKKLRNESRKILGLPD

LLVAGTCVRVPVFTGHSLSIHAEFDSDISPEQAAEILSQAPGVSLDEVPTPLKAAGQNAS

FVGRIRADQSAPVGKGLVLFVSNDNLRKGAALNTVQIAALLAAKLEAKAA

>WP_157361095.1

MGDPVSGGGGWRHADRVVAVESVVSGTVECMARKTLTAAPRLEKLRLEAIAVGDADHLQA

HESYSGGRYVASDLRERELSGISFSECEFVELEASETDLRAATFVDTRFERLNAPIFTAP

RSSFRDVSFEGSRLGSAEFYEANWSSVHFVHCRIGYLNLRGARLEDVLFTDCLIDELDLG

AATANRVSFIDTQINNLDLTRSTLTNFDLRGVELRQLGGVEYLKGATLNSYQLSELAPLF

AHHLGIVLDE

>WP_148224521.1

MGNTTFSDGAIKGLAAALVVCLHFACASLASAEEDASSPGPREVVAAATDNIMALAREAP

AYFDTDPDRYTVAVGEELDRVVDFRGFARGVMGRFASKELYQGLDEAGRNQLREHLMKFT

EVLRSGMVNTYSRGLLAFGGSQVELGEVDMAPGSTRVASVTQRVFGDDGKIYTVKYQMGQ

YRDGRWKLRNLIIENINLGEIYRGQFEAAALEAGGDLNTVIANWDDNRVKSLSTEE

>WP_007228891.1

MYYGEKLNSISHLVGAVIALIALGALLALGIQTGDPLVIVGFTAFGLALVLLYTMSTLYH

SFQEPRLKKAFQLLDHISIYLLIAGSYTPFMLVSLGGSEGKMILSLVWTLAIIGILSEVF

LSGRVVKTIQLVIYLSMGWACSLEFASLKAALPEVGFFWLTAGGIAYTAGIVFYLLDKMN

RLDHSHGIWHFFVLVGSVCHLIAVIGYVR

>WP_040823262.1

MVDVALFPVPNSVNFPGVPCSLHVFEPRYRQMVRQCIDQNLLMGVCHTEKVLHRKEREQT

LEEALNSNQTTYKPRGIFSAGPVELLEELEDGRMLIQVNNEVRLQLGEEKQTLPFGIWAC

EELVDEALDETGELALNQSQSKILQRLLAVTHGNEDAQDMLNSIHWRSMPAQTFSFAVAG

LLGMPPETSQALLEMTSAQMRLDTVLEMINTMGTALS

>WP_009774169.1

MSIQHKLPRSSGSALTILMVLALAIAGCSVNSSPRSEASNASAIPPGAFSATVNYVHDGD

TLYLDTGREELKVRLIGIDTPELAGQQRPDAEECYGAEARALLRDFLPEGTQVWALEDRE

PEDRFGRSLLYVYLDDGTFVNLAMIELGAAEAIKVGLNDQYWPELRDAEDAAHSAQLGMW

GAC

>WP_009774019.1

MKRARSQLPEVASVTQRLAVLLNAGLTPSSAWFHVARGRSADGVAALVALAGEEPGGAAE

RIVRAAEGLAPLDRQAWNSLAAAWFVAGQVGAPLAAALRTHARALRMLVHVQREVATALA

APVATARMVLALPAIGLVFGALLGFDTIGVLVSTPIGWGCLVVGGALIAAAVRWNRRLVR

SATPTQAAPGLECELMAIAVSGGGSLVNARVVVAGALERFGLMGDGDHLEGVLRLSHEAG

VPAAELLRAEADELRLAADADARAGAAALSVRLMLPLGLCVLPAFMVLGVLPLMVAVISS

TVAGL

>WP_009771703.1

MSLILGIILAVGVTLVAAPFLWPAQGERRVRGRSAWSLRLRERLVQAGLPTTSPSIVLIV

SVVFSVAVAAVTFVLTSVIVVVLCSAALALVLPTLAISWRARARRTATQIVWPDVVDQLV

SAVRSGLALPDSLMTLSQTGPLVTRSAFAAFAARYRATGNFSIAVDELKVALADPVADRI

LETLRMSREVGGSELTNVLRSLSVYLRQEAAIRSEIEARQSWVMNAARLGLAAPWVVLFL

LTTRPEAAAAYNSASGVALIVAGLILSLVAYRIMAGIGRLPQQPRWFA

>WP_007234989.1

MTEIPLFPLSSALVPYGYMPLQIFEQRYLDMVAACMRTGTGFGVVWLREGSEISGGSHNT

PDVGKYGTHARITDFDQLPNGLLGITIRGEERFDIAEVWRDSSGLIRAKVSMEAPLAPAS

MTDEWRSLEIVLRGLESHPHIQRMNLTIDYNNAWEVAFTLIQLLPFDEAIKYELLGLSTL

DELIVELDILLNQISGEDG

>WP_007234840.1

MFVCICNGVTDSSIRREMEAGATSFADVQNRLGVARQCGSCEHLARAIVNEFSKPDPRYF

YNACSDESMAAVA

>WP_007234584.1

MTEDKEYSRMRWASRRGLLELDLLLEPFMEACFRALEPQLRDDYQQLMGHEDQDILNWIM

GREALADETLTAITEQIRVHNRSKLR

>WP_007230552.1

MNQTLLIAQVAAFIATSCTVLFIGFSIRDALKRREQLWIKIRASHVSKAEAGAEFVRTVL

LETVRPMSSVQKLLRQTQKNLLRAGVGRNAEDYVADGLWQGLLGGISIMLAVSLLSSPVS

GLLLGTAGGVLWAMWIKPSMLDSNATQRSRLIYRRIPYALDLSVLVLETGGTLREGLEEI

SLQNDPLAEELRITLLEMDSGSTQAAALRGMGQRVGLESLETILTAINRGEETGAPMVAT

LITQAEMFRERRLQEIEKMAVEAPTKMTFPNMMIMVSVLLLIIGPLLIQLVSSGLF

>WP_007230536.1

MGLIIAYALIAVSVTALVYAGLQSFVPVAVTRWRSDQEEIDDKLQNIFYTSSEARTFLIL

KYGGTLAAFFIGLWFMNSLVFGIFLGIVIYLLPEVLLDNILRRRRERLEAQTADVMTALS

ASIKSGMTIEQAFSEMVDNMYPPISEEFALIRERIDAGQPVIAAVKSADERLQVPRLSLI

FQTICISLERGGRLASLMDRLAESTREIERVEERVRTETAGLRLSARIMFLMPFFICGLL

YLIEPDQVMLLFDNLVGNIVLVIAIAMDISAYFIMKKLIELDI

>WP_007227341.1

MAPSAIQSLDALEQRFREDAVEAPVQDLERMEWVGTCLSIAGVPLLIGEGELEEIIETPN

VMAIPGTKPWVQGVGSHMGGLIPIISGDVFFRKRPYSGRVRDYCMVLRRPGFYFGITLSG

LERDMKFLVETRDMTVTTDPDFAEYTLGGFPDQDRVLAVLDIDKLIADSDLSNAAANDPD

SPEERTND

>WP_007226034.1

METIPLFPMHAVLFPHGRMFLQVFESRYLDLIGQCMKEDSGFGLVWLKQGQEVYRSNELV

DPQLAQIGTYAKIVDWDSLPSGLLGVTIEGSDRFRLLTSYQRKDHVHMGEVEWIETAGAT

ELPENYAELWGLLQTLLDHPHVDRLKLNPVVNDVNAVSCLLAQLLPIEERVKFNLLAAAE

PLDRMARIMTLLDQYSE

>WP_007224867.1

MQKIRVYWRDIPSQVIIKRGRLRAKAPLTQRFQVAIDRAAMTAGRGSSEAYVADWRRETT

SISGDGDLGQLAESEALMLENQYDDDRLKQLVANRGLEAE

>WP_007235031.1

MKHSGILVVDDESVIAEELCEFLSSFDYTCQKALSVNEALALIETNLHITLILTDMRMPG

RDGAELIQALQEMPGRQFEYLMISGHLDADEDLKHINNEGVTLMRKPIDIDALLLFLEER

EFTAVPNEN

>WP_007234880.1

MQKEMDTSSLKVLVVDDEAFVLKLTVRILSKLGYDNVVTADNGVVALGEIDNVTTPFDVI

ICDLNMPEMDGIAFMRHAADRNVSAGMILLSGEDERMLETARDLAAAHKLHILGVIPKPL

KPDALSNLLNTFQPTAVVEKQGWHQEGISETELLDGMNSDQLHLVYQPKVNISTGEVTGV

ETLARWMHPEKGLLGPGAFIPLAEETGHIDQLTCAIYRKAMHQAGDWLAQGITMKISVNI

SVNSFTAPGFTDFLIETAQNEGMDLSNVVLEVTETQVMDNALGILETLMRLRMKRFALSI

DDFGTGNASMEQLKRIPFSELKIDRAFVFGAAENAGARAILESSVTLAKSLKMDIVAEGA

EGREDWDLLASLGVDTVQGFYCAKPMNNADLMSFLEDWTGPH

>WP_007235808.1

MVKHSNTEDPWALKVLAAVAELGPQRYYDPVITSEVVKNDLVSLIRASQKLKEKVAIASP

EARHDRRNIIGAIRGYSEMLLEDSEVLPAAVRAHLLQILAAAKNEPKPASESATPTKSVT

LLPSEEPGVILAVDDLPENRELVSRLLQKTGHTVISAESGEEALELLDTMGVDVVLLDLV

MPGIGGAEVLKRLKEDERLRATPVVMISGQQDMDQIVMCIEAGADDYLLKPFNPVLLQAR

ISAGIERKRWHDREELYREQLERREQFIRATFGRYLSDDIVDEILERPEGLELGGDLREV

TIMMSDIRGFTTLVEHLPPQQVVTLLNRYLGRMTEIILEFGGTIDEFLGDAVLAVFGAPR

RNDDDPDRAVRCALVMQEAMADINIANSADGLPEVEMAIALNTGSVVAGNIGSERRSKYG

FVGHAMNVTSRIEDVAKPGEILISQTTYEKLESDYKFGNSRSLSVKGIEAELRVHAVLGG

IQ

>WP_007225300.1

MDTAEQQLLLIDNDEVERKSVAAYLKGAGFIVLEASNVSQGLDILADHQPEVVLCDLNAT

GTDSGPLQAIKSDFADTPVIVMATDGVMSDVVWALRYGAADYLIKPIADMEVLEHAISRC

QEQRQLRQQNLDYRQKLEQANQGLQESFKVLELDQLAGREVQLKMLPPTVKQFGKYQFSH

RIIPSFYLSGDFIDYFTVGDDFVVYFIADVSGHGASSAFVTVLLKNMFARKRSDYLHQND

ASILSPAAMLDIANRNLLTTDIGKHATLCVGVIDLRTDTLSYSVAGHLPLPMLTVDGEPQ

YLQTEGMPVGLFEAAEYTEATISLPPSTVLTLFSDGILEAISVKGVLGQERFLLEQLSRG

PNSIDGVLEALKLNEIGEVPDDIAVLLITKDVHLGDSIIHADTGSDSE

>WP_050756508.1

MDLILASTSPYRRQLLERLQIPFRCESPNVDETAHPGESPAALAQRLAAAKALDIASTNP

GAFVIGSDQVASLSGSCIGKPGSHAAASKQLHDSAGQRVDFYTGLSLINLSIDYHETLIE

RFSVVFRELESLEIETYLQKEKPYDCAGSFKCEGLGIALFEKMIGDDPTTLVGLPLIATC

RLLKAAGAPVLEH

>WP_009772888.1

MRLYLASTSPARLATLRAAGVDPVLLASGVDEDAVAAAAPGPLDGPALVELLARAKAEAV

VGSRINNEPIDGFILGGDSAFELDGELFGKPHEPEIARRRWHAQRGRTGVLHSGHWLIDH

RGGQLRGATGAVSSASVTFASNITDAEIDAYVATGEPLKVAGSFTIDSLGAAFIERIEGD

PHAVVGLSVSLVRQLMRELDAEWTDLWNIERPTL

>WP_040811778.1

MTMKKILVVDDEPDLRDMLRFALETEGFEVLEAADTQKAYWLITDQDPDLVLLDWMLPGG

SGIELLSRLKKEEATQSLPVIMVTAKAREEDIIQGLDMGAHDYITKPFSLKELLARIRTI

FRHTEDDDVNHQLRVGDLVLELDNRRVTLGSQVLMLGPTEFKLLQFFMLHPERAHARKQI

LQHIWGNNACVDARTVDVSIRRLRKTLQSAHPVYSELIQTVRGTGYRFSPRDLVAA

>WP_007227708.1

MSEPTVLVVEDEKAIRDMLRMALEVAKYRFIEAENIRDAHVLIVDERPDIVLLDWMLPGG

SGLELLRRLKREDNTREIPVIMLTAKAAEDNVIQGLEVGADDYVTKPFAPRELIARIQAL

LRRSAKDSAQGRIGLNGLVLNSDSRRVFAGEIALNLGPTEFNLLQFFMSHPERAYSRSQL

LDQVWGANVYLEERTIDVHIRRLRKALQTDHADYGELIQTVRGIGYRFASREHS

>WP_009773595.1

MKFQVNRDVFSEAVSFAVKLLPQRTTLPILSGVLIEATDEGLTLSSFDYEVSARTQIKAE

VDEPGRVLVSGKLLAEIASRLPNAPVRFSTEDNKITVACGTGHFTLSSMPVEEYPTLPQI

SDQVGTLKADLFSAAIAQVAVAASRDDVTPVITGVQLEVSQNNISLVATDRYRVAVRDIE

WDAGASGVESATALVPAKTLVEVGKTFGNSGEISVAITSTDERELIAFHADNKTVTSLLI

KGNFPPVRRLFPETVDNFAVMNTAELIEAVRRVSLVLEREAALRFTFTTEGVTLEAIGSE

QAQASETIDAFLTGDDTVVSLKPQFLIDGLSSVHSEFVRISFTKTENPNKPGPVLITSQS

SKDQPGSDNYKYLLQPNLLLR

>WP_007225086.1

MVETLLIPLMLIVLAVFFLLSPVMMNRSLQRSSRSGVNIEFFKSRLSELESDRARGILDD

DEFEQLKIELERRLLDEADSGHTAPSAHVKTSFKTAIMLALLIPIVAVVVYQQTGAKADW

DIAQTLKNMRLKTADGEAAETDVKQLIRQVEERLEQRPDNGSYLMLLANQQMGLRNYPAA

AAAYQRLRTIYPDDASVLAQYAQAMYLSSDRTLTTKVTDMAELALRQDPQQPTVLSMLGM

AHFEQGDYQRAIDYWQRLLPSLGPVSPNRKIIMAGIEQAKSRLGSSDTTSIDRPDVIKNA

SIQLSVSIDEGIIASSDSVVFVFARSASGPRMPLAVAKLTVADLPVILTLDDSMAMAPGL

NLSSQKEIEVVARIAKNGIANPGPGDIEGRVGPIKLEEVDGVVAIAINKTL

>WP_007225370.1

MSYVINFEAVPEELKTLPQWVCWKAVVRPNGKITKVPMNPLTGIKASSINSKTWASFDKA

ATGMNRHGYDGIGFVFVRGDGLVGVDLDNCMRSHGQLETWAQDIVDRLDSYTEVSPSGNG

VHIICYSGASGLSYNKDGREMYSEGRYFTVTGNEYYVRGHSNEN

>WP_007229947.1

MTLLNTKIHLTDNQMHQNQDILGNISTAVVSLDSELRVVSLNSSGQDLLEASEARSLGQP

MHKLVANPEALMEVLRQVRADRSPLARRGMPLMLLSGREIHADLMLTPVSNSEHGINILL

ELQPVDRLLKISREESLHHAQETTREMIRGLAHEIKNPLGGVRGAAQLLARELSSAELEE

YTNIIIREADRLRDLVDRLLGPNQQMDSQCMSIHEVLEHICNLVRAETDNRVELVRDYDP

SLPDIIGDRSQLVQAVLNIVRNALQAAPSEEDCVITLRTRPQRQFTIGNQLHRLLCRLDI

EDNGSGIPVDMLHSVFMPMVTGRAEGTGLGLTIAQSIITRHGGMLECSSEPGHTRFSIYL

PMDLNHA

>WP_040821591.1

MSTANTVWIVDDDRSIRWVLEKALNQAGITTQTFDSGETILNSLRQNTPDAIISDIRMPG

MDGLELLGKINETHPDLPVIITTAHSDLDSAVASYQGGAFEYLPKPFDIDDAVAMTERAL

LHANEKTADSTEPEENSSSTEIIGEAPAMQEVFRAIGRLSHSNITVLINGESGTGKELVA

HALHKHSPRSSQRFIALNMAAIPRELMESELFGHEKGAFTGATSLRPGRFEQADGGTLFL

DEIGDMPSETQTRLLRVLADGEFYRVGGHVSIKVDVRIIAATHQNLESLVKEGRFREDLF

HRLNVIRIHIPSLRNRREDLPKLMQHFLLKASEELNTETKLLTPESEHYLSKLDWPGNVR

QLENTCRWITVMAAGREIHLEDLPPELLDQTIADGGTDDGNWQDNLRRWADQELSLGKSN

ILDTAIPIFEKLMIDTALKHTHGRKRDAAVLLGWGRNTLTRKMNELGMNTTIIDSTES

>WP_007229945.1

MHSQAKVWVVDDDSSIRWVLERALKQAGINNESFSDADQLLKRIVSETPDVIISDIRMPG

TDGLELLSQINASHPELPVIITTAHSDLDSAVASYQKGAFEYLPKPFDLDEVVAITERAL

AQVRERSIEAPVLEELPETEIIGEAPAMQEVFRAIGRLAHSNITVLINGESGTGKELVAH

ALHRHSPRAASSFIALNMAAIPRDLMESELFGHEKGAFTGANAKRAGRFEQADGGTLFLD

EIGDMPAETQTRLLRVLADSEFYRVGGHTPVKVDVRIIAATHQDLEELVRRGDFREDLFH

RLNVIRIHIPKLSERREDIPRLMQHFFQSAAEELGGEAKILLPATERFLSNLDWPGNVRQ

LENTCRWITVMASGREVHITDLPPELSRDVVPDPQSTDSDWRAMLQNWANNELGQGKQQI

LEQATPAFERVMIEVALKHTQGRKRDAAELLGWGRNTLTRKMKDLEM

>WP_148224536.1

MPKLLIPIVVTLLGTVVLTIGLSFGALEQRKTQLAVSAQDMVSINATLSPLLHFRNQSAI

REALLLKLRLETKPQKIWLRVLHVDGEILAKAPRRDSDRGTTQFIRTFRNRLEVLTAVDL

FAHNSEAHYRGALAAIPFMDAQFKMSTPVFSLIDPLRTDVPRSAYQQTLLQKADQPLPFV

AGYIEQGIFLGDILETIIPTLWQALIMSMIISATMLLAFYYFAVRSRVSISQRTAQTERS

DQKNLPQKMESARSVPTNDETTSAPEDRLTEPSLDETRSNIGDPATLDPVTSLPDRHQLL

EHMAQGMRVAAAEHRCMGLVLIEVCSIRDILRTRGREVSDNVLREMTSRILNSIRRSDFA

SRGYDALGEAILDADQFCIVLCDLDNIQGVGSAAERLLGQLRLPVTVADEALCLNVVASA

ATAPQHSKTPEGLIIAAKSALIQARESRAPNTILFSS

>WP_148224448.1

MFAPTNLDLRGRTPPGYIAVEGPIGVGKTTLARRLAEAFNYQVLLEDAHENPFLDRFYQN

RKEAALATQLFFLFQRSQKIADLRQTDIFEPVRVSDFLIDKDPLFARINLDPDEYSLYEK

VFQQLTIDAPLPDLVIYLQASPDRLLERILSRGVSSERGIDREYLEQINEVYSEFFLYYD

AAPLLIVNANEIDLSQGDEDFSQLVNYLLDIRSGRHYFNPTFFG

>WP_148224378.1

MALLTVLVVILRYGFGVGAIAAQESVIYLHGALFMLGASCTLQAGGHVRVDVVYQRFSPR

ARAWVDALGHVIFTLPLCAMVGFASQDYVFESWVARETSPEPGGIPAVFILKTLLPVMAI

LLALQALSEIIKAVKTLISEVSHCD

>WP_040823729.1

MNDLAPSNIPYAYGGPLISGSLRDQPADFFVEENLGFEPEGEGEHVFLWIEKTDINTQQL

AGDIARLAKLPTRQVSYAGMKDRRAVTRQWFSVHLPGQDNIDWQALNSGQVRLLKQVRHL

RKLRRGAHRGNRFVIIINDVSGDTSQLASAVATIARRGVPNFFGEQRFGYGGSNLMRARQ

LFSGQFKPKKHQRGLYLSAARAYLFNQVLAQRVEANNWDQLTSGELLMLNGSHSVFAQGD

TIDLDARLLDGDIHLTGPLYGKSGSLAPTAEVAAMEADILQATPDFTAGLLQAGLKAERR

ALRLLPVDLQAQLSERQLTLSFALPTGCFATALVRELVNYTETHNHV

>WP_040811348.1

MYPLTRSLLVLLLSAFVSACATSPPSDTSNVCAIFREKSGWYNDAKKARARWDTPISVMM

AIMHQESRFVATAKPPRKKIWGIIPGPRPSDAYGYSQAKDATWEWYERSSGSYGADRDDF

GDAIDFIGWYNDMSFRQNGIAKDDTFRLYLAYHEGHGGFKRKTYRDKQWLVDVARKVDGR

ANTYNTQLKGCVKSLEDDKWWDFF

>WP_040542916.1

MIRIMLSVAVCVAQQAAADDINGVWKHADEPGWIEIQLERGSGTVERNDKFPERVGREIL

KDLATGGEAQTWSGLIYVEKMGEYKNADIILASPDRMKITVKVGFMSRTIGWQRVDEVPA

AP

>WP_040542885.1

MRILLVEDDVQLGESLEAALRLEHYAVDWLRSGEPVRATIGATPYDLMILDLGLPEVPGI

QVLRQTRADKHDIPVLLLTARNTLDDKVDGLDSGADDYLTKPFEIDELFARVRTLLRRRG

EGRSQQLEARGITIDPVDRLVIFEGEMLDLTAREYAILEILIRNAGRFVSRPRLEEGIYS

WGEEVGSNTVEVYISRLRKRFGSDCIETMRGVGYRISQ

>WP_040542483.1

MSFVHKLVQTIDAFTDRSGRVLAWLALAMALLITAIVIMRYGFNTGSIFSQELVTYMHAT

LFMLGTAYALKHGAHVRVDIFYRQFSARGKSWINALGGVVFLIPLCLFIVGVSWNFVNES

WAMRETSSELGGIAAVYLLKALIPLMGINLLLQALAETLRSTLELVEGNT

>WP_040541029.1

MNRVTVLLSAIALLIAAISVYLSFRLLDPTPPKTLILATGTAGSAYEEMGQSYRKILKES

GVEVQLLASGGALENLELLKSGQADIGFLTMGYPAGQDAVNLRSLGAMFFEPLWVFTQDN

DLLEGNLDSLRRTRISIGPSKSRSNSASRKLFELNGLEISDLNLFELDPTTAAQQLKQGT

LDTLFITGNSISPVIKQLLSSRETVLVDFKRADAYVALFPELTKLVLPAGVGDLALNMPS

SDTRLLAFTAMLGVNKDLHPITQSLVLEAAERIHAKPDLFHQAGVFPQARDQLIILSDSA

KAYYADGRPLLLRLLPYPVAVLFMQLIAAAIPLLGIAYPMFKLLPSAFHWIMRHQFYRVY

SELRQIDRSIGNITEAELKTHLERLENLEQKVTGLKVPITYSTMLYALKGHIGSVLKRVR

DALG

>WP_040541869.1

MNEALQQVINRNDTWQGHLAGQVLSANGDSHWDEDRLSTGYSTLDKELRSDGWPLGSTVE

VLSDGCGLGSMGLFLPAMEKLSAEGRWQVFIAPPFTPYAPLLKARGIDTDQILLVHPKSR

EDLLWATEQALRSTTSSAVFSWLGADEYSYSELRKLQLAAASGDSLSVLFRPQEAARNHA

PASLRLQMREYRKVHILKQRGGNQYIDVTLPPSEDVPEHPQLWEVPSWQASPGQASPGQA

SPKAQPAFSFA

>WP_009773271.1

MISHHYRRAMLPIATLGIAGMVLSGCASGSGGTGGPGDAGDPDGIVTIYGTIADTEAELL

EESWADWESENGIDIQYEASKEFEAQISIRAQGGNAPDLAIFPQPGLLADLASRDYIQPA

PAGVQANVDEFWSADWAAYATTGDTLYGAPLMASVKGFVWYSPADFADWGVEVPETWDEL

LALTQTIADKTGTAPWCAGFGSDAATGWPGTDWVEDLVLREAGPETYDKWVSHEIPFSDP

AIVSAFDSLGEILLNPEYVNAGFGDVKSINSTPFGDPARALGDGTCALHHQASFFDGFIQ

DPKNGNATVGPDADIWAFVTPSVEAGGNAVTGGGEIVGAFSNDEETIAVQEYLSSAEWAN

SRVKLGGVISANNGLDPASASSPILQQAITILQDPDTTFRFDASDLMPGVVGAGSFWTGM

VDWINGKSTEDVLSTIDASWPSE

>WP_009773251.1

MKRTATSVVAAVAAVGLTISMSACSTTSAAGSGDAEGPLTVWVMGDSGANFEMLVADSGI

EVEVVAIPWDSIDEKLTTAVASGSGPDILQIGLSKLRTFADAGALLPLDDEIANHPGIDP

ANFPAGVSGTATSVGGEIVSVPWTSDTRVLFTRTDILSEAGIDAPPATWDELRADAKTLA

ARGDDQYGYYIPQWDAPLPIEMTWSMGGEVIDADGNVNFDTPEFQKAVDVYTGLYADGSV

PVNGDFDQTQGFISGVAPMLVSGPYLGRGIADSAPELDGKWQASPLPAGDGGSISLFAGS

NLGVWFNTDQKETSLDLLEYVSQPEQQLEWYSMTGELPTVSSALEDGDLNSDPNVQVYTD

QLKTAKVLPLVSNWDGAVGTELLNALNAIVLTGADTKSSLDGLYSTTAGLTIN

>WP_009772227.1

MTDQNTNPTAPRRVVVAEDESLIRMDIVEILRDAGYEVVGEAGDGETAVALATELRPDLV

IMDVKMPQLDGISAAERLSANHIAPVVLLTAFSQKELVERASEAGALAYVVKPFTPSDLL

PAIEIALSRYAQIITLEAEVSDLVERFETRKLVDRAKGLLNEKMGLSEPDAFRWIQKASM

DRRLTMHDVSQAIIDQLSAKK

>WP_009771998.1

MHEIICPHCKKAFTIDEAGYADILKQVRDREFKTELHAQLALAEKEKIIAVELAESKIAS

GLGKEAAKKETEIEHLKAELKATDMEKQLAVKDAVSAVEKERDEAKNEREKNVVEKDAAI

ELLRAELKSTELAKQLAINEALSAVERDRDDLVRNLKATEIEQKLLESTLKEQHSTEVRI

LTETIDSYKDFKARLSTKMVGETLEQHCEIEFNRLRSAAFPDAYFEKDNDAKSGSKGDYI

FREHSASNVEIMSIMFEMKNESDMTATKRKNEDFLKELDKDRNEKGCEFAVLVSLLEPES

DLYNGGILDVSHRFPKMYVVRPQFFIPIITLLRNAALSTVQVKLELARVQEQNVDITKFE

GNLRAFKEGFSRNYSLAADQFQESIKRIDEAIKDLEKTKENLLKSSNNLRLANEKADGLT

IKSLTRGNPTMAAKFAELESPDDPENFK

>WP_007235241.1

MKILLAEDDAQVRTELRELLVEQGYDVTTAIDGIDAFEKFRADEMIEILLLDIRMPRATG

IQTLDAIKKIESANERVFETLFITGASDNNAIVSALKLGAFAFLFKPIVVEELLKELSEA

TDSINQKHYRNFQNSMFNPNLESRAPGKSGSIKEGMGATSEIAAIGAEHYAPGIEQHVHR

ISEMALCIAVRLGLETQHCQQIRLASLLHDIGKLGGPTDIYTAERALTEEEFEKTKDHTR

LGAALLEHYDDPIIEVAQNIALQHHENWDGSGYPAGLKGDEISIEAAIVHAVDTYDNLRS

HRPYRAALPHHVAMEILVSGDEKSNPDHFHPGVLQALLSQHREIEAIYERYRPIDVESKI

TDQTPA

>WP_007235034.1

MKVLVVDDESDIREELGEFVEQLDFSVVLASNGEEALGKYFDDPEISIILSDLMMPGLNG

LEMLDNINSAPGAHQRVRRVIFMTGNGNTQSVIRAMHLGAKEFLLKPVDLDQLERHIMSA

KREVATDRARQTEEILLKQQVSLNNAQISSLNRDVEEAYAEALACLAAAAEHKDPETGQH

IIRIGEYAAVLAQALGWDKEWSEMIRLAAPLHDVGKVGMRDSVLLKEGPLDDDELHHMRQ

HPETGYQILSVSNYPTMKMAARIARCHHERWDGTGYPRGLKGSEIPVEASITTLVDVYDA

LRSKRPYKPAFDHKQVMDIILNGDGRTEPKHFRPDVLAAFEAAQDKMADIFERLSDDTDS

SSHTSSSKREVL

>WP_007233471.1

MSELLIDVGNSAVKWAVCHGLNLKSQRHSGSFSDLAEAMWTSAKGDSTVWIASVRDEQSD

QVLVSELHAVGFSNVHLCGTAQKEDGLLNSYAEPSRMGADRWFAMLGARACKRGPLLVID

AGSAVTCDLVASDGRHLGGYIFPGPALMEAALQSNTQKVRYSDSLKLALDPGQSTAECVA

SGISVAMLGAIKQVCDQYPAHQVIFTGGAASGLNAVGLVGDWRPDLVLEGLLSRAHGTEV

AFAT

>WP_007230922.1

AAFHYFYMREVWVMSGDTPTDFRYIDWLLTVPLLMIEFYLILAAITKVAGGVFWRLLIGT

LVMLVPGYMGEAGYLNVTVGFGIGMLGWAFILYEIFFGEASKVAANEAPAAVQKAYTLMK

WTVTIGWSIYPLGYFFGYMAGGTDVGVLNIVYNLADVLNKIAFGLFIWYAANEDTSAKA

>WP_007230850.1

MKILLVEDDLATREEVSDLLETLGHHTVDSDCAEEALQYIRSDGSADLILFDLNMPGTSG

LDMIREVRHTSTQSISTMPAICMTGSRDAHSVVELLKTGITDFLFKPLRLADLKSSLQKV

ESEISRVRAQESQAAALNDKLNEKDKLLEELSLELSESQTESVLCLAYAAEYKDLGTGAH

LRRISKYAERMAELLGWSEERCSSIALAAPLHDVGKIGIPDTVLNKTGTLTSREFNCLKS

HTTLGAEILSASKSPVMRLGAKIAHYHHENYDGTGYPSGLVGSQIPIEAMITAIVDVYDA

LRTSRPYKEAMDHASAIDTMCNGDERTDVSKFHPELLKTFLHNHHDFGEIYRKNTVADVP

PIMLSAVAH

>WP_007230835.1

MSAEFDLKRSCFGDFMKILVVDDEPLIGSETSEYLSLHGYVSDHCHSCDAAMEILSSDPD

IRLVITDLRMPEKDGFTLIEATQPLQRHIEFIVVTGHGGKDEAINAVRTGVSEFLAKPVN

HFELLKAVKNAAQKIADHDHELSVKSSLQGKVFAGEKKIDRLLGNLDTSYAELTYCLATA

SEYKDPETGQHISRIGSYAALIAGLMGWTERKVEMIRLAAPLHDIGKIGTPDSILLKPGK

LGDVELRVMRTHSSIGHAILSQSTSPVLKMAANIALAHHERWDGSGYPNGLSAEEIPVEA

AITALADVYDALRSKRPYKPALDHRTTCDILLYGDGRTEAKHFSPELLQIFKENHDKFDE

IYESMHDQAVSC

>WP_007230577.1

LSLTGTDLEVELVGRVKLASAGDSGELTLPARKLVDICKSLPEGSEISFAAEDSKVTVKS

GRSRFTLSTLPAREFPNVEDSMGTHQFTIKQGQLKRLIDRTGFAMAQQDVRYYLNGMLWE

LKDKQLKVVATDGHRLALCTLPEKIEAGDDAQVILPRKGVLELARLLLAEDEDIAIVIGS

NHIRATTEEFTFTSKLVDGKFPDYQRVLPRSPNKIVLGSRLELRQAFTRTAILSNEKYRG

VRLKLTDNSLDIVANNPEQEEAEEAVPVDYQGESLEVGFNVSYLLDVLAVLSGEQIKLSL

SDPNSSALLEESDEGDSLYVVMPMRL

>WP_007229328.1

MDPIGTALVVDHDTARLEALASLVSSLGFTPENYTDANAAREYLSTRPDLDVILCEMDME

GLTWDSAHRSLQEMDVQIPVILFSDEAQASRMMRALRFGASDFFVRPVDDVEALQRSLDR

CVRQRQVRRELEQSRQRLQAANTELRGTIHVLEQDQQAGRQVQMRMLPATPLVLNDYVFS

HTVIPSLYLSGDFTDYFTVGDHFVTFFMADVSGHGSSSAFTTVLLKNLFARKRSDFLRQN

DDTILSPLALLKRANKEVMDLEVGKFATMVVGVLDMKSNNLRYSVAGHLPQPVLVSGDGA

RYLRGEGSAVGIMDDASYEEHMIDLPDSFMLALFSDGILEILPPKNLIEKEKYFLNVFEE

TANSPEEMVTRLGLDQADTAPDDIAALFISKRN

>WP_007228310.1

MSLCLQLDVGNSSAKWRLLEQGDVLSRGRYSAADANTQRELLESTASVDQIWVSSVAGGD

TEAELKEMLEQRWGVTPWFARTPAATGDLRNSYADPARMGVDRWLAMLGARARCGKRVCV

VDAGSALTIDLISATGQHEGGYIIPGPALMERALLLDTDRVRFTDEVSYDLAPGSSTAEA

VRHGIAVAQVGSVSIVLDGCASEPPALIFCGGAGQVLQQLLDRGGEFIPELVFEGLEIMA

AAP

>WP_007228237.1

MKNVLQHLTRIRDWWIALCTGRDPMRPLDEEHYRQRILAITSFFCLITVIGVPVVIPLVI

DISPQGRFAATTLLAIIGLSVLVSVLILRYLNNRIAALHLLLLVYTGAFAIACAYFGGTR

SPTFALLILAPVMASVVGGTGAGLFWTALVLIIWSVILGLERLGVQFTQIILPQNYNMAI

TLSYGAMGLSVISIIKVYAEMNKHLREALQGANSELEFLSNHDDLTGLYNRRFYEQRMAH

LLERAEITGKTIGLIMFDLDDFKQVNDTHGHGMGDALLKMLGERLRHQVRDIDLIARLGG

DEFVVLMENMRSSDDLPEIAAKLVAAVEQPVKARNEVMALKVSCGFALYPGDGLSRAELE

EKADKAMYRAKKRGSSPDPSLILH

>WP_007227932.1

MTARVLPFVLLLLAGCTTSPPSNVNNICEIFEEKSGWYGDAHDAKKEWGSPIPVMMAIMH

QESRFVAKAKPPRKKIFGFIPGPRPSDAYGYSQAKKSTWKDYKRGGGNYGADRDDFGDAI

DFIGWYNEQSKKRSGISKRDTYGLYLAYHEGHGGYNRRTYKSKKWLTDVARKVERRAGSY

QQQLSTCEKDLEKGGWFFGW

>WP_007226969.1

MILECDIGNTRCKWRVVGEGAEENRGAFDCADGFGELPSLDGIRRVKVSSVAGSTVNEEL

TRTLASAKLEIEFARTSPLKAGVENAYADASKLGVDRWVAMIAGYNRCRGPVLILDAGSA

LTVDLVAANGKHLGGYITPGIQLMKSSLLAETDGVRFDRDNHSSGTAFGTDTASAVHAGV

VAAQVGAAIVAIEEAGRKVSAGFAILLTGGDANVICTNLPATISAEVTMVPELVLDGLQW

VLP

>WP_007225387.1

MTTNLSASDPVGMSFWLISMAMVAATVFFLIERDRVSGKWKTSLTVAGLVTLIAAVHYFY

MRDVWVATGETPTVYRYIDWLLTVPLLIIEFYLILSAITKVPVGVFWRLLAGSLIMLGAG

FVGEVNPDYVVSGFVVGMLGWVWIMYEIFLGEASKINAASGNAIAQKAYGAMRLLVTVGW

AIYPIGYVLGYFTGSTDSATLNLWYNVADLWNKVAFGLVIWAAAVADSE

>WP_007225373.1

MNEIKCPHCKKAFTIDEAGYAEIVKQVHNSEFDQQLHERLELAERETRNAVKLAEEQARS

KLLEAESTKNDEIRKLQSELEAGDYARKLAVSKALKAVEKERDELANKLIQAKNDSTNAS

KLADANHSKKLEKTSAEKDAAIRLLEEKLSASEDTKNHAVTKAVNVAERERDELKSKIYR

TKLENEIAETSLKDNYEAQLKDRDHEIERLKDMKARLSTKMVGETLELHCETEFNLIRAT

AFPRAYFEKDNDSRTGSKGDYIFRDSDEHNTESVSIMFEMKNEIDETATKKKNEDFFKEL

DKDRNEKQCEYAILVSLLEPENSLYNSGIVDVSHRYQKMYVIRPQFFIPIITLLRNAAEK

SLKYKKELAVVKEQNVDITNFESELDEFRSGFARNYELASKKFKTAISEIDKTIDHLQKT

KDALLGSENNLRLANNKADDLTVKKLTKSNPTMEAKFKELNNDEDI

>WP_007225063.1

MNIKQWLNPFDLVVSVRTGFALIIIRRALLLFAVLSAPLAFSADISGVWKHSKNTAWIEI

SLADSSATVLRNDKFPERVGRTILKDLQVDTSTQGLWHGLIYVEKLGDYKDVEVSLPEAG

RMLLKGKVGFMTRTVEWLRVDNIR

>WP_007224727.1

MKLLLVEDDEALAKALLIALRNEGFSVDHVATGNEAIAHGKNNIADIIILDLGLPDIDGL

TVLKELRANKIVTPMLILTARDDLSDKITALDGGADDYLSKPFEIKELMARIRALGRRMN

SSISSVITAGRVSLDSANHHVEVDAIETPLSRREYTVLKALMENVGRIQTKAALENKLYS

WGEEISSNAIEVHISNLRKKMPEGFIKTVRGVGYTIDRSSS

>WP_157361222.1

MAALGVVAALALASCSAGDGGSGESATGDLRVWLVGTDTPQEARDYLIDTFESENPGSTL

TIEEQAWGGLVDLLTTNLSGSDSPDLVEVGNTQAAAFTSAGAFLDLTADYDALGGDDLLP

GFVEAGSYDGKFYAAPLYSGSRLVFYKKDALAAAGLSVPTTLDEYVSNGEALAEANPGAS

GIWWPGQDWYNALPFIWENGGEVATFESSGWKSQFSSPGSIAGLKQVQDVMTNASRAPKD

ANETNPQVGYCEGTTLQLSAPSWVKWSILAPLDAETPGCPDEEANLGVYAMPGKDGGAAQ

VFAGGSNIAVSAKSAHPELAKKALAIILSDGFQEIYGANGLVPAKLSLADTLGTDEVAAA

ISEAAGAARLTPPSPKWADVEASGALTDFFVQIAQGGDVASLAKDLDAKIDSILNG

>WP_157361184.1

MVQNRIGRAVLAAAAAVAVSIAMVGCAPASESEQGEVELRFSWWGTDSRHELTNEALDLF

EEKNPGITVVRDFGGFDGYIDKLLTQAAGNNSPDVFQLYEEVLREFASRGQLYDLNEATS

QGLSLDGWDQGLLDTSTIDGSLSALQFGLTTQAFIFNTELFDQAGVSIPTEGWSWDDLAT

AAKKVSDGTDSGTFGVTDLSTGYQVFEVWANQNGESYLTDDGLGFSAGTLEDFWNYWADL

RASGAATPGSLTSEYPTPFDAIIASKAASGFIFANQMAAVQSSIEAEIAVDRMPGESPEA

GSYLRTAMNIAIGSKTEHPKEAAMLVDFLLNDPEAYAILGIDRGVPANPAVGDAATANVD

DITAKGLTVIDGVREDGAAPPVPPKPGAGNVNALFAELAQEVQFDRMSIKDAVASFIERA

EQELS

>WP_156788299.1

MSRGILGPALVFMVMAIICLSLILPLRQLLIAGTLIVCTDLAVILLYFSGAFVINIDPVQ

AFAAPASWFSYSIVPIYASILIMLVLSRFNKGIEQVLTELEEEKSAAFYLSEHDHLTGLP

NMRVMEIRAHQAILMADRGESKPALLFIDLDHFKVINDKRGHDVGDLVLQEVAIRIQSVI

REGDIVARAGGDEFLVLLPQANDTVDAETVSQKICVTLANPFRLDDDEIYISASFGIAMW

PEHGRDLKSLTRSADQAMYAVKTSGKNGFKVSTSSIENG

>WP_156788178.1

MGDDAGNKTFGSLLKSELRDTDVIARTGGEEFALLIRTDSYSVTIIKAKMIC

>WP_157361215.1

MPVWFEVTSLSILLLILAADLIMAYKRPHIPSTRESALWVGFYVSLALIFALMMFLLGDV

EQAGQFIAGWVTEYSLSIDNLFVFVIIMARFSVPRKYQQEVLMVGILIALVLRGIFILLG

AQLIENFSFIFYIFGAFLLYTAIRQVFENHDDMEETESGIIRFLRKHINISPVFDGGKMR

TVIDGKKVFTPILVVFVALGITDLVFAIDSIPAIFGITTDPFIVFTANIFALMGLRQLYF

LLGDLIDKLEYLHYGIAFVLGFIGVKLFFHALHINELPFINGGEHVEWAPEIGTWTSLIV

ILVSMAVSVIASLVKMNVDKKRELVSIDSE

>WP_157361178.1

MLSSPTFCWVLSRPQDPCQGLPAVITAHYCCGVSDTNNTPDIVPFVARRLLTADQQRTAV

AVAAPVGAPSSASPTSLGVDPGSQQLTAQILHEFGPLAPYVGRPTITDVFVNGAQQVWVD

RGGGLEPVNDLGLTEPELRALAVRLISLGGRHIDEATPCVDVRLAGGVRVHAVLPPISAT

GTLLSIRIPSREPFGLAELDLAGFFTEVPMQRVKGLVDARENLLISGASGAGKTTFLGAL

LGAASETERIVAIEDVAELRVEHAHFVSLEARQANLEGTGSYGLPALVREALRMRPDRLV

LGECRGAEIRELLSALNTGHDGGAGTLHANSLRDVPSRLEALGALAGLDASAIARQAVSA

IGAVLHLDRVGGRRRLTQVGRLILDENERLAIADDE

>WP_157044971.1

MVTAAIGAGDPAQGRALLWLNVAALLLVLLSCFAVIHSTHATRELYTQLQVLESRQWHLQ

EDYGRLLLEESTWASHYRVEKVARTELGMAEPDLAHYKVVRR

>WP_156788350.1

MVLSSLNVARAEVAEIVAAPIEFPSTMAWSAYNLGTTGYNQAVAIGKVLKDHYDTNLRVL

PGKNDVSRLLPLQRGRVQFSANGAATYFAQEGVFQFAEKQWGPMPLRIVMASNGETNQAL

GVAADMGIATYSDLRGKRVPFVRGAPALNVTTEAYLACGGLTWDDVERVDFPGYSAMWTG

IVNDQVDAAYGTTVSGPTRKLEASPRGIFWPPAPHDDAECWARMAKIVPFFQPHMATRGA

AISIANPHEGATYPYPILITLAKTDPDLVYDLAKVIDIHYDEYKSADPGSIGWAMDRQVF

RWVVPFHGGAVRYFKSIGVWDDATQAHNDRLIGRQDVLAKAWRVHKASYPGKEGFADAWG

KARVQALDANGFDPVWR

>WP_156788318.1

MKVPRLLTKLILCALFTSLVAAGFATKNLRDFLNTPMDIQGEGLAYLLEKGGSLSQVGVD

LSLLGVLENRRWLSIYSRISGRGTAIEAGEYWLEPGLTPLELIAKFEQGDVRFFQLTLVE

GWDMSQVLSRLRSADALINTFGADTRVLTADMLGLETSFPSLEGLLFPDTYRYHSGTTDR

ELLLQAYQRMQKVLNDEWSDRSKNLPYDNMYQALIMASLVERETGVAWERAQISGVFVRR

LKLGMRLQTDPAVIYGLGASYTGNLRSRHLKDGSNKFNTYRHHGLTPTPIALAGREAIHA

ALHPADGKTLYFVAKGDGTHYFSETLKEHQKAVRKYQIEQRRKDYSSTPVIKPAG

>WP_007223442.1

MDLATVIGVVGALAIIITSMVLSGGIGMFTNMSAVLIVFVGSMFVVLSKFGMSQFLGAGK

VAAKAFFFKSTDPSAMIDEIVVLADAARKGGLLSLEGKEVGNDFLQGGIQLLVDGHDPDV

VKALLSKDKDKTVERHEQGASIFAALGDVAPAMGMIGTLVGLVAMLSNMDDPKSIGPAMA

VALLTTLYGAMLANMVAIPISDKLILRRGEEEMNKSLVIDALLAIQSGQNPRVIDSMLRN

YLPASQRPQADE

>WP_009774002.1

MVNAGIRIGVVGATGQVGAVVRRLLEERDFPVAEIRYFASSRSAGTTLPFKGEQITVEDA

STADPTGLDVAIFSAGATTSKAQAPRFAAAGVTVIDNSSGWRMDPDVPLVVSEVNPHAID

QAVKGIIANPNCTTMAAMPVLKVLDAEAGLERLIVSTYQAVSGAGLAGGEELLEQAAAAV

AQNTMGLVEDGAAVTMPAPNKFPKNIAFNVVPLAGSIVDDGLNETDEEKKLRNESRKILE

LPGLLVSGTCVRVPVFTGHSLSINVEFARPLSPARATELLATAPGVSLSDIPTPLDAAGA

DPSFVGRIRADEGVPDGRGLALFISNDNLRKGAALNAVQIAELVAAKITAKVSA

>WP_007235113.1

MSEAIDIAVVGATGAVGEAMMEILEQREFPVGKLYALASERSAGKTVRFRGKSITVSDLA

EFDFSKTAIALFSAGGSVSEEHAPRAAASGCVVIDNTSHFRRQEDIPLVVPEVNPGALAA

YRSTRIIANPNCSTIQMLVALKPIYDAVGISRINVATYQAVSGTGKAAIEELAGQTARLL

NGQPTEAKVYSKQIAFNALPHIDTFEENGYTREEMKMVWETQKILEDPDITVNATCVRVP

VFYGHSEAVHIETNTKITADAARKLLQDAPGVTLTDGREDGAYPTAVTDGAGSNPVYVGR

IREDISHPTGLNLWVVADNLRKGAALNSIQIAELLVKEHF

>WP_007227922.1

MSELYDIAVVGATGAVGETMISILEERDFPVGNLYPLASSRSAGKTIMFNGNTVKVTDLA

EFDFSQAQIGLFSAGGSISEKYAPIAAEAGCVVVDNTSHFRRDEDIPLIVPEVNIEALAG

YMTRGIIANPNCSTIQMLVALKPIYDAVGIERINVCTYQAVSGTGKEAIEELAGQTARLL

NGQEAQCEVYPKQIAFNVLPHIDSFQENGYTREEMKMVWETQKIFGDHSIQVNPTCVRVP

VFFGHSEALHIETVDKISAEQARELLQNAPGVQVMDEQADGGYPTAVGDSAGSDPVFVGR

IREDISHPRGLDMWVVSDNVRKGAALNSVQIAESLIATYLD

>WP_009773280.1

MPDTHSERVIEDEIVTDFSERMSYGSYLELDTLLSAQTPQSTPEHHDEMLFIIQHQTTEL

WLKLVIHELTSARDLIANDNLSIALKRIARVKHIQRTLTEQWSVLATLTPSEYSQFRDYL

GSSSGFQSYQYRAVEFLLGNKNAGMLKVFESHPEAHALLSKLLAEPSVYDEFLRYLSRHG

YDIPEAVLNRDVTRGYEQNDDLIETFRHIYDNESEHWLAYEACEEFVDLEDNFQLWRFRH

MKTVMRIIGMKRGTGGSSGVGFLQKALDLTFFPELFAVRTEIGRS

>WP_007236374.1

MDMTIVYGVGMFTAIVLALVMVILAARSRLVSSGNVSIEINGGKTIEVPAGGKLLQTLAD

ANLFLASACGGGGTCAQCKCQVSDGGGSMLPTEESHFTRRQANDGWRLSCQTPVKQDMRI

QIPEEVFGVKQWECTVESNDNVATFIKELVLRLPEGESVDFRAGGYVQLECPPHNVNFDN

FEIGEEYKGDWERFGFFKYGSASEDTTIRAYSMANYPEEKGIVKFNIRIATPPPGSEGIP

PGIMSSWVFDLKPGDKITVYGPFGEFFAKETDAEMVFIGGGAGMAPMRSHLFDQLKRLNS

KRKITFWYGARSLKEMFYVEDYDGLQAENENFTWHTALSDPQPEDNWDGLTGFIHNVLFE

EYLKNHPAPEDCEYYMCGPPMMNAAVIKMLVDLGVERDNIFLDDFGG

>WP_007228144.1

MNAVQETLADWRDSLIDLFPDARPVRWVLLALVAYLLIAIIVGMIWSLPPDHFDPSEKAA

EYAAQDGGEVVTGSTTTAALMGVMETLLEKPGGYLHNDRFPPGIWLDNMPNWEYGALVQV

RDLSRAMREVFSRSQSQSTEDKDLAMAEPRYHFDSDSWILPSSESEYRQAQDYTRGYFRR

LSDSTQAEAQFFARADNLRYWLSTVNTRLGSLSQRLSASVGQRRINTDLAGDAGASQSTA

APREMEVKTPWLEIDDVFYEARGTTWALIHFLKALEVDFADVLAKKNARVSLQQIIRELE

ASQETLWSPLILNGTGFGLVANHSLVMASYISRANAAIIDLRDLLLQG

>WP_007223884.1

MDWQAIKERLVVWREDRFDTATESNSTKVVLIVAAVYLLLAISVGMYWSMMPAQFPVQEN

AIVMAERSQQSVAVGSTTTAALIQVISTLLDKPGGFMSNDVMPPGLWLDNIKNWEYGALI

QSRDLTRALRESFSRSQSQSKEDLDLGKAEPSLNFSSDSWTLPASESEYKSAVKHLNRYL

VRLEDTGNSGSQFYARSDNLRYWLAIVESRLGSLSQRLSASVGKRRLNTDLAGDSAAQQS

TSAPSELEIKTPWTEIDNVYYESRGTSWALLHFLRAIEVDFHEVLKKKNALVSLQQIIRE

LEATQQTIFSPMILNGSGFGVLANHSLVMASYISRANTAIIELRDLLSQG

>WP_009772683.1

MSMFGLKESEIATLVRAVSVDGGGNLDRASIENRFARATWSGLAEPGDRLAGRAIQQLGS

ARSLTAVVEHWDAEQFATELSADGDPVSGDDMRQAIDRWMPRLKSDTALIALRQAARFGS

RLLIPDDSLWPERLHDLDWHAPSALWVRGTDAALAGIVDGIALVGARAATGYGEHITMEA

SAGLVDRGYTIVSGAAYGIDGMAHRAALASHGLTVAFLAGGVDRFYPSGHDSLLSRIVEN

GAVISELPCGSPPTKWRFLQRNRLIAAASIATIVLEAGWRSGSLNTAGHAAALGRPLGAV

PGPVTSAASAGCHRLIRDYDAVCVTNPDQMAELAPLDRAPETDATMTTPESQPPTNLPPE

SSAPKVDADSPPSTETTRLLDALSVRSARTADDIASRSGLALATVRAHLGLLELDGRVVE

SERGWKQASQSRTA

>WP_007234196.1

MSYVLARNAAADQLWFEFLATKMLDRASKRKLLAQGFTPPQLLEPHGTWPFDGAELRASL

RVSKSRQVARLKPLLACHLLHWGRQSSGCDVYPPLLSGLSDAPLALFVSGDINCLSRPAI

AIVGTRRPSRDGLKLADQMGYQLAAAGFLVVSGLARGIDAAAHRGALRSGGQTLAVMATG

MDRIYPSEHYRLAEEVAAQGALLSEFCPGVVPHRGHFPRRNRTLSGLCLATVVIEAGHPS

GSLITANAAVEQGREVFAAPWSLFHRGGAGCLRLLSQGAQLIDTPAAVIEHLAVHLSGWA

ELTADALDYSSNAGETVSLAPLEPAKRQLLTLLGDGEHDLASLASALQCSSRQLLAMVTQ

LELQGYVEQTSAGLRAVRHP

>WP_007224080.1

MKDDETRAWLALSRIPHLPRRVLHRLILATSSAEEIFQLSAFELTAAKVGAEAQKMLREG

VDLRQVEQDFKTVQLQHIKLLPVSSTLYPALLKEINDPPPLLYIRGDLSVLDLPSLAMVG

SRRSSQAGGANAFRFARELAGAGFSIVSGMALGVDTQCHRGALAAGGSTVAVLGTGIDIV

YPRRNKELFESIVCQGAVISEFPMGTDPHPARFPRRNRIISGMSLGVLVVEAALQSGSLI

TARCAMEQGREVFAIPGSIHNSGSKGCHQLIKQGAKLVESVSDVMEELKGWCADAPPVLE

EKGAQNKVSKDLHERERLLLDIIGYDPVSIDSLQQRTDWPMHDLVALVTALELRGLLDCV

AGSYQRTV

>WP_007236375.1

FLAVSKKIQAAFGLGVAVVVVLTITVPVNNLIYQYLLADGALAWAGLPDVDLSFLGLLSY

IGVIAALVQILEMFLDRYVPALYAALGVFLPLITVNCAILGASLLMVERDYTFGESAVFG

AGAGVGWALAIVALAGIREKLKYSDVPDGLQGLGITFITVGLMSLGFMSFGGIDI

>WP_009772673.1

MADFSLEDLKTLRERLGTGMVETKNALVEAGGDLEKATELLRLRGAKSNAKRSDRSTSEG

LIAAQSSGTSTTIIELACETDFVAKSDKFVALGEAVAAAVAAAGASTVEEGLAAPAGSST

VAQLIDDEAAILGEKFELRRLTKLEGDSFEVYMHRTNKDLPPQVGVVVAYSGDDAETARG

IAQHISFAAPTYLSREEVPADDVENERRIVEEISRGEGKPDAALPKIVEGRLGAFFKQVA

LLDQDYARDNKVSIAKVSADAGITVTGFARFKVGA

>WP_148224520.1

MMSVRWTMPLLIAAVLMSSFAIIHSTHASRAYYANLQRLEGTHWYLQEDYSRLMLERSTL

ASPHRIAKMAQDELIMRAPDLATYRTIVEGAY

>WP_148224355.1

MIVILSGIVICFTLGKQALHAPMNLPQPDATVIVEQGDSLKQILTKLKSRGFIESSRLLE

LWARWQGVDRQIHTGEYLLVPGLSGIGFLERLGRGDVLSYKITLPEGITLQQALQRLHDD

RRLVRELRDAHDPLLLELVSPMTSPEGWFLPETYRFVAGDSDYDILRRAHHLMQRELIRV

WEARSSDTPLMTPYEALTLASIVERETSVAKERATIAGVFSRRLQAGMRLQTDPTVIYGL

GSDFDGNLKRRHLKDAANPWNTYRIKGLPPTPIALPGVAALEAAVRPASGAALYFVARGD

GYHVFSETIEEHNAHVQRYQLSRKVDYRSTPKGGD

>WP_007229556.1

MSNQQLGDFLAHSVELESEARERYLELAQAMIAHHNTDVAGFFNRMAEESRLHLEEVAEI

AQDIELPGLKAWEFGWPEEESPEAVSYEAVHYRMSLRQAMLLALENERAAEKFYRSFADA

SSDGETRHLAAQFSAEEASHAAQLEKMLGKLPPDREHHLEEDDQPHMPE

>WP_007234052.1

MSKTNIFVGVFLSLLLGAASQAQSSDLTNPTALVDNGPYTGDLQGLVERRIVRVLTVYGP

GRYYLDNGAKGVTAEYANRLEKVINESFDTGHLKVAVFVLPVARDELFLALEQGRGDIII

AGTTITPAREQRAAFTIPSSKPLKEILVMGPSAPPINSIDDLSGKSVYLRASSSYSDSIA

TLNERFTREGKALVTVEPMSELLEDEDLIEMVDAGLLPWTIVDDYKPTQWSGVFTNLTVR

NDIIFRKGSRHAWAVRQDNPELKKFLNNFLKDNKEGTLFGNILKNRYVRDFNWAANTNAE

SELQRYRDLEALFRRHGVSYGIDPALLAAQGFQESRLDQNVRSGAGAVGVMQLLPSTAAD

KNVGIPNIHEVDPNIEAGAKYLAFLKRRYFSTPGMDPLNGALLALAAYNAGPAKVRRLQE

TARTRGYDPYRWFDNVEVIAAEKIGRETVQYVANIFKYYLSYQMINRESARREAARQAAG

APTHEQRQN

>WP_007235497.1

MIKPRGRRIGLIRLSAIGDVCHAVATVQALQRHAPEDDITWIIGRTEAALVSDLPGITFI

VFDKKQGLTAFRNVLNEIAEPFDVLLHMQVSLRANILAAVVPAKAKLGFPKHLSKELHGM

VVNRRVPMPETPHVLEGFQHFAYALDVPTFAPTWSIPISEADQAWVRERLTAQKPYVVIA

PSASNAERNWLVDRYAALANHLQYRGYNVVLTASPAPSEVALAQQITALAGSNIINLAGQ

TTLKQLLAVVADATLVVAPDSGTAHMAVTQNTPVIGLYAHSNPNRTGPYRFQFLTIDAYQ

KNLQHLFSNSAKSNKWGVRLKGAHLMEDIALSEVIAKADEVLSEAPNPSDHNS

>WP_007235565.1

MFKIRTFNAISVKGLERFPRQSYEVGGEIGSADAMLLRSHKLQADEISASVTAIARAGAG

VNNIPLSHCTELGIPVFNTPGANANAVKELVAAGLLLASRDILGGIDFVNSLSEDLDEQA

MGPLLEAEKKRFAGAELKGKTLGVLGLGAIGSLVAQLGLELGMDVVGFDPAISIEAAWQL

PSSVKRMENMQALFSRADYISIHVPAIESTHHLINQETLKYFRSDACLLNFAREQIVDTE

AVAAALDKQGLGRYITDFPHPLLRGRKDCILMPHIGASTAEAEENCAIMGADQLRAFLEH

GNIRNSVNFPRLELERTTGSRIAITNTNLPGTLSHILTAIGDSQINVVDLLNKSRDEIAY

NLIDLNTTPPADLLEQLRGIEGVINVRCIPDQAAD

>WP_007227712.1

MAKQVLTLNQISLKGLERLPRDSYEIASEFSHPDAILLRSHKLQAQDIADSVLAIGRAGA

GVNNIPVAECSQRGIPVFNSPGANANAVKELVAAGLLLGSRGIVEGIQYVDTLSAMADKT

EMNKTLEAQKKQFKGSELEGKTLGVVGLGAIGSMVAEMALTMGMDVVGYDPALSVEAAWR

LSSQVRKADTLSALFGRCDFITLHLPVLDSTRGLINAELLSSTREGTCLLNFARQEIVDE

EALVQALDGDKLRKYIADFPSPALIGRDNVILMPHIGASTDEAEDNCAIMAANQLKDFLE

NGNIRNSVNFPNLSLERVSGCRLSVTNENVPKILGSVLSILADENINVIDMLNKSRNDIA

YNLIDVVGHTSDEVLDKMRALEGVVNVRMIGDCA

>WP_007226314.1

MSSISKTVAKTFGLILFFISIANAEITGIVVSVTDGDTIKVLDENSNQHKVRLTGIDAPE

RGQPFGQASKKYLASMVSGKEVFVESNKKDRYGRVLGKVWVQPADCPSCGKTLDINHAQL

LAGMAWWYRYYAKQQSPEDRGRYESAEDEAKARGWGLWSAASPINPYNWRKGRR

>WP_007229969.1

MSATKPDLVWDRIRTETQKHAQEEPVLASFLHSTILNHDSLECALSFHLASQLDSPTVTS

LLLREVMLQAMRADDAIGEAIRADLLAVVERDSASHELYIPFLYFKGFHALQSHRIAHWL

WHNNRKSMGLFFQNRISVEFGVDIHPAAKMGQGIMLDHATGLVIGETAVVGNNVSILQSV

TLGGTGKQDGDRHPKIGDGVLISAGAKILGNICVGDGAKVGAGSVVLEDVPPHTTVAGVP

AKVVGRPATNAPALDMNHDFFCDSGDVEG

>WP_007228043.1

MEELLSLFIRSIFIDNMALAFFLGMCTFLAISKKIDAALGLGIAVIVVLTITVPVNYLIY

NYLLADGALAWAGQPDLDLSFLGLLSYIGVIAAIVQIMEMFLDKFVPALYNALGVFLPLI

TVNCAILGATLFMVERDLDFAESVVFGAGSGVGWALAIVALAGIREKLKYSDVPDGLKGL

GITFIIVGLMSLGFMSFGGIDL

>WP_007227945.1

MDMTIVFGVAMFTAIVLALVAIILFARSALVSSGNVSIEINGEKTITVPAGGKLLQTLSE

SGLFLPSACGGGGTCAQCKCIINDGGGSMLPTEEGHFTKRDAAEGWRLSCQTAVKQDMKI

EVPEEVFGVKQWECTVESNPNVATFIKELTLKLPEGEHVDFRAGGYVQLECPAHHVKYSD

FDIEEEYRGDWEHFNFFKHESVVKEDVIRAYSMANYPEEKGVVKFNIRIATPPPGSEGIP

AGQMSSWVFNLKPGDKVKVYGPFGEFFAKDTDAEMVFIGGGAGMAPMRSHLFDQLKRVHS

DRKISFWYGARSLREMFYVEDYDMLARDNENFDWHVALSDPQPEDHWDGLTGFIHNVLFE

EYLKNHPAPEDCEYYMCGPPMMNAAVIQMLIDLGVEPENIMLDDFGG

>WP_050774034.1

MEVIVGEPRKPLPRGSDFYTPEQRLRRDSSVWTTVQGVLAPLQFLAFALSLVFVINFLAN

GTGYSAAVISVLIKTLFLFTIMVTGAIWEKVVFGRYLFAPAFFWEDVVSMLVIFLHVAYV

VSWLFDLQAPREQMWLAIAAYTAYVINAAQFLLKLRAARVGSSQNNTDSVNEYAVEVSR

>WP_040823654.1

MLLAGVDEVGRGPLAGDVVAAAVILDPANPIRGLDDSKKLTEKKREALFPEIQEKALSWF

VARASVREIDELNILQASLLAMKRAVEGLVLQPEHVLVDGNKLPRWAYSAEAVVRGDSRV

QVIGAASILAKVVRDREMVAFDDEYPGYGFAGHKGYPTRVHMTALDVLGVTPIHRSSFGP

VKRKIAQMNRP

>WP_009772453.1

MATPNPLDAVINLAKRRGFVFQSGEIYGGSRSAWDYGPLGMALKENIKKQWWQTIVQGRD

DVVGIDSAVILPRKVWEASGHVEVFSDPLVESLHTHKRYRADHLLEAYEEKHGHPPVNGL

ADIRDPDTGQPGSWTEPQNFSGLLKTFLGPVDNEEGMHYLRPETAQGIFTNFANVMGAAR

MKPPFGIGQVGKSFRNEITPGNFIFRTREFEQMEMEFFVEPGTDEEWHQYWIDESMKWYT

DLGIKPENLRFYEHAQEKLSHYSKRTVDVEYRFRFAGSEWGELMGIANRTDFDLRTHSEA

SGADLSYFDQAKDERWTPYVIEPAFGLTRALMAFLIDAYAEDEAPNAKGGVDKRTVLRLD

RRLSPVKVAVLPLSRNERLSPLARSVAADLRKFWNVDFDDAGAIGRRYRRQDEIGTPFCV

TIDFDSLDDNAVTVRERDTMEQKRMPLEELRGYLAQELIGC

>WP_007236092.1

MSEDRTVWVVDDDRSIRWVMEKALTQAGLLCQSFETAEALLEAITSGAPDVVISDIRMPG

IDGLALLGQLRAAYPELPVIITTAHSDLDSAVASYEEGAFEYLPKPFDVDEIVATVLRTP

TMRKERKAPVTELPDKPTEIIGNAPAMQEVFRAIGRLAHSQITVLINGESGTGKELVARA

LHRHSPRKDGPFIALNMAAIPRELMESELFGHEKGSFTGATARRAGRFEQADSGTLFLDE

IGDMPAETQTRLLRVLADGEFFRVGGAAPVKADVRIIAATHQNLETLVANGQFREDLFHR

LNVIRIHLPRLADRQEDIPKLMQFFLGKAAQELGVEGKVLSTSASRYLCQLPWPGNVRQL

ENTCRWLTVMAAGREIHPSDLPPELLEPAQSQRVDNATTWQDTLATWAQQRLAAGESNVL

RKALPEFERIMIAAALTHTGGKRAEAAETLGWGRNTLTRKIKELEEDGTPAKGA

>WP_007226588.1

MATYRNFALVAAIVIVAFYQTLVDLMGNWLKFDESQSHGLIIIALFIHLFTGQLKQLPSP

PATPNWLGLMGLSASSLVWCLAAMLNIEAIEQLILLPILFFLCWSSLGLRSTVTLTPSIA

LLIFAIPIWDYLTPTLIDASSYVVMTLIQLSSITAFIDGNSIYLPHGRIDIADGCSGLRY

FIIAIALAYYLILTSKTTHLTKVKVLGIAIALGLFTNWLRIYIIIMVAHFTEMESSLVKD

HELFGWFLFFIVCLPLVYFARSLPHYEPTTPSATSAGVTKLTLVVSVIALTSGPLLYQLM

NTKVTAPNLGNWQQLGYQQLSSPTNGPFQLPPSNLNLRKQSGATLRDVAIHWQNSQDSDL

VPYIANSLNRDYWTQLQTSTLQTPKQQSLQLNLYNRKATNQYRCTVSWYRVGGMETTHYN

IAKLLQIPALLSQHNQFSAAVISINSETANCDPHQQQLIDAAIETHNDIVQLTGLTEQ

#### 1.2 Testing dataset

**Bacteriocin**

**----------------------------------------**

>BAC005

TPVVNPPFLQQT

>BAC008

DIDITGCSACKYAAG

>BAC013

CVQSCSFGPLTWSCDGNTK

>BAC052

MNFLKNGIAKWMTGAELQAYKKKYGCLPWEKISC

>BAC076

ATYYGNGLYCNKQKCWVDWNKASREIGKIIVNGNVQHGPWAPR

>BAC110

GGAPATSANAAGAAAIVGALAGIPGGPLGVVVGAVSAGLTTAIGSTVGSGSASSSAGGGS

>BAC112

NKWGNAVIGAATGATRGVSWCRGFGPWGMTACALGGAAIGGYLGYKSN

>BAC116

KSYGNGVQCNKKKCWVDWGSAISTIGNNSAANWATGGAAGWKS

>BAC127

AVNDYEPGSMVITHVQGGGRDIIQYIPARSSYGTPPFVPPGPSPYVGTGMQEYRKLRSTL

DKSHSELKKNLKNETLKEVDELKSEAGLPGKAVSANDIRDEKSIVDALMDAKAKSLKAIE

DRPANLYTASDFPQKSESMYQSQLLASRKFYGEFLDRHMSELAKAYSADIYKAQIAILKQ

TSQELENKARSLEAEAQRAAAEVEADYKARKANVEKKVQSELDQAGNALPQLTNPTPEQW

LERATQLVTQAIANK

>BAC149

MSKRDCNLMKACCAGQAVTYAIHSLLNRLGGDSSDPAGCNDIVRKYCK

>BAC177

DWTCWSCLVCAACSVELLNLVTAATGASTAS

>BAC213

KCPWWNLSCHLGNDGKICTYSHECTAGCNA

>BAC215

VLSIVACSSGCGSGKTAASCVETCGNRCFTNVGSLC

>BAC217

GWVACVGACGTVCLASGGVGTEFAAASYFL

>BAC220

MGAIAKLVAKFGWPFIKKFYKQIMQFIGQGWTIDQIEKWLKRH

>BAC221

KRKKHRCRVYNNGMPTGMYRWC

>BAC226

ITSFIGCTPGCGKTGSFNSFCC

>BAC227

ANLGNYTSQCYSSQCYSSKCYSDSCYSSNCYTGRHMCGYTHGYSC

>BAC228

GIGTAQCAYFKALCYSGGSEWLGGYGGCGSTQNNCELARKYC

>ACA04496.1

MNKKNILPQLGQPVIRLTAGQLSSQLAELSEEALGGVDASTSIAPFCSYDGVDASTSIAP

FCSYDGVDASTSIAPFCSYDD

>CAA74348.1

MENKKDLFDLEIKKDNMENNNELEAQSLGPAIKATRQVCPKATRFVTVSCKKSDCQ

>AAK32694.1

MNKELNALTNPIDEKELEQILGGGNGVIKTISHECHMNTWQFIFTCCS

>sp|Q38L35|Q38L35_STRSL

MKNSKDVLNNAIEEVSEKELMEVAGGKKGPGWIATITDDCPNSIFVCC

>AAZ76602.1

MKSNLLKINNVTEVEKDMVTLIKDEDMELAGGSTPACAIGVVGITVAVTGISTACTSRCI

NK

>sp|Q52052|Q52052_9ZZZZ

MENLSVVPSFEELSVEEMEAIQGSGDVQAETTPVCAVAATAAASSAACGWVGGGIFTGVT

VVVSLKHC

>BAD74571.1

MAKLDDFDLDIVVKKQDNIVQPNITSKSLCTPGCITGILMCLTQNSCVSCNSCIRC

>BAB04172.1

MVNSKDLRNPEFRKAQGLQFVDEVNEKELSSLAGSGDVHAQTTWPCATVGVSVALCPTTK

CTSQC

>NP_940772.1

MENSKVMKDIEVANLLEEVQEDELNEVLGAKKKSGVIPTVSHDCHMNSFQFVFTCCS

>ABI99444.1

MNKKNILPQQGQPVIRLTAGQLSSQLAELSEEALGDAGLEASVAACITFCAYDGVEPSCT

LCCTLCAYDGE

>WP_013079673.1

MTSRFQLLRLGKADRLTRGALVGLLLEDITVARYDPM

>YP_142020.1

MATQTIENFNTLDLETLASVEGGGCSWRGAGGATVQGAIGGAFGGNVVLPVVGSVPGYLA

GGVLGGAGGTVAYGATCWWS

>NP_297556.1

MRELTSIEMNNVSGGDLATRIEASIVFGVSAFFAGSIWGGTRGGDGGGILGVGSIAQGVG

MVYGGIVGGIGGLIAGFVLDKNVTYNYAVGFYNSLFNGTFTK

>CAA11804.1

MKNLKEGSYTAVNTDELKSINGGTKYYGNGVYCNSKKCWVDWGQASGCIGQTVVGGWLGG

AIPGKC

>AAL39164.1

MKQYKVLNEKEMKKTIGGESVFSKIGNAVGPAAYWILKGLGNMSDVNQADRINRKKH

>AAU29394.1

ATRSYGNGVYCNDDKCWVNWNEANQQIAGIVISGWASGLAGMGH

>NP_964622.1

MKQFNYLSHKDLAVVVGGRNNWQTNVGGAVGSAMIGATVGGTICGPACAVAGAHYLPILW

TAVTAATGGFGKIRK

>AAL77872.1

MKKFKELKENELTAITGGSFVGYYLGRFLASATHYYGKTVTKGHMHSSTINN

>NP_664144.1

MTTMKELTINDMASISGGNAPGDAVIGGLGGLASGLKFCKLPHPVLTGGCVVGFTVGGAY

LGYTAN

>AAL09346.1

MMKKIEKLTEKEMANIIGGKYYGNGVTCGKHSCSVNWGQAFSCSVSHLANFGHGKC

>AAZ76605.1

MWGRILAFVAKYGTKAVQWAWKNKWFLLSLGEAVFDYIRSIWGG

>CAA75396.1

MDKIIKFQGISDDQLNAVIGGKKKKQSWYAAAGDAIVSFGEGFLNAW

>CAA75397.1

MNNALSFEQQFTDFSTLSDSELESVEGGRNKLAYNMGHYAGKATIFGLAAWALLA

>AAG02567.1

MKISKIEAQARKDFFKKIDTNSNLLNVNGAKCKWWNISCDLGNNGHVCTLSHECQVSCN

>YP_395172.1

MMIFKKLSEKELQKISGGVGIQKCSLGFSSREYLNKITKWIKHH

>ZP_03845684.1

MLYKIIYRSMILMEKFIELSLKEVTAITGGKYYGNGVHCGKYSCTVDWGTAIGNIGNNAA

ANWATGGNAGWNK

>AAY44084.1

MQNTKELSVVELQQILGGKRASFGKCVVGAWGAGAAGLGAGVSGGLWGMAAGGIGRELAY

MGANGCL

>YP_025353.1

MSGGDGKGHNSGAHDSGGSINGTSGKGGPSSGGASDNSGWSSENNPWGGGNSGMIGGSQG

GNGANHGGENTSSNYGKDVSRQIGDAIARKEGINPKIFTGYFIRSDGYLIGITPLVSGDA

FGVNLGLFNNNQNSSSENKGWNGRNGDGIKNSSQGGWKIKTNELTSNQVAAAKSVPEPKN

SKYYKSMREASDEVINSNLNQGHGVGEAARAERDYREKVKNAINDNSPNVLQDAIKFTAD

FYKEVFNAYGEKAEKLAKLLADQAKGKKIRNVEDALKSYEKHKANINKKINAKDREAIAK

ALESMDVEKAAKNISKFSKGLGWVGPAIDITDWFTELYKAVKTDNWRSLYVKTETIAVGL

AATHVTALAFSAVLGGPIGILGYGLIMAGVGALVNETIVDEANKVIGI

>AAN76832.1

METAVAYYKDGVPYDDKGQVIITLLNGTPDGSGSGGGGGKGGSKSESSAAIHATAKWSTA

QLKKTQAEQAARAKAAAEAQAKAKANRDALTQRLKDIVNEALRHNASRTPSATELAHANN

AAMQAEAERLRLAKAEEKARKEAEAAEKAFQEAEQRRKEIEREKAETERQLKLAEAEEKR

LAALSEEAKAVEIAQKKLSAAQSEVVKMDGEIKTLNSRLSSSIHARDAEMKTLAGKRNEL

AQASAKYKELDELVKKLSPRANDPLQNRPFFEATRRRVGAGKIREEKQKQVTASETRINR

INADITQIQKAISQVSNNRNAGIARVHEAEENLKKAQNNLLNSQIKDAVDATVSFYQTLT

EKYGEKYSKMAQELADKSKGKKIGNVNEALAAFEKYKDVLNKKFSKADRDAIFNALASVK

YDDWAKHLDQFAKYLKITGHVSFGYDVVSDILKIKDTGDWKPLFLTLEKKAADAGVSYVV

ALLFSLLAGTTLGIWGIAIVTGILCSYIDKNKLNTINEVLGI

>CAA33859.1

RFAHDPMAGGHRMWQMAGLKAQRAQTDVNNKQAAFDAAAKEKADADAALSTAMESRKKKE

DNKRDAEGKLNDELAKNKGKIPGLKIDQKIRGQMPERGWTEDDIKNTVSNGATGTSFDKR

SPKKTPPDYLGRNDPATVYGSPGKYVVVNDRTGEVTQISDKTDPGWVDDSRIQWGNKNDQ

>prf||1615299A

RFAHDPMAGGHRMWQMAGLKAQRAQTDVNNKQAAFDAAAKEKSDADAALSSAMESRKKKE

DKKRSAENKLNEEKNKPRKGVKDYGHDYHPDPKTEDIKGLGELKEGKPKTPKQGGGGKRA

RWYGDKGRKIYEWDSQHGELEGYRASDGQHLGSFEPKTGNQLKGPDPKRNIKKYL

>prf||1814449A

MSGGDGRGHNSGAHNTGGNINGGPTGLGGNGGASDGSGWSSENNPWGGGSGSGVHWGGGS

GHGNGGGNSNSGGGSNSSVAAPMAFGFPALAAPGAGTLGISVSGEALSAAIADIFAALKG

PFKFSAWGIALYGILPSEIAKDDPNMMSKIVTSLPAETVTNVQVSTLPLDQATVSVTKRV

TDVVKDTRQHIAVVAGVPMSVPVVNAKPTRTPGVFHASFPGVPSLTVSTVKGLPVSTTLP

RGITEDKGRTAVPAGFTFGGGSHEAVIRFPKESGQKPVYVSVTDVLTPAQVKQRQDEEKR

LQQEWNDAHPVEVAERNYEQARAELNQANKDVARNQERQAKAVQVYNSRKSELDAANKTL

ADAKAFIKQFERFAREPMAAGHRMWQMAGLKAQRAQTDVNNKKAAFDAAAKEKSDADVAL

SSALERRKQKENKEKDAKAKLDKESKRNKPGKATGKGKPVNNKWLNNAGKDLGSPVPDRI

ANKLRDKEFKSFDDFRKKFWEEVSKDPELSKQFSRNNNDRMKVGKAPKTRTQDVSGKRTS

FELHHEKPISQNGGVYDMDNISVVTPKRHIDIHRGK

>YP_194414.1

MVGSITPKLVYRLNGMHHVVAQVGAVNGDHVFALQLLHSAHDVLVYRKHKGLTKDINYTN

PHLVMTGFGHTQTWVPANDNDEYFVGAKPNSGNWTTQIARVKYPRLLSENYTSNTQLPRL

SHLNRVTDVPYDGHNHLHRVEASVSPNGKYFMIASIWNNGSGHFGLFDLDEVNQKLDENG

TTNTPITDLHCLSAFHIDNFDNPSVAPDEEEPTMIDSVQGYAIDNDKNIYISNQLSPKIN

HETGEVTTWARKIVKFPWGETDSNNWQVAMIDGIDLPDRYSEVESIHVNAPDDIYLTVAY

HQKIVKGDEYALRTLENQIFHIDNL

>AAT85003.1

MEMVDQKINAQVLSGVNDDISEMKSLTTLRKRVVTDGEVVSKSQNAFRLAGGKTGVILRN

DGNDFYALVTPEDQAQDGQWNTLRPLSFNLKTGRVSLRNGVDISGGAVVSHDAGISARTT

GPSPIINGQTYSSPSIHTDFTSGNITTQMMMCARVEAGKQDYGLLSYRDWQGSWNELRVR

SNAELDAGQFTKRNSEGWIKAAGNRNVNNDKDRKTNALWIQGAGDLSADFYHYERIGQHH

FLGLHVANGGAQGWYEFRNDGHAYTNGAWNSSSDARMKTQVEKIDNALEKLDCISGYTYL

KQGVTEAGVIAQELEEVLPQAVSKTELTLNDGSVLKDARSININGVVALLIEALKEERQA

RLALEKRLADLEARSGQETE

>AAT90328.1

MAEQKKNALVLNGINDDITELKSLTTLRKRVVSDGEIVSKGVNGFRLAGASTGVILRNDG

KNFNFLTTADGQARDGAFNTLRPFAFSLTTGRVSLRNGVDVSGGAFISHNAGITAQTTGP

DPLINGQTYRAPDIHTDFTTGKKTTTMLMGSRIVTGQEDYGLISYRDMKGSWNELHLKPN

AELSVGQLTKRNTEGWYKAAGVRKVNNGKDNKTNALWIQGAGDLSADFYHYERIGQHHLL

GLHVANGGAQGWYEFRNDGHAYTNGAWNSSSDARMKTDIEKIDNALDRLDRIGGYTYLKQ

GKPEAGVIAQEVETVLPQAVTQTALTLNDGSVLEDARAVNINGVVALLVEALREEKQARL

ALEARLQVLEGTDAVHS

>ZP_00378412.1

MNNLYRDLAPVTDSAWAEIEEEARRTFKRNIAGRRIVDVEGPTGFETSSVGTGHIRTLGS

TGGDISIKQRISQEFIELRVPFTVTRQAIDDVERGSGDSDWQPVKDAATTIAMAEDSAIL

HGLDSAGIGGIVPGSSNTPVAIPDAVEDFADSVAQALSGLRKAGVDGPYSLLLSSEEYTK

VSESTDHGYPVRDHLSRLLGDGEIIWAPALEGALLVSVRGGDYELHLGQDLSIGYHSHNG

DSVELYLQETFGFLALTDESSVPLHR

>YP_121242.1

MNNLHRELAPITSEAWAAIEEEAGRTFKRHIAGRRVVDVAGPHGVDFSAVGLGRTTGIAA

PDEGVQARQRVVAPLVELRVPFTLSREELDNVERGAKDTDLDAVKEAARRIAFAEDRAIF

EGYPAAGITGIRAAGSNAPITVPDDARLVPEAITQALTALRLAGVDGPYSVLLSAELYTE

VSETSDHGYPIRTHIERLIPDGEIIWAPAIDGAFVLTTRGGDYELTLGQDVSIGYLSHDA

DTVRLYFQQTMQFLVHTAEAAVALRR

**Non-bacteriocin**

**-------------------------------------------------------------------**

>WP_177374305.1

MPLDPIHAFYCRKDYLSLAQSCKVKSGGICARCGGVFDLNELRPHHKIELTLDNIDDTNI

TLNPDNIEVLCHACHNAVHSRFGNAIGAKRVYLVYGSPYAGKTTYVALVATRNDIVVDLE

RIHAAICVCGQYDKPDATKRIAFNIRDYLLDEIRTATPRRKWQDAYIIGSYPDRIDRDNF

VREYNAELVHIDTPQDACVKRAYEDIKRVAARDAVVGWIADYWRRYNE

>WP_142482129.1

MIEVSTRQDRANFYGSNTWRKLRLKALERDHYECQWCKEQGKVTTINDAILEVDHIKELE

HYPELATDMDNLRTLCKDCHNKRHGRMNYRGEERKKKFDDEWW

>WP_149877315.1

MTELVVVLFYWGDSMLVACSRCGVIHERGDCKIQDGYSERRIKKRGEVERFRSSALWQRK

RKKILDRDKHLCRVCLDGKYVPKAITNQRLEVHHIVPIVENEKLKLADDNLISICAFCHV

LAEKGNVPRDYLFGLVKIPPRGHYVEN

>WP_121704945.1

MVYIRKRHWVTYNSEKCKMYLRNDFQFECAYCGMKERDNVIGEGLFEKDHFVSRQSDVAW

NLDSYGNMVYSCCKCNGTKSDQNIEIILDPCKDDIYGGQHPHIRRLGAENHYKLYGVTPQ

GQQFIDDLKLNSRFYRKMRQTQAQNEEIRREIYQLLDKSSDFQPSGIDRKIEAYLENGTL

IDERSDEFRCGTSKAGEDVYRVLEKLKERDIKYELLFADDDLDVRVEYCGNIYDCEIRVT

DYAGTEKRGPIVKREKKKTWLKTGNVCGVLYYYKEQDIMDLYIYPNEERTEIVKLG

>WP_086414226.1

MSKEKTASRWGGKGVRVGIIACVIVMVAGIVLWMVQLTGGMIQTGMRNLDAWGLYLTLFM

FFVGLSAGGLIISSIPNAFGMKGFGDISKVAIWSSVCCTCMAIGFVVVDLGGPLRLWELF

VYSNLSSPLMWDILVLSIYLVLSLVYLWAYVRYEQGRMKHTGIRFVSAVALIVAILVHSV

TAWIFSLSPAHEFWHTALMAPWFVASALDCGTALVLIVVIVLRKVGYLELDQHNIVNLAK

MLAVFVCVDLYFFACDLLTSGYFGGTDGAEVVATLTTGSIAPFFWIQMAFMALALVILFV

PKLRTNGGVVVASALVIAGVFCKRCQIMLGGFQIANIDFADTANAFTITNWTDGYSLAGY

SGLVYWPEPIEFGVSLGVIALGALFLLLGLRYLPLRQAKRVSE

>WP_160213184.1

MYGPLIIAYLFFGGTAAGAMLVMAWWSLRFYRKANRPTSRMARAFAAMQQRVYPIGFVLL

LVSMLCLLGDMNYLERAFLVFTRPHPTPITFGAYALAAEMVLAAALSVANILQPLFFTGK

VRRFLEILTVPCSVLLMVYTGVYLFSIMGVPLWNNPAIIPLFFCSSLSSGISAVLLVDYF

ADGSTLLLRAAKPLQKAHMTCIAAEAIVAIAYGASLALDPAAEASLSLLTSPGIAPVLLI

GFAGFGMAVPFCMEGYTLARKECRTIPVSDFVCLVGGFCLRWCVIMCATH

>WP_152931844.1

MLAYLIGLVVTSTLIFIFSEEKVTYRLFAAAITGLTWPLSLIPSIISLMIRKSD

>WP_099730807.1

MIFAPVIRRAAYAQAPRSADLALQRFLMGALAQPAAAPAAGCTVTQDEKATTLQLDVPGL

AREQLSISIEGQVVQVQSVEGAPRKVQRAWELPTEIDASASTAKLENGVLTLTLVRLEPV

SKATTLTIH

>WP_048781921.1

MKASTLRKERKAQSIKSEIIGEVLNAVTHGIGVALAITALVLLLMKAVAVNNTTQIIAFS

VYGASLILLFLASTLYHSFKFTKAAKVFQRIDHSSIYLLIAGTYTPFCLIGIGGQQGFIF

CIAIWVFAIGGVIIEAFFLEKFSKISVFLYLAMGWVSIFTLKPLYESMGWGGILYLFLGG

LSYSLGTIFYKRKYHNFYHVVWHLFVLAGAIFMFLAVFKYL

>WP_005865615.1

MLTAMVIVFLVGYLMIALEHPLKINKAGTALLIGTILWVMYTYAAPFFIPRASAEEFSLF

LESFPSLGSLTFKEQCTRFVVEHQVLDSIGEIAETLIFLIGAMITVELIDAHGGFMFITN

HITTKKKKKLLALIAVITFFMSAVLDNLTTSIVMIMLIRKLLGNYKERWVFGSIIIIAAN

SGGAWSPIGDVTTIMLWVRGNISTSSTIPHLILPSIVSALIPVLIAMRFLHGNVTPPNAF

SQMEADNELLKKLKDKEKLSILIIGVLCLLFVPVFKTVTHLPPFMGILMGVGILWFYTEM

LYARKPIDEDLKLRLSKVVHRIDGATLLFFLGILLAVDALRCSGVLSDFAFWLDDTVGNV

YAVNLIIGALSSIVDNVPLVAGAIGMYPVATDAMVAAATDPAYLANFMQDGVFWQFLAYC

AGVGGSMLIIGSAAGVVVMGLERINFIWYLKNISLLALAGYLSGAVVYILQNLIL

>WP_169170392.1

MIFAPVVRRAAYATRLPMSDLALQRFLRAALARPAAAPGCSAAQDEKAITLQLDVPGLAR

EQLDITIDGAVVRVRSVDGAPRQVQRAWELPEAIDAAASGAKLEHGVLTLTLAKLAPVNR

ATHLTIQ

>WP_110511594.1

MAHYISLFVRAVFVENMALAFFLGMCTFLAVSKKVSTAFGLGVAVTVVLGISVPVNNLIY

NLVLRDGALVEGVDLSFLNFITFIGVIAALVQILEMILDKYFPSLYNALGIFLPLIAVNC

AIFGGVSFMVQRDYNFPESIVYGFGSGIGWMLAIVAMAGIREKMKYANVPAGLRGLGITF

ITTGLMALGFMSFSGVQL

>WP_116624776.1

MFQTLFSSSDVTSTALSVATVATLATAVLTMLGLSWTSERWRVPVALSAVALLASGLVYQ

SALNLWLTGHQLTPATRYVAWFVVQPLQICSVFFFARISGAVPSGVFWRTGAAAILMVLS

RYLGDAQIFNPTLGVLLSIAFWLYILGEMYFGAMAEVVRKSSRPIRLGYFWVRLIMTIGW

AIYPILHFVDVVIGAGHVPSVIVLYTVADLVNLIAVSLIVLAVAGEERF

>WP_140455306.1

MRTSPTRLLGTGLLTALLASLCCIAPLLALVGGVTGAISAFGWVEPFRPYLAGVTVAVLA

LAWYQRLKAGKSAAACACEGEASPTFWKSNKFLLAVSCVALLLLAFPEYAGAFYRQQPVA

KAASVQTDFTQSVKLQVKGMTCTGCEAHVNQEIGKLAGVFSVSTSYEKGNAIIKYDSTKV

KPMQILQAAKKTGYTVAIEDKKP

>WP_168247034.1

MPITKGHGNPTWTREETILALDLLYLHGKPVDRKHQDVSQLSEFLRRVDIHPAQSRTEKF

RNPDGVALKLQNLFSAVEPGRGLTYSKTDLEIVTAFPNSRKSELAEIARLLRSSLLTHEL

VEEHVDEEEVFIEGRWLTSRHRYRDIRLRKRLLQSLPKLCCEICDFSPPSLSRSIQESFF

EAHHTIPISAAEGSVATKVLDMALLCASCHRFIHRLIAEEKRWVTPAEARDYLTGKRNDK

LEDRS

>WP_120447158.1

MGLLGGFFNDLQKVVNDSVGNDYRKIYFSAHPEEQQECACCGATLYRGDSDFTIDHIIPQ

KYNGTNFVTNLQPMCRSCNSRKKDKIDALTLKYSGTMLINEIKNLNRKKEW

>WP_120424551.1

MRKHSKEYSSYMKSDAWSAKREERLQLDGNRCVMCGRPNGLQKDSVTPVLQVHHICYSNL

GNEPMSDLVSICPGCHKKIHKYYRRLRSWEDKEVVARA

>WP_120423357.1

MIKVERKITEKSRRAMDSLERERLKNGSYNTPEVNAALKEMFHGKCYICENKQITSYHIE

HLNPHHGNIELKYSWDNLFLSCAHCNNIKSDKFDPIIDCTKENVEDMIAFRKEGYFGRDE

KLIFDMLDSRIETQNTIKLLQEVYYGSTPQKKMEATILRRTLRKELSDFKEYVREYQESE

DEEKEDLMYLLQMQLSSSSPFAAFKRWLIRDNKDVYPELLEYID

>WP_160581195.1

MKKHQKTLAVLLTAAMLVSLTACSSGGKDSTTAAEAAKTETSAAAGTQAKAETKAEASQE

PVDIAVIVPQKRGDLGFTDSIYKGVEQVMADYADRVNITFTECAGDSSKFESTIYDVCDQ

GPDLIITPSGSGFADLIATKAAHDYSDIKFVLVDNSAAYAGITTDNVAGMSYKQNEATFL

TGALAALLNETGMIGYVAGMSNAVINDFTVGYIQGAQYINPDIKIQISYIGDFADSAKAK

ELAATQIGLGADVVAQVAGTAGLGVLDAAKEAGVWGIGVDADQAAAYKESNPEMSAIIAS

SAMKNGASLLVSIVDRFIGENDLPWGGIESQGLVEGAVEIAPIADNVPDEVKKQISELQE

KVIKGEIEVKSAFTISEEEFNEYVNSCQ

>WP_135856548.1

MATSYDYAPLFRSTVGFDRIFNLLENAQRARSISDWPPYDIIKTGDDSYRISVAVAGFAE

DELDITFQSNLLTVTGKKQDASADEYLHRGIAGRPFEHRFELADHVRVNGADLRNGLLSI

DLVREIPEALKPRKIDIQTSPALQHKVAPAQIEAQKAA

>WP_135901797.1

MSESRIRFRHLQAFLEVARQRSVAKAADFLHVSPPAVTKTLRELEEALGVAVVERDGRGI

RVTRIGEIFLRHAGTAITALRQGVDSVRQDGAINRYPIRIGALPTVSAKVMPHAMSLFLK

ENTSAAIKIVTGENAVLLEQLRTGALDLVVGRLAAPENMTGFFFEHLYSEQVLFVVRSGH

PLLELGADIFARLDAFPVLMPTRESVIRPFVDRLFITNGMTAPATEIETVSDSFGRAFLR

QSDAVWIISAGVVANELGSGAFVALPVDTEETKGPVGLTMRTDTAPSPAFSILLQTIREA

ARPRA

>WP_120435514.1

MLFIWESWHFWLFFVLGACFYYRRNEAEYE

>WP_120446042.1

MKKLLEEIEKNQELKAKIEELDKNPKSTPKDYIQAAAEYGIEIKEEDFKTTRGELSDDEL

DAVAGGKVCSCFVGGGGEGGRRDKICACVAFGAGEDNYADTLRCFCPLAGSGDTHDH

>WP_007225590.1

MDKLIRDSKAAFIIAMALTIWVCAPGAALAGPAYDTYGTVTFDGIKTDVNEYTGGVSSGS

LDLQWFNDHESKNFRYADNVTNALLWEINESSDSPTVWSLNVFFEVPTDARRMIWEDGCT

WIKGGIEGTSCDGLKGLPNGEAILDAYADGSHHFSSKKESKKESKKESKKGSKKESKKES

KKESKKESKKESKKESKKESKKESKKESKKDHEKHSQEGKKEAKMSYSTQTGSEEFSIGE

GEAANNWFGLQKWQDEDENVKDDGSWLTSREYLIENELCDTTFCDAWDSSFSVELLFLFN

TQAGAQNKIKSLTDESVNYAMRLHLSDEANGIDSVTVPEPGPGILLILGLAGLGFARRKA

Q

>WP_140972653.1

MNAYIREPVNAFTHLGGAVLSFIALLAMIVKVSVKMPSFASITAVILFGIGMMVLYTASA

VYHSVVASERVIYFFRKLDHSMIFILIAGTYAPFCLITLHSASGLLLFCLVYATAICGIV

FKMFWFSCPRWLSTAIYITMGWLIVLFFAPLAANLSTGGMVLLVLGGILYTIGGFIYGTK

PKWLEFKYMGHHEIFHVFVLLGSLAHFLSVYCYVI

>WP_169252559.1

MPDSVPTGLIAILRGVRSDEVLDIAEGIVDAGFSAIEVPLNSPDPLASISALVEKFGDSV

EIGAGTVLTADQVRECRQAGARIIVAPDTDRDVITTALELGLTPYPGAATPTEAFAAVKA

GATNVKLFPSSAVGISGMKAWREVLPSGTELFPVGGVGADNAAEWRKAGAAGLGLGSSLY

RRGDRPDDVRTQAQAIASAWAQSI

>WP_169251584.1

MLAVLIGACVGLGVLLWSGSTRRRLQSLLGEGRTDPTGTEAATPEAGPAESGTEPAVADD

QLAFDLDLVAICLTSGLPIPVALTLTAEATDDRSDLQRIARAMTIGGRRLADDDRLLPVL

EVFEFSEHTGVGPAPLIESVAEELRASSRRRRQEAAASLGVQLVLPLGVCILPAFLLLSV

VPVVISLLTDLTTVFF

>WP_169253902.1

MNTDEDVRAAIASLLRIGEPGDGLLKRLVDGIGPVAAQGIIMAVGRGETTAHEAVHGLTV

TGTEAAEHGQMPEAIDRWAVRAGDVETRGDDLDKIARIGGRLVIPDDDEWPRMLDDLGPA

APLGLWVRGAASLSTVLARAVAVVGARAASSYGTKCASDLAWDLAARGITVVSGGAFGID

AAAHRAAIAREAPTVAFMAGGVDRFYPAANADLFEQILSTGAIVSETAPGMTPMRHRFLL

RNRLIAASAQVSVIVEAGWRSGALNTARHALELSRQVAAFPGSVYSASSTGAHKLVREHE

AELVTCSDDVIALMDDETPALFDATAAGAAAENGERPPPPDPREALDEREKICLNALTVS

KPLDVGTIASRAGLTIADALNSLTTLDLAGMAERRDTGWVKLRTSRG

>WP_169253261.1

MNIRRMTVAAVAAVALALTGCGSDGGSGGGGETTDDLTLGTGGTSGTYYPLGGELASIFE

DNVDGVTVNYVESGASAENLGKIYQGEWQLGFTQSDTANTAVNGELEDLDGTKIDNVGWL

ASLYPEAAHIIVREDSGIESVEDLKGKKIAVGDAGSGTRAISDAILDAAGIGESDYTPEI

TDFGASTDMLADKQIDATIFVVGTPVAGLTQLAATTDVKLLGLDDDTTKTIEEGSGAESY

DIPADAYDFLDEDVPTVSVFASLVASTDQVSEDTAYNLTKALFEHTDDITLDVGKLITKD

SAMLGVGDVPLHPGAQKYFEEEGIELP

>WP_040823762.1

MKIFQRVSLRLSLALMCTALFAGTVAAESTASMPLVSPGPHEVVEKTTQQVMEVITSAKG

YYATDPQRFYSEIESVLEDVIDFDGFSRGVMGQYASKKMYVSLETDEEKSAFKERMRRFS

ATFRNGLVQTYAKGLLAFNGNRIDVLPPIESKDLSGTDSVTVTQHIFGEAEKPFVIQYKL

RPNRAGEWKLRNVTIEAINLGIVYRGQFNSAVRLYDGDIDKVIDNWSVDPTGSAKSS

>WP_007235350.1

MQRIDVYWRDIPAQVLIKRGRDRGKHLLSHRFQAAIDRAAMKAGKGGSDAYLEEWRRVTT

SIEAEGSVKDIAQEFGEQIEAQYSDDDIARLVAQKGFDEALT

>WP_007229782.1

MTTISLFPLSGVLLPHGKVPLQIFEQRYIDLVRSSMKTGDPFGIVWIRRGSEVAGRGRAS

SELGDWGTLARIVDWDQLPNGLLGITIQGEGRFDLYETETQSNGLVLGEVVYRDNPASVS

MEAKWQPMLDVLQSLESHPHVQQMGLQLDYGDAWNVAWALIQLLPLEEYLKYELLGLDAI

DEVMSELDLILNQISGED

>WP_007227784.1

MEKVIVCWRDVPAQVIIKHRRKRATVELSERFQKAIDKAAMRAGKADSDAYMEDWCRRSS

PHAGVGSLEDIAKNVADSIESDYSDDVLSKMVADVGYRVR

>WP_007228934.1

MAMDRPQVLIVDDDPRVCRLIQNMACSEQFEYTDIVDPRRLKTVYEDLLPEKIFLDLSMP

GMDGIEALTFLKDSGSTSHICLISGWSEKVLRSSCALGKKMGLNMSPPIHKPFRATEIRA

FLKADQFTHTSVPVSPILPRSRKFKEELKHAIYNVGEIQPFYQPIVDLKTGEVDSLEALC

RWHHPTRGILCPGDFLPLVEKYGLMKDLTYSLLKIILEDMACWDSMASRPNVSINLFPEL

LEEQSLPDVFMAHMKSAGIDPSRITIEITEQSNYGDSVQMMNVISRLRIAGFGLSLDDFG

TGFASMEKVKEIPFTELKIDRSFVSDLLHDPDAKAIVKSSISLAQEIGIPTVAEGIENRE

TLSWLISNGCTRGQGYLLCPPGDFDKSILLAQSSTNHKYEIGEFDSGDVSSSTDTVSTKE

CQTTATLTVTPDIT

>WP_007227234.1

MTGTILLVEDNELNRDMLIRRLVRAGKEVVSAADGQQALDLMRSEKPAVVLMDMNLPILN

GWTACRQARADDTIKHIPIIALTAHASDADRLNALEAGCDDYATKPVDFPGLLIKIEKLT

GNC

>WP_007225255.1

MTLILLVEDNDMNRDMLSRRLQRRQYRVTTANNGAIAVEKATLEKPDLILMDMELPIKDG

WTASREIKATLDTPIIALTAHALSGDRDKALAAGCDDYTTKPINFDHLVAMIEQYLGKKN

N

>WP_007226686.1

MQPPSNHPPLILASSSAYRRQLLLKLNLSFDCVNPCIDETAGSNETADQLVARLAREKAL

AVIHSHPAHLIIASDQVAVLDGVIMTKPGDHGSAIAQLRQCSDKKVIFYTGLTLLNSSTG

RLQNAVEPFSVYFRKLDATTIERYLANEKPYDCAGSFKVEGLGITLFKKLEGDDPNSLIG

LPLIQLTSMLANEGILRP

>WP_007233776.1

MTTHKVLIVDDELPIRDMLRMALETAGYECLEAETIDAAYHQIVDDRPDIVLLDWMLPGG

SGIELLRRIKRAEMTQDLPVIMLTAKAAEHNVIQGLDVGADDYITKPFALRELLARIKAL

LRRAKSSDDRNLLVVRDLTIDIDSRRAFVGEEALQLGPTEFNLLLFFMSHPERAYTRSQL

LDRVWGANIYVEERTVDVHIRRLRAALDAAEGDYSQLIQTVRGTGYRFSEQGG

>WP_007225159.1

MEEKTILIVDDEAPIREMIRMSLDMAGFNCREAADTREAYRVIADSKPDLVLLDWMLPGG

SGIELLRRLRKEELTADLPVIMLTAKTDEDNKIQGLDVGADDYITKPFAPREMLSRIKAL

LRRTSTGIGGSVIEVQGLKLDISSHRVYIDTRPVDMGPTEFRLLSFFMTHQERAYSRGQL

LDHVWGGNVYVEERTVDVHIRRLRKALESEGGCYNECVQTVRGTGYRFSSKSIPPA

>WP_007226710.1

MTQDTNPPGVPEGFRTLRNSAHAETHVGPFYYKKDDDELTLGFLAGDQHSNAIGGVHGGV

LMFFADYAVVMSAMKGQKENCATISASCDFVSSAHTGEWVEAEATITRRTGSMVFVSGRI

YVGDKTVMTVQSVLKRIIPREK

>WP_040541238.1

MSQRIVIDPNDPVTCPDCSHEFPLVQGISHHLIERYEEEYDQKLAEEREALEARAVRTAE

RQLSGRFEEQLGDLTDKLEDAQAEREKAHKKLTKEKARAADQAREEAAEELSDLKQQLGE

KDEKLEDFRKEELALRKAKQVLDQEKRDLELTLQRQLEEQQSALRAELGNEFQLREAELR

KKIDDAHSANEDLKRKLEQGSQQLQGEVLELELEEILSQAFPIDQVDAVSKGVRGADVIQ

TVNLRSGASAGKIVWETKRAENWSNKWVSKLKDDQQSVGGEIGVLVSTAYPANVDEPFTQ

IDGIWLVRPEFAKPLADALRAILIEAFRQRTASSGKNEKMEALYDYVCSAQFAQKVRAVL

DAYAAMRDDLEREKAAMQRLWKKREGQLERITVNVVGICGELQGLSTASLPHLDEIAPIE

VA

>WP_009773675.1

MGKNITVVAAIAVATLLMGACSSGEVIDPAEGNPGADLRAGEAYDPRAFEGESINMLLIE

HPFVNSLRPLIPDFEAATGITVNLEVLNEQQGFDKLQADLSAGVGNYDLFMTDPLHNWQY

SAAGWIEPLDGYVENDAITMPDYNIDDFAPGVLDAGRWNRELLTGLGEGSLWALPVNFES

YNLTYRPSMFEDAGVEVPTTYEDVLDVTESLATSLSGNNYPIVTRFDKYWDLTYLTFGSM

AESYGVNLLNDDGEVDIASDASVEVTDLFIDIIKAGSPQDASAFTWYEVLQGMASGRFAL

ALNEADLFAATYENDAESEIADDVGYALIPEGPEKRAASAWIWQLSMAQASADKGAAWTF

LQWLTSADVLMQTHLAGNMNPVRLSAWEDPELAALVDTWGSEPGQYREVLEGTAEIAAIN

YPPHPELTRALDRWAEAVQQSFFDGNTKANLESAASDIERILLP

>WP_009773511.1

MSPTIVRGMTWEHERGYGSVVKAAEAYRSVAPDVEVQWEYRSLQAFADQDLESLVEQYDL

LVIDHPHIPIAAEEKLFTPLNGRGFDTELATLATQSVGRSHESYKHLGQQWGLALDAAAQ

VAAYRPDLLESPPRNWDEVMALAEEGRVLWPFKPVDAYSSLITIAAGLGEDPMATAGVFL

SEEMLTRAMELLVRLARLVPADNAGFNPIQVADVLAESDIFAYSPLLFGYTNYSRVGYRS

KRVQYTDIPSSTRGVAGSLLGGAGIAVSSRSRVMDAAIAHAFWLASGPVQEGSYYDGGGQ

PGNAVAWESARTNSDSLDFFTGTRATLEGAYMRPRFATYIELQNAVSPFVTSALLGEITI

TELRERLDAGVAEWLVR

>WP_007235958.1

MTIDIDHRRVLIVDDQSTRAHHLIDALGMDAFEFDVANEVPDLQGALGPDSPWDCVLCNA

GLINVSWASVRRAMRNFDVQVPVIVVADEQNVDSMKTALGLGATDFFVKPHARPGLLKRS

IERCVNHRYLQRELKASKEDVERSNTELRHSLRVLEQDQQAGRQVQRALLPSGALHQGDY

WFSHTIVPSLYLSGDFTDYFSVGEDQIAFFLADVSGHGSSSAFATVLLKNLFARKRSDFL

RRGDHSVVSPKDMLELANNELLELAINKYATMIVGVLNFKSHQLTYSIAGHLPHPVLLDE

NSVRYLEGEGPPVGLMRDARYTQHEVVLPEHFVLALLSDGILELLGNGNLIEKEASLLSL

LEGPLESPRSLATRLGLEAVDPNHLPDDVAALFITRGFS

>WP_007227112.1

MGNCETKVLILEDDPLAASELKDCLAREGLHPSIARSKEHFEKIVEQHEFQLLIVDIGLP

DGSGLDVIREVREQSSVGIIVVSGYTTESDVVAAIELGADDYIKKPISIKELRAKVRRMM

IRTSGNGYSRSIAQPNNNEQKFFGDWHLDLDSHRLFYKINHEVGLTSAEYKILLALLNNC

DQVLSRHSLLNHLQSISSPYDERTIDGLINRVRKKLAIPASYEPVQKVRNAGYMFCETVR

TEHQQTTESPGSGFSLTAEKMTIDKAPSPTLEPPVFSSAPESGSTKLTH

>WP_007224478.1

MSSVEQLIQDSRLWRGKHYRDDHSQQTGNSISSGITQLDQQLHWRGWPLHSSSELLCEHW

GIGELSLLMPLLKKVSHKGRIAWINPPFIPYSPALLSQGITPEKCLLLYPSESDQWWAAE

QVLASSAFAIVMTWFTRQASNATPYRRLQAAAEKGHCLHFHFRPLSSKQQSSPARLRIQL

SSSASQLAVEVLKQPGGWSGQQLVISRPESLLFKQQAVEKWPVYHSSRPSYQVVNGRTDI

PSIIDPQHSDRLNDDQSIIHQPSSSAPTQPH

>WP_007223999.1

MAILALIQNNIDRFSDVTGRILAWLCLLLMLLSCSVVFIRYGLGAGSIALQESVTYLHGT

IFMLGAAYTLRHDGHVRVDIFYRNMSARSKAWVNCGGGIIFLLPLCVYFFISSWGFVQQS

WEFREISSEPGGIPAVFLLKTLIPLMAVNLGLQAFAETLRNLLILIAREDSVQL

>WP_007223681.1

MSDLSPQEIEILHEALDDEYLAWSTYDQVIEDFGEISPFINIREAESRHIEALCTLFNRY

GVPVPPNPWLGRVERYKSIQEACEAGVKAEIANGEMYERLMVATQRRDFLEVLGNLQEAS

QKRHLRAFERCVSRRGSGCGAGRGRGRGNGGRC

>WP_007234293.1

MDLATLLGLLGGLAVVGTAIFYGGAGPTFYNVPSILIVIGGTFMTVMVKFSLKQFLGAFK

VAGRAFSNKSHDPESLIAEIVNLANIGRKEGLLALEKAAISESFLKDGIQMLVDGSNQEV

VKAVMAKDMQQTMDRHNWGERVWRAVGDVAPAMGMIGTLVGLVGMLVNMNDPKAIGPQMA

VALLTTLYGAVLANMVALPIADKLHLRKSNEKLIHQMCIDGVLAIQAGQNPRVIESMLKA

YLDPAHRDKNANSGK

>WP_007236262.1

MSAVLVKELRERTGLGLLECKRALKEADNDIDAAIEALRKSSGMKAAKKAGRIAADGVVT

TRTAEDGSYGVLVEVNSETDFVARDENFLGFVGSVADTLYESRSADIDALKSGSLEQARE

ALVQKIGENIGIRRASLVTAENGVVGSYVHGNNRIAVLVELRGGDQDLARDVAMHVAAVN

PQVVSPADMPEALLEKERDIFTAQAQESGKPAEIIEKMIGGRIKKYLAENSLSEQAFVKD

PDVTVGQLVKAADAEVISFSRFEVGEGIEVDKVDFADEVAAQLKG

>WP_007225796.1

MAAVSASMVKELRDRTGLGMMECKKALVEAGGDIDAAIEEMRKNSGMKAAKKAGRTAAEG

VVTAKVAEDGSYGIVVEVNSETDFAARDESLLAFVATVSEKVFTEKQTDVKALMEGDLNT

AREALVQKIGENISVRRSEVVDSDGVVGSYVHSNNRIAVLVSLTGGDAELARDIAMHVAA

VNPQVVRPEDMPEDVVTQEKNIIKAQPDMEGKPEAIVEKMMIGRINKFLKENSLLEQAFV

KDPEITIGKLAKNAGAEVVSFVRYEVGEGIEKEEIDFAAEVAAQLNG

>WP_007233347.1

MSVATRVRTFFQQARDSLVLSGFAKTDGPMGKIVAGVGALYLIVMIVLAIWWSAAPPAFD

MTTKTQDYSTASGQPLVPGSATTLALIEVIDTLLEKDGGYTHNDLLPPGLFIDNMPNWEY

GVLVQSRDLARALREVLSRSQSQSREDVDLTLAEPRINFQSDSWILPASEREYRSANKYL

KAYLARLPEKGPEGARFYARADNLGFWLGMIEKRLGSLSQRLSASVGQRRLNTDLAGDPT

ASAATIDPEEQEIKTPWSEIDDVFYESRGAAWALIHLLKGAEIDFAGVLEKKNARVSLQQ

IIRELEATQGIVWSPIILNGSGFGLWANHSLVMANYISRANAALIDLRELLAQG

>WP_007230569.1

MSVVDRDLRAAGTEVLSLADPLYPPLLKTIPDPPPVLHVRGNPMLLARPQLAIVGARRAS

AAGLQAAHKLAVAAVRAGLGVTSGLALGVDGAAHRGALSAGGDTVAVMATGIETIYPHRH

EPLGQEIASSGCLVTEFPPGTKPLPYHFPKRNRIISGLSLGVLVVESALPSGSLITATSA

MEQGREVFALPWSISHKGGAGCLSLIRDGAKMVLGIEDILEELDSLFGLQQELSQVSTIP

SPESISEQDCLLLELLGFEVISLDQLVVASGLPVGQVMGELSSLELAGRVNRCPGGYIRS

R

>WP_007226893.1

MEHYLSLFVSAIFIKNMALSLFLGMCTLLALSKKMNAAIGLGIAVVVVLSITVPVNYLIY

TYLLREGALVWLSPEFASVDLSFLGLLSYIGVIAALVQILEMFLDKFVPALYNALGVFLP

LITVNCAILGASLLMVEREHDFGESVVFGVGAGVGWAIAIILLAGIREKMKYSDVPAGLQ

GLGITFITVGLMSLGFMSFGGIDI

>WP_007225006.1

MYKILTLNQISTKGLDKFPREDYEIASEFVTSDAVLVRSHKLQPADIQDSVLAIGRAGAG

VNNIPVDYCTEQGIPVFNTPGANANAVKELIVSALTLGSRGILEGIDYVNTLDDLTDGAA

MSKLLEKEKKRFKGNELSGKTLGVIGLGAIGSMVADTALALGMKVAGYDPALSVDAAWRL

SSEVEKVDNITSLVSRADFITLHLPVLDATRKMINRELLSHLKSGAVLLNFAREEIVDTT

AVVEVLDSGKLSKYIADFPTPELIGKRGAVLTPHIGASTDEAEENCAIMAAVQLKDFLEN

GNIKNSVNFPPLYLERTPQSGSVRLSISNRNVPKILGSILSILADENINVIDMLNKSRED

IAYNLIDLQSSPPEQVLEIMRKIDGVVNVRLIG

>WP_007235663.1

MTALSINLNKIALVRNSRVTTVPNIVSHAEMCISAGADGITVHPRPDQRHIRAQDCFDLQ

SALDVELNIEGNPFTEPRASDQPHVGDYPGFIALVQAISPAQVTLVPDSDQQLTSDHGFD

VARDGKRLEPLIKIFKDLGCRVSLFMDPDPSAMATVASLGADRIELYTESYARAHEVGDF

EVSLAAFQETAEAAFAARLGVNAGHDLNLSNLPDFKVPHLEEVSIGHSFTVDALRWGIAN

TIPRYQQALGKNC

### 2 Supplementary Tables

#### 2.1 Pearson correlation coefficient reduced features

**Table S1.** List of features obtained from correlation analysis

| aac_1 | dipep_67 | dipep_153 | dipep_239 | dipep_325 | pseudo_11 | dist_17 |
| --- | --- | --- | --- | --- | --- | --- |
| aac_2 | dipep_68 | dipep_154 | dipep_240 | dipep_326 | pseudo_12 | dist_18 |
| aac_3 | dipep_69 | dipep_155 | dipep_241 | dipep_327 | pseudo_13 | dist_19 |
| aac_4 | dipep_70 | dipep_156 | dipep_242 | dipep_328 | pseudo_14 | dist_20 |
| aac_5 | dipep_71 | dipep_157 | dipep_243 | dipep_329 | pseudo_15 | dist_21 |
| aac_6 | dipep_72 | dipep_158 | dipep_244 | dipep_330 | pseudo_16 | dist_22 |
| aac_7 | dipep_73 | dipep_159 | dipep_245 | dipep_331 | pseudo_17 | dist_23 |
| aac_8 | dipep_74 | dipep_160 | dipep_246 | dipep_332 | pseudo_18 | dist_24 |
| aac_9 | dipep_75 | dipep_161 | dipep_247 | dipep_333 | pseudo_19 | dist_25 |
| aac_10 | dipep_76 | dipep_162 | dipep_248 | dipep_334 | pseudo_20 | dist_26 |
| aac_11 | dipep_77 | dipep_163 | dipep_249 | dipep_335 | pseudo_21 | dist_27 |
| aac_12 | dipep_78 | dipep_164 | dipep_250 | dipep_336 | pseudo_22 | dist_28 |
| aac_13 | dipep_79 | dipep_165 | dipep_251 | dipep_337 | pseudo_23 | dist_29 |
| aac_14 | dipep_80 | dipep_166 | dipep_252 | dipep_338 | pseudo_24 | dist_30 |
| aac_15 | dipep_81 | dipep_167 | dipep_253 | dipep_339 | pseudo_25 | dist_34 |
| aac_16 | dipep_82 | dipep_168 | dipep_254 | dipep_340 | pseudo_26 | dist_35 |
| aac_17 | dipep_83 | dipep_169 | dipep_255 | dipep_341 | pseudo_27 | dist_37 |
| aac_18 | dipep_84 | dipep_170 | dipep_256 | dipep_342 | pseudo_28 | dist_38 |
| aac_19 | dipep_85 | dipep_171 | dipep_257 | dipep_343 | pseudo_29 | dist_41 |
| aac_20 | dipep_86 | dipep_172 | dipep_258 | dipep_344 | pseudo_30 | dist_44 |
| dipep_1 | dipep_87 | dipep_173 | dipep_259 | dipep_345 | amphipseudo_21 | dist_47 |
| dipep_2 | dipep_88 | dipep_174 | dipep_260 | dipep_346 | amphipseudo_22 | dist_49 |
| dipep_3 | dipep_89 | dipep_175 | dipep_261 | dipep_347 | amphipseudo_23 | dist_50 |
| dipep_4 | dipep_90 | dipep_176 | dipep_262 | dipep_348 | amphipseudo_24 | dist_52 |
| dipep_5 | dipep_91 | dipep_177 | dipep_263 | dipep_349 | amphipseudo_25 | dist_53 |
| dipep_6 | dipep_92 | dipep_178 | dipep_264 | dipep_350 | amphipseudo_26 | dist_55 |
| dipep_7 | dipep_93 | dipep_179 | dipep_265 | dipep_351 | amphipseudo_27 | dist_56 |
| dipep_8 | dipep_94 | dipep_180 | dipep_266 | dipep_352 | amphipseudo_28 | dist_58 |
| dipep_9 | dipep_95 | dipep_181 | dipep_267 | dipep_353 | amphipseudo_29 | dist_59 |
| dipep_10 | dipep_96 | dipep_182 | dipep_268 | dipep_354 | amphipseudo_30 | dist_61 |
| dipep_11 | dipep_97 | dipep_183 | dipep_269 | dipep_355 | amphipseudo_31 | dist_62 |
| dipep_12 | dipep_98 | dipep_184 | dipep_270 | dipep_356 | amphipseudo_32 | dist_63 |
| dipep_13 | dipep_99 | dipep_185 | dipep_271 | dipep_357 | amphipseudo_33 | dist_64 |
| dipep_14 | dipep_100 | dipep_186 | dipep_272 | dipep_358 | amphipseudo_34 | dist_65 |
| dipep_15 | dipep_101 | dipep_187 | dipep_273 | dipep_359 | amphipseudo_35 | dist_66 |
| dipep_16 | dipep_102 | dipep_188 | dipep_274 | dipep_360 | amphipseudo_36 | dist_67 |
| dipep_17 | dipep_103 | dipep_189 | dipep_275 | dipep_361 | amphipseudo_37 | dist_68 |
| dipep_18 | dipep_104 | dipep_190 | dipep_276 | dipep_362 | amphipseudo_38 | dist_69 |
| dipep_19 | dipep_105 | dipep_191 | dipep_277 | dipep_363 | amphipseudo_39 | dist_70 |
| dipep_20 | dipep_106 | dipep_192 | dipep_278 | dipep_364 | amphipseudo_40 | dist_71 |
| dipep_21 | dipep_107 | dipep_193 | dipep_279 | dipep_365 | comp_1 | dist_72 |
| dipep_22 | dipep_108 | dipep_194 | dipep_280 | dipep_366 | comp_2 | dist_73 |
| dipep_23 | dipep_109 | dipep_195 | dipep_281 | dipep_367 | comp_3 | dist_74 |
| dipep_24 | dipep_110 | dipep_196 | dipep_282 | dipep_368 | comp_4 | dist_75 |
| dipep_25 | dipep_111 | dipep_197 | dipep_283 | dipep_369 | comp_5 | dist_76 |
| dipep_26 | dipep_112 | dipep_198 | dipep_284 | dipep_370 | comp_6 | dist_77 |
| dipep_27 | dipep_113 | dipep_199 | dipep_285 | dipep_371 | comp_10 | dist_78 |
| dipep_28 | dipep_114 | dipep_200 | dipep_286 | dipep_372 | comp_11 | dist_79 |
| dipep_29 | dipep_115 | dipep_201 | dipep_287 | dipep_373 | comp_13 | dist_80 |
| dipep_30 | dipep_116 | dipep_202 | dipep_288 | dipep_374 | comp_15 | dist_81 |
| dipep_31 | dipep_117 | dipep_203 | dipep_289 | dipep_375 | comp_16 | dist_82 |
| dipep_32 | dipep_118 | dipep_204 | dipep_290 | dipep_376 | comp_17 | dist_83 |
| dipep_33 | dipep_119 | dipep_205 | dipep_291 | dipep_377 | comp_18 | dist_84 |
| dipep_34 | dipep_120 | dipep_206 | dipep_292 | dipep_378 | comp_19 | dist_85 |
| dipep_35 | dipep_121 | dipep_207 | dipep_293 | dipep_379 | comp_21 | dist_86 |
| dipep_36 | dipep_122 | dipep_208 | dipep_294 | dipep_380 | tran_1 | dist_87 |
| dipep_37 | dipep_123 | dipep_209 | dipep_295 | dipep_381 | tran_2 | dist_88 |
| dipep_38 | dipep_124 | dipep_210 | dipep_296 | dipep_382 | tran_3 | dist_89 |
| dipep_39 | dipep_125 | dipep_211 | dipep_297 | dipep_383 | tran_4 | dist_90 |
| dipep_40 | dipep_126 | dipep_212 | dipep_298 | dipep_384 | tran_5 | dist_91 |
| dipep_41 | dipep_127 | dipep_213 | dipep_299 | dipep_385 | tran_6 | dist_93 |
| dipep_42 | dipep_128 | dipep_214 | dipep_300 | dipep_386 | tran_10 | dist_94 |
| dipep_43 | dipep_129 | dipep_215 | dipep_301 | dipep_387 | tran_11 | dist_96 |
| dipep_44 | dipep_130 | dipep_216 | dipep_302 | dipep_388 | tran_14 | dist_97 |
| dipep_45 | dipep_131 | dipep_217 | dipep_303 | dipep_389 | tran_16 | dist_99 |
| dipep_46 | dipep_132 | dipep_218 | dipep_304 | dipep_390 | tran_17 | dist_100 |
| dipep_47 | dipep_133 | dipep_219 | dipep_305 | dipep_391 | tran_18 | dist_102 |
| dipep_48 | dipep_134 | dipep_220 | dipep_306 | dipep_392 | tran_19 | dist_103 |
| dipep_49 | dipep_135 | dipep_221 | dipep_307 | dipep_393 | tran_20 | dist_105 |
| dipep_50 | dipep_136 | dipep_222 | dipep_308 | dipep_394 | tran_21 | ss_1 |
| dipep_51 | dipep_137 | dipep_223 | dipep_309 | dipep_395 | dist_1 | ss_2 |
| dipep_52 | dipep_138 | dipep_224 | dipep_310 | dipep_396 | dist_2 | ss_3 |
| dipep_53 | dipep_139 | dipep_225 | dipep_311 | dipep_397 | dist_3 | ss_4 |
| dipep_54 | dipep_140 | dipep_226 | dipep_312 | dipep_398 | dist_4 | ss_5 |
| dipep_55 | dipep_141 | dipep_227 | dipep_313 | dipep_399 | dist_5 | ss_6 |
| dipep_56 | dipep_142 | dipep_228 | dipep_314 | dipep_400 | dist_6 | qso_1 |
| dipep_57 | dipep_143 | dipep_229 | dipep_315 | pseudo_1 | dist_7 | qso_8 |
| dipep_58 | dipep_144 | dipep_230 | dipep_316 | pseudo_2 | dist_8 | qso_15 |
| dipep_59 | dipep_145 | dipep_231 | dipep_317 | pseudo_3 | dist_9 | qso_16 |
| dipep_60 | dipep_146 | dipep_232 | dipep_318 | pseudo_4 | dist_10 | qso_17 |
| dipep_61 | dipep_147 | dipep_233 | dipep_319 | pseudo_5 | dist_11 | qso_20 |
| dipep_62 | dipep_148 | dipep_234 | dipep_320 | pseudo_6 | dist_12 | pssm_2 |
| dipep_63 | dipep_149 | dipep_235 | dipep_321 | pseudo_7 | dist_13 | pssm_18 |
| dipep_64 | dipep_150 | dipep_236 | dipep_322 | pseudo_8 | dist_14 | pssm_85 |
| dipep_65 | dipep_151 | dipep_237 | dipep_323 | pseudo_9 | dist_15 | pssm_274 |
| dipep_66 | dipep_152 | dipep_238 | dipep_324 | pseudo_10 | dist_16 | pssm_295 |

#### 2.2 CVFE reduced features

In the following tables S2-S5, *c*, *e* and *p* indicate count of disjoint sub-parts, count of iterations and ratios of recurring iterations for the extraction of common features, respectively.

**2.2.1 Table S2.** List of features obtained from CVFE (*c* = 2, *e* = 5, *p* = 0.4)

| pseudo_3 | dist_93 | dipep_22 | aac_2 | aac_13 | aac_4 | qso_8 |
| --- | --- | --- | --- | --- | --- | --- |
| pseudo_5 | pseudo_2 | comp_16 | aac_3 | pseudo_26 | dist_75 | dipep_220 |
| pseudo_28 | dipep_43 | comp_3 | dist_82 | aac_12 | dipep_191 | aac_1 |
| aac_5 | dipep_211 | comp_10 | dipep_11 | dipep_280 | comp_2 |  |

**2.2.2 Table S3.** List of features obtained from CVFE (*c* = 2, *e* = 5, *p* = 0.6)

| pseudo_3 | pseudo_28 | dist_93 | dipep_43 | dipep_22 | comp_3 | aac_2 | dist_82 |
| --- | --- | --- | --- | --- | --- | --- | --- |
| pseudo_5 | aac_5 | pseudo_2 | dipep_211 | comp_16 | comp_10 | aac_3 | dipep_11 |

**2.2.3 Table S4.** List of features obtained from CVFE (*c* = 2, *e* = 5, *p* = 0.8)

| pseudo_3 | pseudo_28 | dist_93 | dipep_43 | dipep_22 |
| --- | --- | --- | --- | --- |
| pseudo_5 | aac_5 | pseudo_2 | dipep_211 | comp_16 |

**2.2.4 Table S5.** List of features obtained from CVFE (*c* = 2, *e* = 10, *p* = 0.4)

| pseudo_3 | aac_5 | comp_3 | dipep_43 | aac_13 | dipep_191 | dist_75 | aac_12 |
| --- | --- | --- | --- | --- | --- | --- | --- |
| pseudo_5 | pseudo_2 | dipep_211 | aac_11 | dipep_22 | comp_16 | dist_99 | comp_2 |
| dist_93 | pseudo_28 | aac_3 | comp_10 | dist_82 | dipep_11 | comp_18 | dist_3 |

#### 2.3 HFE reduced features

The lists of features obtained from the HFE method—using bin values of 5 and 10 to discretize each feature—are presented in Tables S6 and S7. In each tables, the first 91, 181, and 302 features correspond to β values of 15, 30, and 50, respectively.

**2.3.1 Table S6.** List of features obtained from HFE with bin = 5

| pseudo_11 | dist_22 | dist_28 | dipep_316 | dipep_376 | dipep_79 |
| --- | --- | --- | --- | --- | --- |
| ss_5 | dist_19 | amphipseudo_26 | dipep_246 | dipep_16 | dipep_133 |
| dist_72 | tran_20 | dipep_152 | dipep_399 | dipep_356 | dipep_202 |
| dist_75 | dist_27 | dipep_135 | dipep_191 | dipep_46 | dipep_170 |
| ss_6 | pseudo_29 | tran_19 | amphipseudo_25 | dipep_337 | dipep_119 |
| pseudo_1 | dist_79 | amphipseudo_38 | dipep_331 | dist_88 | dipep_4 |
| pseudo_2 | dipep_228 | dipep_97 | dipep_320 | dipep_226 | dipep_206 |
| pssm_85 | aac_16 | dipep_17 | dipep_312 | dipep_103 | dipep_144 |
| pseudo_20 | dist_105 | dist_61 | dipep_334 | dipep_361 | dipep_68 |
| ss_3 | aac_14 | amphipseudo_28 | dipep_193 | dipep_168 | dipep_165 |
| pseudo_4 | dipep_388 | dipep_328 | dipep_251 | dipep_85 | dipep_300 |
| pssm_18 | tran_17 | dipep_166 | dipep_330 | dipep_208 | dipep_307 |
| ss_1 | dist_58 | dipep_368 | amphipseudo_24 | dipep_322 | dipep_243 |
| pseudo_13 | pseudo_22 | dipep_35 | dipep_181 | dipep_238 | dipep_349 |
| pseudo_10 | dist_78 | dipep_241 | dipep_383 | dipep_139 | dipep_255 |
| aac_2 | dipep_286 | dipep_156 | dipep_267 | dipep_162 | dipep_49 |
| aac_11 | comp_11 | dist_29 | dipep_189 | dipep_34 | dipep_194 |
| pseudo_14 | dist_85 | dipep_310 | dipep_387 | dipep_105 | dipep_136 |
| comp_16 | dist_100 | dipep_390 | dipep_210 | dipep_200 | dipep_39 |
| pseudo_16 | pseudo_12 | aac_19 | amphipseudo_31 | dipep_394 | dipep_109 |
| dist_70 | pseudo_24 | dipep_52 | dist_11 | dipep_185 | dipep_265 |
| pseudo_6 | dipep_219 | dipep_33 | dipep_379 | dipep_91 | dipep_205 |
| pssm_274 | dipep_62 | amphipseudo_35 | dipep_7 | dipep_398 | dipep_65 |
| dist_69 | aac_1 | dipep_160 | dipep_81 | dipep_345 | dipep_355 |
| dist_4 | pseudo_3 | dipep_237 | dist_76 | dipep_149 | dipep_98 |
| pseudo_8 | dipep_26 | dipep_314 | dist_18 | dipep_327 | dipep_116 |
| dist_7 | dipep_108 | dipep_288 | dipep_317 | dipep_318 | dipep_326 |
| dist_67 | dipep_21 | dipep_213 | dipep_131 | dipep_89 | dipep_378 |
| pseudo_17 | comp_13 | dist_15 | amphipseudo_30 | dipep_13 | dipep_73 |
| pseudo_9 | dipep_8 | dist_103 | dipep_252 | dipep_382 | dipep_324 |
| dist_26 | dipep_182 | dipep_301 | dipep_375 | dipep_256 | dipep_249 |
| dist_10 | dipep_102 | dipep_295 | dipep_146 | dipep_296 | dipep_121 |
| aac_15 | dipep_143 | dipep_341 | dipep_5 | dipep_58 | dipep_245 |
| dist_23 | dist_55 | dipep_303 | dist_1 | dipep_63 | dipep_370 |
| dist_24 | pseudo_30 | dist_65 | dipep_100 | dipep_227 | dipep_115 |
| comp_15 | dist_90 | dipep_45 | dist_30 | dipep_386 | dipep_126 |
| dist_96 | dist_97 | dipep_145 | dipep_53 | dipep_289 | dipep_93 |
| dist_20 | amphipseudo_36 | dipep_348 | dipep_51 | amphipseudo_27 | amphipseudo_32 |
| dist_21 | amphipseudo_40 | dipep_12 | dipep_99 | dipep_106 | dipep_400 |
| aac_6 | dipep_24 | dipep_147 | dipep_371 | dipep_113 | dipep_374 |
| dist_102 | dipep_64 | dipep_362 | dipep_231 | dipep_90 | dipep_94 |
| pseudo_19 | tran_4 | dipep_11 | dipep_396 | dipep_333 | dipep_377 |
| pssm_2 | pseudo_25 | dipep_294 | dipep_50 | dipep_285 | dipep_59 |
| dist_53 | tran_10 | qso_1 | dipep_315 | dipep_140 | dipep_47 |
| comp_4 | aac_13 | dist_68 | dipep_302 | dipep_28 | qso_8 |
| dist_66 | tran_1 | dist_93 | dipep_72 | dipep_130 | dipep_293 |
| aac_5 | tran_21 | dipep_372 | dipep_257 | dipep_258 | dipep_235 |
| aac_8 | dipep_159 | dipep_3 | dipep_9 | dipep_360 | dipep_163 |
| dist_86 | dipep_282 | dipep_332 | dipep_27 | dipep_215 | dipep_127 |
| pseudo_28 | dist_52 | dipep_204 | amphipseudo_39 | dipep_209 | dipep_283 |
| aac_3 | dipep_311 | dipep_384 | amphipseudo_23 | dipep_203 | dipep_278 |
| tran_18 | dipep_6 | dipep_14 | dipep_69 | dipep_15 | dipep_244 |
| pseudo_7 | tran_5 | dist_74 | dipep_175 | dipep_290 | dipep_347 |
| pseudo_26 | dipep_158 | dipep_111 | dist_62 | dipep_187 | dipep_269 |
| comp_17 | dipep_321 | dipep_188 | dipep_325 | dipep_381 | dipep_346 |
| ss_2 | amphipseudo_34 | dipep_230 | qso_20 | dipep_55 | dipep_389 |
| dist_91 | dipep_268 | dipep_236 | dipep_272 | dipep_19 | dipep_260 |
| pssm_295 | aac_18 | dipep_75 | dipep_336 | dipep_216 | dipep_358 |
| tran_16 | dist_89 | dipep_132 | dipep_56 | dipep_229 | dipep_342 |
| dist_73 | dipep_36 | dipep_233 | dipep_261 | dipep_266 | dipep_367 |
| dist_99 | pseudo_21 | dipep_151 | dipep_280 | dipep_291 | dipep_87 |
| dist_82 | dipep_211 | dipep_150 | dipep_154 | dipep_118 | dipep_29 |
| dist_16 | dist_25 | dipep_120 | dipep_344 | dipep_254 | dipep_161 |
| pseudo_23 | tran_3 | dipep_60 | dipep_217 | dipep_253 | dipep_32 |
| dist_81 | dist_59 | dipep_110 | dipep_319 | dipep_357 | dipep_350 |
| comp_2 | dist_37 | dipep_207 | dipep_276 | dipep_277 | dipep_393 |
| dist_56 | comp_3 | dipep_364 | dipep_172 | dipep_222 | dipep_78 |
| dist_8 | tran_2 | dipep_313 | dipep_297 | dipep_71 | dipep_122 |
| dipep_31 | pseudo_27 | dipep_287 | dipep_183 | dipep_177 | dipep_271 |
| dist_12 | dipep_48 | dipep_197 | dipep_44 | dipep_247 | dipep_66 |
| ss_4 | dist_13 | comp_6 | dipep_1 | dipep_23 | dipep_299 |
| dist_49 | qso_17 | dipep_80 | dipep_74 | dipep_42 | dipep_82 |
| dist_77 | dipep_43 | dipep_76 | dipep_54 | dipep_84 | dipep_134 |
| dist_9 | tran_11 | dipep_273 | dipep_124 | dipep_117 | dipep_125 |
| aac_12 | dipep_61 | dipep_292 | dipep_262 | dipep_234 | dipep_174 |
| comp_5 | comp_1 | dipep_199 | dipep_164 | dipep_270 | dipep_167 |
| comp_10 | dipep_141 | amphipseudo_37 | dipep_38 | comp_19 | dipep_373 |
| dist_5 | qso_16 | amphipseudo_29 | dipep_214 | dipep_20 | dipep_275 |
| dist_2 | dist_3 | dipep_107 | dipep_391 | dipep_385 | dipep_129 |
| comp_21 | dipep_305 | dipep_196 | amphipseudo_21 | dipep_86 | dipep_178 |
| dist_63 | dipep_157 | dipep_397 | dipep_155 | dipep_248 | dipep_138 |
| dist_47 | dipep_88 | dipep_190 | dipep_225 | dipep_195 | dipep_179 |
| aac_17 | dipep_232 | dist_14 | dipep_153 | dipep_148 | dipep_259 |
| pseudo_15 | dipep_223 | dipep_308 | dipep_335 | dipep_218 | dipep_365 |
| aac_4 | dist_84 | dipep_41 | dipep_239 | dipep_123 | dipep_95 |
| dist_34 | dist_41 | dipep_284 | dipep_306 | dipep_352 | dipep_242 |
| dist_38 | dipep_395 | dipep_10 | dipep_176 | dipep_173 | dipep_70 |
| dist_35 | dist_44 | dipep_212 | dipep_343 | dipep_169 | dipep_279 |
| dist_71 | dipep_171 | dipep_112 | dipep_128 | dipep_392 | dipep_67 |
| dipep_104 | comp_18 | dipep_304 | dipep_380 | dipep_309 | dipep_329 |
| dist_83 | dist_80 | dipep_30 | dipep_57 | dipep_339 | dipep_298 |
| tran_6 | dipep_114 | dipep_250 | dipep_37 | dipep_186 | dipep_353 |
| dipep_2 | dipep_192 | aac_7 | dipep_363 | dipep_263 | dipep_354 |
| dist_6 | dipep_264 | dipep_22 | dipep_137 | dipep_224 | dipep_366 |
| dist_50 | dipep_220 | amphipseudo_33 | dipep_142 | dipep_338 | dipep_184 |
| dist_94 | aac_10 | dipep_281 | dipep_351 | dipep_83 | dipep_40 |
| dist_17 | pseudo_5 | dipep_96 | dipep_274 | dipep_180 | dipep_101 |
| dist_87 | tran_14 | dipep_323 | dipep_77 | dipep_25 |  |
| pseudo_18 | aac_20 | dipep_18 | dipep_221 | dipep_198 |  |
| dipep_201 | qso_15 | dipep_92 | dipep_340 | dipep_359 |  |
| dist_64 | aac_9 | amphipseudo_22 | dipep_240 | dipep_369 |  |

**2.3.2 Table S7.** List of features obtained from HFE with bin = 10.

| pseudo_2 | dipep_395 | aac_1 | dist_1 | dipep_288 | dipep_73 |
| --- | --- | --- | --- | --- | --- |
| pssm_2 | dipep_22 | dist_65 | dipep_271 | dipep_366 | dipep_179 |
| pseudo_4 | dipep_268 | tran_20 | qso_1 | dipep_309 | dipep_57 |
| pssm_18 | dist_81 | dist_22 | dist_89 | amphipseudo_33 | dipep_217 |
| pseudo_6 | dipep_182 | dipep_68 | dipep_159 | dipep_221 | dipep_344 |
| pssm_274 | dipep_108 | dist_3 | dipep_256 | dist_30 | dipep_276 |
| pseudo_11 | dist_97 | dist_59 | dipep_51 | dipep_372 | dipep_172 |
| pseudo_14 | dist_105 | dipep_306 | aac_9 | dipep_113 | dipep_343 |
| pseudo_20 | dist_44 | dipep_281 | dipep_9 | dipep_341 | dipep_297 |
| ss_5 | comp_17 | dipep_262 | dist_61 | dipep_303 | dipep_155 |
| pseudo_10 | pseudo_26 | dipep_75 | dipep_46 | amphipseudo_37 | dipep_338 |
| dist_75 | dipep_286 | dist_14 | dipep_400 | dipep_161 | dipep_183 |
| pssm_295 | dipep_390 | aac_16 | dipep_158 | dipep_39 | dipep_54 |
| pseudo_7 | aac_4 | dipep_142 | dipep_310 | dipep_42 | dipep_352 |
| pseudo_16 | dipep_114 | dipep_131 | dipep_162 | dipep_32 | dipep_167 |
| pseudo_1 | dipep_181 | aac_14 | dipep_321 | dipep_72 | dipep_170 |
| dist_47 | dipep_241 | dipep_35 | dipep_71 | dipep_45 | dipep_238 |
| dist_72 | dipep_384 | dipep_290 | dipep_351 | dipep_283 | dipep_346 |
| pseudo_13 | comp_5 | dipep_296 | dipep_340 | dipep_145 | dipep_225 |
| dist_7 | pseudo_27 | dipep_291 | pseudo_21 | dipep_377 | dipep_137 |
| ss_6 | dist_35 | dipep_246 | dipep_116 | dipep_348 | dipep_195 |
| pseudo_15 | dist_19 | dipep_228 | dipep_55 | dipep_335 | dipep_239 |
| pssm_85 | pseudo_24 | dipep_261 | tran_2 | dipep_147 | dipep_91 |
| dist_77 | qso_17 | comp_3 | dipep_361 | dipep_194 | dipep_149 |
| pseudo_17 | dipep_210 | tran_5 | dipep_254 | dipep_139 | dipep_354 |
| ss_3 | dist_29 | dipep_301 | dipep_16 | dipep_132 | dipep_285 |
| dipep_31 | dist_73 | dipep_364 | dist_84 | dipep_332 | dipep_275 |
| dist_16 | tran_16 | dipep_188 | dipep_186 | dipep_67 | dipep_380 |
| ss_1 | dist_68 | dist_25 | amphipseudo_24 | dipep_260 | dipep_381 |
| dist_91 | dist_58 | comp_6 | dipep_226 | dipep_176 | dipep_78 |
| dist_64 | dist_94 | dipep_212 | aac_10 | dist_74 | dipep_248 |
| dist_17 | dipep_213 | dipep_193 | dipep_223 | dipep_336 | dipep_140 |
| aac_2 | dipep_110 | dist_13 | dipep_50 | dipep_56 | dipep_363 |
| dist_10 | dist_87 | dipep_388 | dist_28 | dipep_126 | dipep_198 |
| pseudo_19 | dist_63 | pseudo_22 | dipep_43 | dipep_229 | dipep_84 |
| aac_11 | dist_56 | dipep_166 | dipep_273 | dipep_230 | dipep_165 |
| dist_70 | dipep_171 | dipep_107 | dist_41 | dipep_236 | dipep_200 |
| dist_2 | dipep_64 | dist_100 | dipep_360 | dipep_369 | dipep_235 |
| comp_16 | dipep_331 | dipep_28 | tran_19 | dipep_69 | dipep_376 |
| dist_82 | pseudo_30 | dipep_206 | dipep_121 | dipep_307 | dipep_356 |
| dipep_61 | pseudo_23 | dipep_106 | amphipseudo_31 | dipep_60 | dipep_85 |
| dist_69 | dipep_284 | dipep_37 | dipep_257 | dipep_20 | dipep_398 |
| pseudo_8 | comp_2 | dipep_274 | amphipseudo_22 | dipep_40 | dipep_129 |
| dipep_211 | dipep_282 | aac_20 | dipep_34 | dipep_317 | dipep_103 |
| pseudo_9 | dist_50 | aac_19 | comp_19 | dipep_18 | dipep_329 |
| dist_4 | dipep_146 | dipep_322 | dipep_232 | dipep_337 | dipep_83 |
| dipep_201 | aac_12 | dipep_124 | amphipseudo_23 | dipep_41 | dipep_227 |
| dist_67 | aac_13 | dipep_7 | dipep_23 | dipep_308 | dipep_90 |
| dipep_102 | dist_38 | aac_7 | dipep_70 | dipep_112 | dipep_300 |
| qso_15 | dipep_292 | dipep_115 | dipep_394 | amphipseudo_39 | dipep_185 |
| dipep_2 | dipep_4 | dist_103 | dipep_218 | dipep_327 | dipep_345 |
| dipep_220 | dipep_33 | dipep_30 | dipep_375 | dipep_316 | dipep_318 |
| dist_85 | tran_14 | dipep_74 | qso_16 | dipep_79 | dipep_89 |
| pseudo_3 | dipep_251 | dipep_392 | dipep_305 | dipep_163 | dipep_58 |
| dist_26 | dist_8 | dipep_383 | dipep_88 | dipep_323 | dipep_249 |
| dipep_111 | dist_12 | dist_62 | dipep_240 | dipep_92 | dipep_259 |
| dist_79 | dipep_10 | dipep_153 | dipep_157 | dipep_242 | dipep_279 |
| dipep_26 | dipep_80 | amphipseudo_34 | amphipseudo_29 | dipep_173 | dipep_15 |
| dist_23 | ss_4 | dipep_13 | dist_80 | dipep_247 | dipep_222 |
| dipep_311 | dipep_386 | aac_18 | dipep_175 | dipep_177 | dipep_93 |
| dist_5 | dist_49 | dipep_101 | pseudo_5 | dipep_133 | dipep_245 |
| dist_99 | pseudo_25 | dipep_396 | dipep_148 | dipep_334 | dipep_180 |
| dipep_6 | dipep_12 | dipep_144 | dipep_252 | dipep_169 | dipep_347 |
| aac_15 | pseudo_29 | dipep_266 | dipep_192 | dipep_374 | dipep_19 |
| dipep_104 | dipep_214 | dipep_17 | amphipseudo_25 | dipep_253 | dipep_298 |
| comp_15 | dist_9 | dipep_270 | dist_11 | dipep_29 | dipep_243 |
| dist_24 | dipep_76 | dipep_27 | dipep_154 | dipep_47 | dipep_82 |
| dipep_21 | dipep_313 | dipep_399 | dipep_325 | dipep_379 | dipep_123 |
| dipep_191 | dipep_382 | dipep_287 | amphipseudo_28 | dipep_330 | dipep_350 |
| pseudo_12 | comp_10 | qso_8 | dipep_119 | dipep_244 | dipep_367 |
| dist_96 | dipep_151 | amphipseudo_35 | amphipseudo_26 | dipep_267 | dipep_357 |
| dipep_204 | comp_21 | tran_3 | dipep_152 | dipep_134 | dipep_277 |
| dist_20 | dist_71 | dipep_197 | amphipseudo_38 | amphipseudo_32 | dipep_117 |
| dist_78 | dipep_48 | dipep_250 | dipep_333 | dipep_136 | dipep_355 |
| dipep_219 | dipep_3 | comp_13 | dipep_97 | dipep_299 | dipep_365 |
| dist_21 | dist_34 | dipep_199 | qso_20 | dipep_81 | dipep_178 |
| dipep_11 | dipep_362 | dipep_8 | dipep_328 | dipep_370 | dipep_95 |
| aac_6 | dist_93 | dipep_143 | dipep_263 | dipep_168 | dipep_125 |
| dipep_391 | dipep_295 | dist_55 | dipep_368 | dipep_234 | dipep_385 |
| dipep_208 | dist_37 | dist_15 | dipep_312 | dipep_339 | dipep_59 |
| dist_102 | dipep_302 | tran_1 | dipep_160 | dipep_122 | dipep_86 |
| dist_66 | dipep_264 | dipep_203 | dist_76 | dipep_63 | dipep_184 |
| dist_53 | dipep_207 | dipep_202 | amphipseudo_30 | dipep_359 | dipep_25 |
| dipep_36 | dipep_231 | tran_4 | dipep_156 | dipep_258 | dipep_342 |
| aac_17 | dipep_371 | dist_52 | dist_18 | dipep_389 | dipep_349 |
| tran_6 | dipep_135 | amphipseudo_36 | dist_88 | dipep_293 | dipep_378 |
| dipep_24 | dist_90 | amphipseudo_40 | amphipseudo_27 | dipep_109 | dipep_353 |
| tran_18 | dist_6 | dipep_164 | dipep_233 | dipep_49 | dipep_265 |
| comp_4 | dipep_304 | dipep_320 | dipep_66 | dipep_5 | dipep_94 |
| aac_3 | dipep_77 | dipep_294 | dipep_289 | dipep_100 | dipep_205 |
| aac_5 | dist_83 | dipep_44 | dipep_278 | dipep_255 | dipep_326 |
| pseudo_28 | dipep_387 | dipep_189 | dipep_38 | dipep_269 | dipep_65 |
| ss_2 | dipep_216 | dipep_150 | dipep_130 | dipep_99 | dipep_98 |
| comp_11 | dipep_141 | dipep_209 | dipep_128 | dipep_53 | dipep_138 |
| dipep_14 | pseudo_18 | dipep_324 | dipep_224 | dipep_319 | dipep_358 |
| dist_86 | tran_10 | comp_1 | dipep_190 | dipep_174 | dipep_373 |
| aac_8 | dipep_314 | dipep_393 | dipep_52 | amphipseudo_21 | dipep_87 |
| dipep_120 | tran_11 | dipep_397 | dipep_105 | dipep_118 |  |
| dipep_280 | tran_17 | dipep_315 | dipep_96 | dipep_187 |  |
| comp_18 | dipep_215 | dipep_1 | dipep_237 | dipep_272 |  |
| dipep_62 | dist_27 | tran_21 | dipep_196 | dipep_127 |  |

**3. Supplementary Figures**

**3.1 Hypergraph**

**Figure S1 –** Hypergraph representation of the training dataset

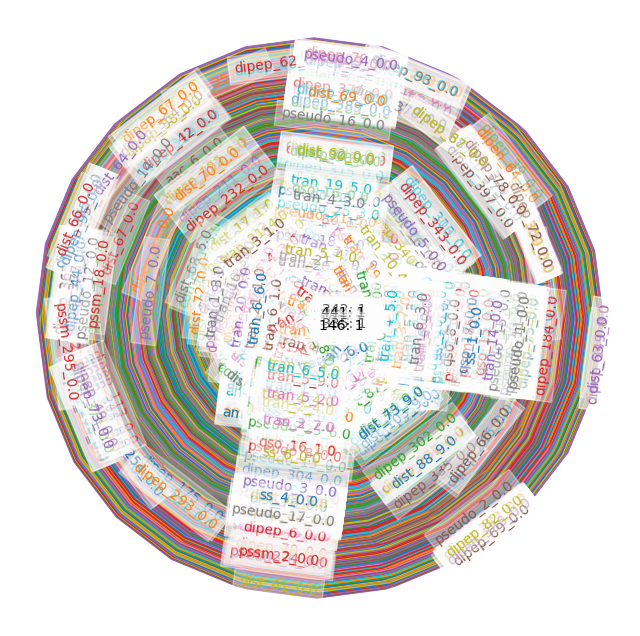

**3.2 Confusion matrices**

**Figure S2 –** Confusion matrices of the machine learning model built with the reduced feature sets

| 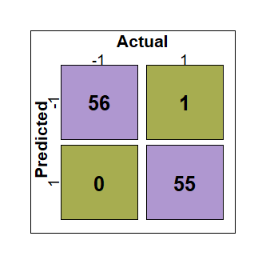   (a) **CVFE (*c* = 2, *e* = 5, *p* = 0.4)** | 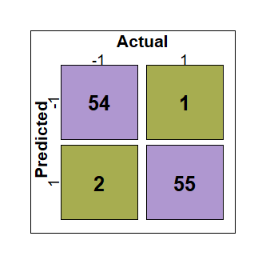   (b) **CVFE (*c* = 2, *e* = 5, *p* = 0.6)** | 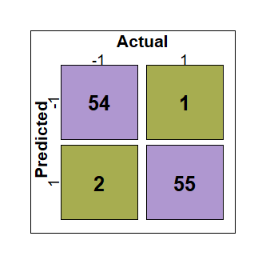   (c) **CVFE (*c* = 2, *e* = 5, *p* = 0.8)** | 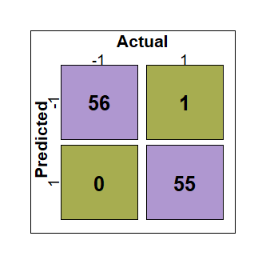   (d) **CVFE (*c* = 2, *e* = 10, *p* = 0.4)** |
| --- | --- | --- | --- |
| 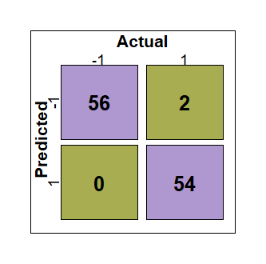   (e) **HFE (bin = 5, *β* = 15)** | 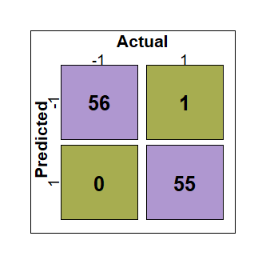   (f) **HFE (bin = 5, *β* = 30)** | 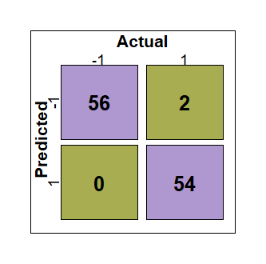   (g) **HFE (bin = 5, *β* = 50)** | 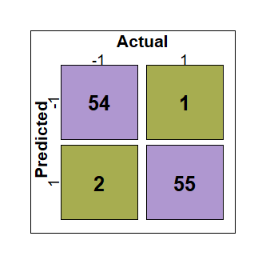   (h) **HFE (bin = 10, *β* = 15)** |
| 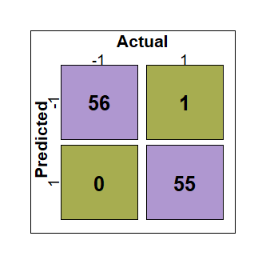   (i) **HFE (bin = 10, *β* = 30)** | 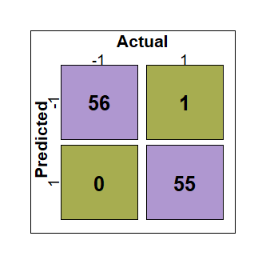   (j) **HFE (bin = 5, *β* = 50)** |  | |

**3.3 SHAP summary plot**

**Figure S3 –** SHAP summary plot of the best XGBoost model for all features

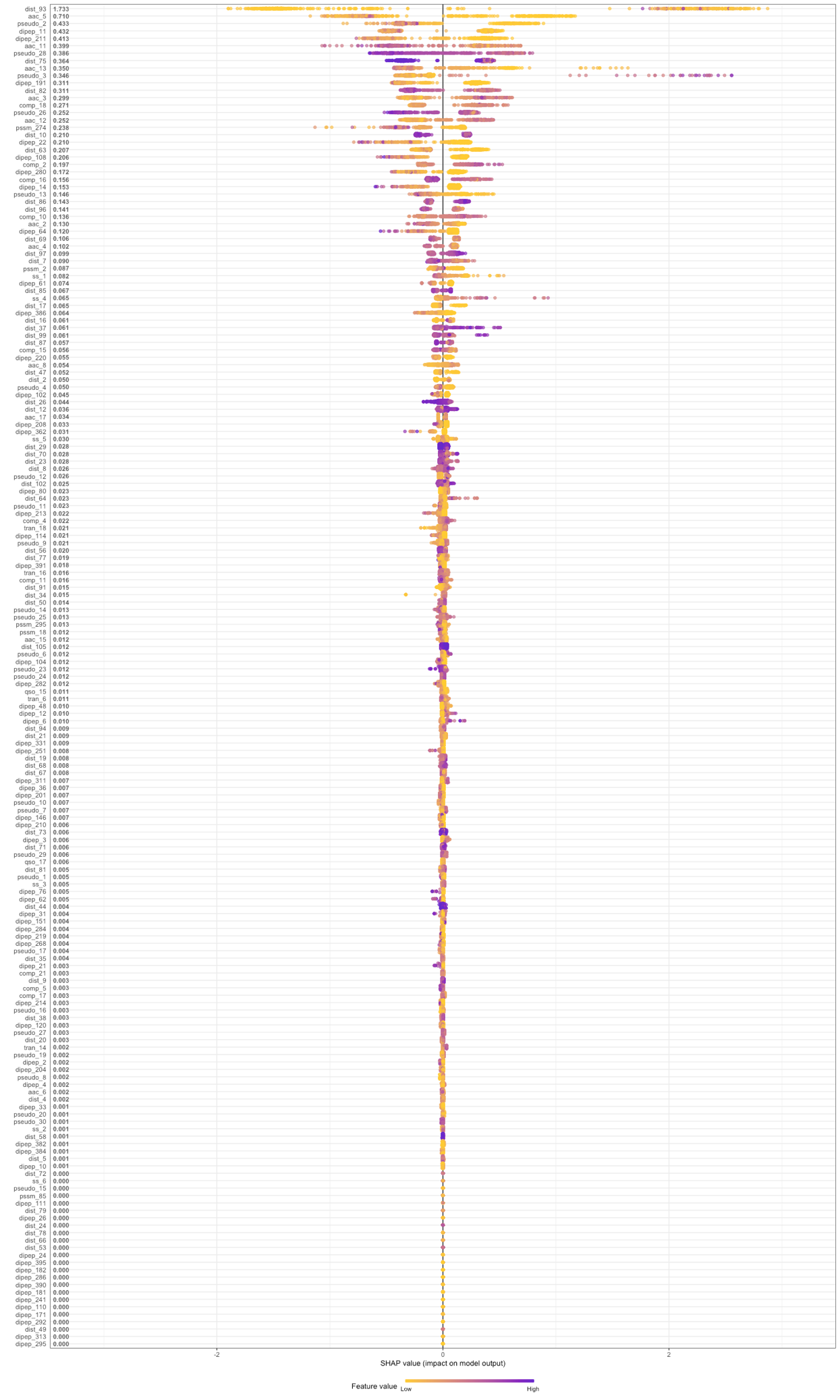
